# T cell-derived IFN-γ Suppresses T Follicular Helper Cell Differentiation and Antibody Responses

**DOI:** 10.1101/2024.12.31.630815

**Authors:** Eleonora Sala, Maria Nelli, Chiara Laura, Pietro Di Lucia, Cristian Gabriel Beccaria, Elisa B. Bono, Marta Mangione, Davide Marotta, Marta Grillo, Valentina Sperto, Leonardo Giustini, Fabio Tosi, Jia Nie, Daehong Kim, Giuliana Furiato, Chiara Malpighi, Eleonora Consolo, Burkhard Becher, Eyal David, Merav Cohen, Amir Giladi, Ido Amit, Remy Bosselut, Luca G. Guidotti, Matteo Iannacone, Mirela Kuka

## Abstract

CD4^+^ T cells play a critical role in antiviral humoral and cellular immune responses. We have previously reported that subcutaneous lymphocytic choriomeningitis virus (s.c. LCMV) infection is characterized by a stark compartmentalization of CD4^+^ T cells, leading to strong T_H_1 polarization but virtually absent T follicular helper (T_FH_) cells, a key driver of humoral immunity. Here, we investigated the mechanisms responsible for this impaired T_FH_ differentiation. We found that T-bet^+^ cells induced by s.c. LCMV infection encompass a T_H_1 subset expressing Granzyme-B (GzmB) and a Tcf-1^+^ subset that retains the potential for T_FH_ differentiation without expressing mature T_FH_ markers. Interestingly, IFN-γ blockade enables full differentiation of Tcf-1^+^ cells into T_FH_, formation of germinal centers and increased antibody production. Of note, the suppression of T_FH_ cells by IFN-γ is not directly mediated through CD4^+^ T cells but rather involves another cell type, likely dendritic cells (DCs). Our study provides novel insights into the mechanisms directing early CD4^+^ T cell polarization and affecting humoral responses to viruses, laying a foundation for the development of effective vaccine strategies.

## Introduction

CD4^+^ T cells play a crucial role in orchestrating adaptive immune responses against pathogens, guiding a complex array of signals and differentiation processes. Following their priming in secondary lymphoid organs, antigen-specific CD4^+^ T cells undergo both clonal expansion and differentiation into effector cells (Zhu *et al*., 2010). Throughout this process, these T cells are exposed to a diverse range of cytokines from infected cells, dendritic cells (DC), and stromal cells. In response, CD4^+^ T cells embark on specific differentiation pathways, leading to the formation of distinct T helper cell subsets (Mempel *et al*., 2004; Walsh and Mills, 2013; Eisenbarth, 2018; Tuzlak *et al*., 2021).

Infection by viruses or intracellular bacteria primarily leads to the generation of T_H_1 and T_FH_ cells (Sheikh and Groom, 2020; Kuka and Iannacone, 2021). T_H_1 cells, characterized by the expression of the master transcription factor T-bet and the production of high levels of IFN-γ, promote macrophage activation and bolster CD8^+^ T cell responses (Szabo *et al*., 2000; Schoenborn and Wilson, 2007; Sercan *et al*., 2010; Snell *et al*., 2016). Furthermore, autocrine IFN-γ plays a role in promoting the expansion and maintaining the T_H_1 phenotype (Bradley *et al*., 1996; Lighvani *et al*., 2001; Whitmire *et al*., 2005). Conversely, T_FH_ cells, which express Bcl-6 and CXCR5, migrate to B cell follicles, where they interact specifically with cognate B cells, facilitating germinal center reactions and the subsequent generation of high-affinity, class-switched antibodies (Crotty, 2011; Vinuesa *et al*., 2016).

Ideally, T_H_1 and T_FH_ subsets coexist, each contributing to adaptive immune responses by predominantly supporting cellular or humoral immunity, respectively. Based on literature, the bifurcation between T_FH_ and T_H_1 fates seems to take place within a few days upon CD4^+^ T cell activation (Choi *et al*., 2011; DiToro *et al*., 2018). However, previous work has shown some degree of overlap or competition between these two CD4^+^ T cell subsets (Nakayamada *et al*., 2011; Lönnberg *et al*., 2017). For example, it was reported that T_FH_ and T_H_1 share a transitional phase expressing both T-bet and Bcl-6. While the cells progress into reinforcing T_H_1 phenotype, T-bet suppresses further T_FH_ differentiation by competing with Bcl-6. This competition seems to be cell-intrinsic since CD4^+^ T cells lacking T-bet differentiate into T_FH_ (Nakayamada *et al*., 2011; Lönnberg *et al*., 2017). In another study, Bcl6-expressing T_FH_ cells generated upon viral infection expressed T-bet, which was critical for their development and function and transcriptionally required for proper T_FH_ cell programming (Weinstein *et al*., 2018). However, T-bet expression by T_FH_ has also been reported to render these cells dysfunctional, like in severe malaria infection (Obeng-Adjei *et al*., 2015; Ryg-Cornejo *et al*., 2016; Hansen *et al*., 2017). In this context, concomitant blockade of IFN-γ and TNF-α resulted in enhanced T_FH_ differentiation and improved antibody responses (Ryg-Cornejo *et al*., 2016). Thus, the interplay between T_H_1 and T_FH_ is marked by controversy and needs to be further dissected.

We have recently reported that subcutaneous (s.c.) infection with lymphocytic choriomeningitis virus (LCMV) results in almost exclusive T_H_1 differentiation and impaired T_FH_ induction (De Giovanni *et al*., 2020). While this observation is in line with the strong cellular responses and the weak neutralizing antibody (nAb) responses triggered by non-cytopathic viruses such as LCMV (Hangartner *et al*., 2006), the precise cellular and molecular mechanisms influencing this bias remain elusive.

In this study, we uncovered the heterogeneity within LCMV-induced T-bet^+^ cells, identifying two distinct subsets: a Tcf-1^+^ subset and a granzyme B (GzmB)^+^ subset. Surprisingly, neither subset required the canonical T_H_1-polarizing cytokine IL-12 for differentiation. Instead, IFN-γ emerged as a key regulator, driving the expansion of GzmB^+^ cells while suppressing the maturation of Tcf-1^+^ cells into fully differentiated T_FH_. Notably, blocking IFN-γ restored the T_FH_ population and enhanced germinal center B cells and Ab production. These findings reveal novel mechanisms that limit T_FH_ differentiation and Ab production during viral infections, providing valuable insights for developing innovative vaccination strategies.

## Results

### Heterogeneity of T-bet^+^ CD4^+^ T cells upon LCMV infection

In aiming to understand the determinants responsible for the impaired T_FH_ differentiation upon s.c. LCMV infection, our initial step was the comprehensive transcriptional profiling of antigen-specific CD4^+^ T cells. We utilized single-cell RNA sequencing (scRNA-seq) to analyze the diverse transcriptomic landscapes. To this end, naïve Smarta CD4^+^ T cells, which are reactive to the MHC-II-restricted GP61-80 epitope of the LCMV glycoprotein (Oxenius *et al*., 1998), were adoptively transferred into wild-type (WT) mice. This was done one day before the mice were subjected to s.c. (intrafootpad) infection with rLCMV (a recombinant LCMV clone 13 expressing the LCMV WE glycoprotein recognized by Smarta TCR-transgenic cells (Fallet *et al*., 2016; De Giovanni *et al*., 2020). Five days post-infection, a time point characterized by notable expansion of Ag-specific CD4^+^ T cells (De Giovanni *et al*., 2020), we performed the isolation and FACS-based sorting of Smarta CD4^+^ T cells from footpad-draining popliteal lymph nodes (dLNs) (**Fig. 1A**). Consistent with our previous report (De Giovanni *et al*., 2020), LCMV-specific CD4^+^ T cells generated in this setting predominantly expressed T-bet, a key transcriptional regulator of T_H_1 differentiation, while T_FH_ marker CXCR5 was notably absent (**Fig. S1**). For comparative analysis, Smarta CD4^+^ T cells from mice infected with rVSV, a recombinant VSV that expresses the LCMV WE glycoprotein and induces T_FH_ differentiation (Fallet *et al*., 2016; De Giovanni *et al*., 2020), served as a control (**Fig. S1**).

**Figure 1.**
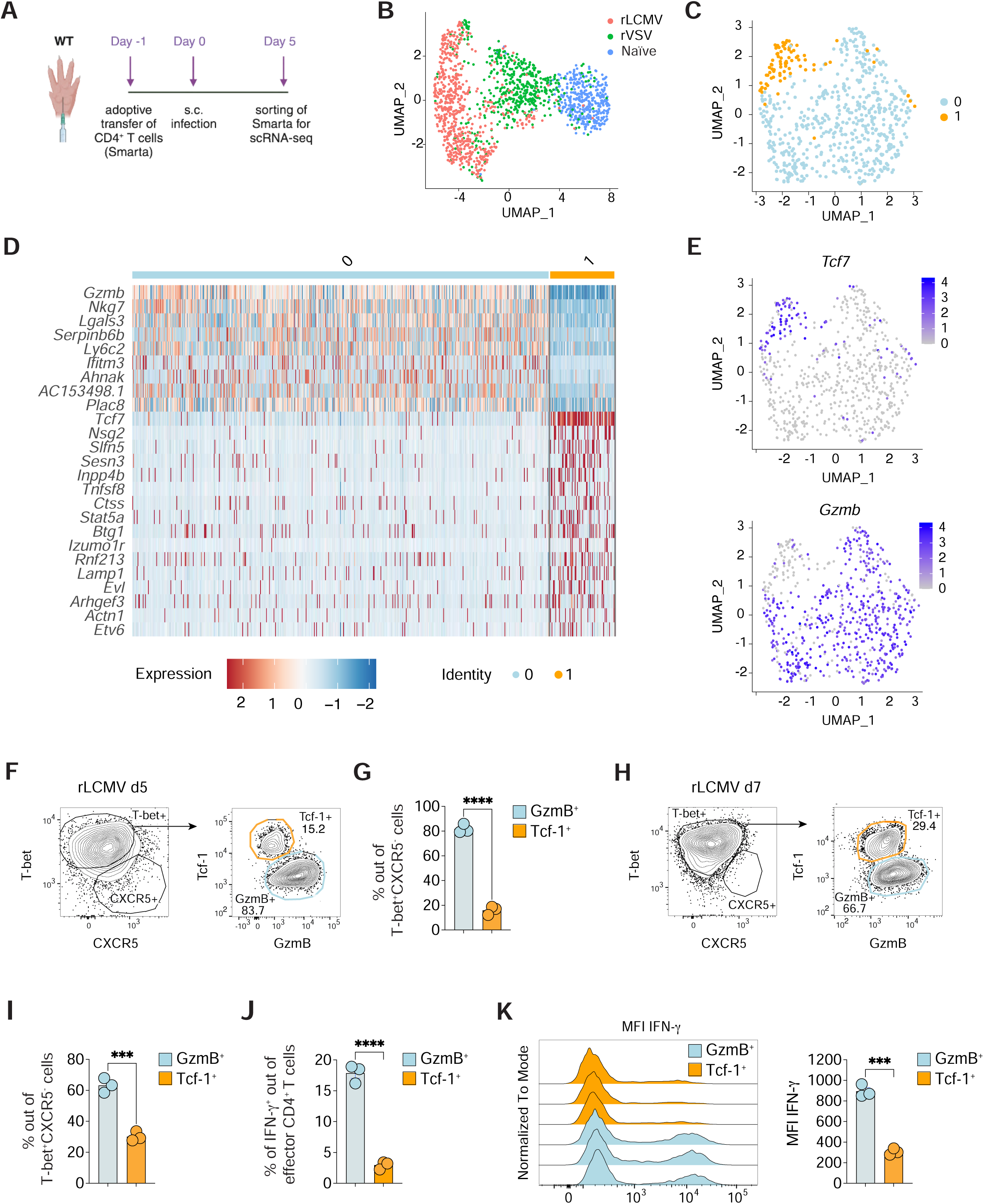
*Heterogeneity of T-bet^+^ CD4^+^ T cells upon LCMV infection.* **A)** Schematic representation of the experimental setup for the results described in B-E. 0.5*10^6^ purified Ag-specific CD45.1^+^ CD4^+^ T cells (Smarta) were transferred into CD45.2^+^ WT recipients one day before s.c. infection with either rLCMV or rVSV (1*10^5^ FFU or PFU /footpad). Smarta CD4^+^ T cells were FACS-sorted from the dLNs five days upon infection based on CD45.1 expression. **B)** UMAP projection of sorted cells. Each dot corresponds to a single cell, colored according to different samples. rLCMV cells (red, 643 cells), rVSV cells (green, 498 cells), naïve Smarta CD4^+^ T cells (light blue, 369 cells). **C)** UMAP projection of Smarta CD4^+^ T cells sorted from rLCMV infected mice. Each dot corresponds to a single cell, colored according to the unbiased clusters identified: cluster 0 (light blue, 557 cells) and cluster 1 (orange, 86 cells). **D)** Heatmap of normalized and scaled expression values of the top marker genes identifying the two clusters (logFC threshold: ±1and adjusted p-value < 0.05 filters were applied)). Color coding of the bar on the top of the heatmap as in (C). **E)** Feature plot representation of the expression level of the natural-log normalized expression level of *Gzmb* and *Tcf7* on the scRNA-seq subset described in (C). **F)** 0.5*10^6^ purified CD45.1^+^ Smarta CD4^+^ T cells were transferred into CD45.2^+^ WT recipients 1 day before s.c. rLCMV infection (1*10^5^ FFU /footpad). dLNs were analyzed 5 days post infection. Representative flow cytometry plots showing the frequencies of Tcf-1^+^ and GzmB^+^ cells among T-bet^+^CXCR5^-^ Smarta CD4^+^ T cells in the dLNs. Numbers represent the percentage of cells within the indicated gate. **G)** Quantification of GzmB^+^ and Tcf-1^+^ cells expressed as percentages of T-bet^+^CXCR5^-^Smarta CD4^+^ T cells in dLNs of mice described in F. *n=3*. Mean ± SEM is shown. Data are representative of at least five independent experiments. An unpaired two-tailed t test was applied. **** *p-value* <= 0.0001. **H)** CD45.2^+^ WT mice were infected s.c. with rLCMV (1*10^5^ FFU /footpad) and dLNs were analyzed 7 days post infection. Representative flow cytometry plots showing Tcf-1^+^ and GzmB^+^ cells among T-bet^+^CXCR5^-^ endogenous CD4^+^ T cells in the dLNs. Numbers represent the percentage of cells within the indicated gate. **I)** Quantification of GzmB^+^ and Tcf-1^+^ cells expressed as percentages of T-bet^+^CXCR5^-^ endogenous CD4^+^ T in dLNs. *n=3*. Mean ± SEM is shown. Data are representative of three independent experiments. An unpaired two-tailed t test was applied: *** *p-value* <= 0.001. **J**) Quantification of IFN-γ^+^ cells among the GzmB^+^ and Tcf-1^+^ subsets out of effector CD44^+^ CD62L^lo^ CD4^+^ T cells in dLNs of mice described in H. *n=3.* Mean ± SEM is shown. Data are representative of three independent experiments. An unpaired two-tailed t test was applied: **** *p-value* <= 0.0001. **K)** Representative flow cytometry plot (left panel) and quantification (right panel) showing the fluorescence intensity of IFN-γ expression within GzmB^+^ or Tcf-1^+^ CD4^+^ T cells of mice described in H. *n=3.* Mean ± SEM is shown. Data are representative of three independent experiments. An unpaired two-tailed t test was applied: *** *p-value* <= 0.001.

We then performed massively parallel single-cell RNA-sequencing (MARS-seq) on QC-positive single T cells. Using the Seurat R package (Stuart *et al*., 2019), we scrutinized the scRNA-seq dataset, revealing distinct cellular clusters through uniform manifold approximation and projection (UMAP) analysis (McInnes *et al*., 2018). It was evident that control Smarta CD4^+^ T cells isolated from naïve mice (*n* = 369, marked in blue) were characterized by a higher expression of canonical naïve T cell markers such as *Ccr7, Sell,* and *Klf2 (***Fig. S2A**). In stark contrast, the CD4^+^ T cells from the infected hosts were discretely clustered based on the infective agent, confirming the considerable divergence between the CD4^+^ T cell subsets post VSV or LCMV infection (**Fig. 1B and S2B and Table 1**) (De Giovanni *et al*., 2020).

Focusing on the rLCMV condition, we discerned two distinct clusters within the Smarta CD4^+^ T cells (**Fig. 1C**). The dominant cluster 0 (*n* = 557) showed upregulated expression of genes such as *Gzmb, Nkg7,* and *Ly6c2* (**Fig. 1D** and **E and Table 2**). Conversely, cluster 1 (*n* = 86) was characterized by elevated levels of *Tcf7*, a transcription factor implicated in T_FH_ differentiation (Xu *et al*., 2015) (**Fig. 1D** and **E and Table 2**). Intriguingly, cluster 1 overlaid with cells at the rLCMV and rVSV interface when backgated onto the original UMAP, hinting at a transitional phenotype (**Fig. S2C)**. In addition, a GSEA analysis showed that the rVSV signature was significantly enriched in cluster 1 but not in cluster 0 (**Fig. S2D)**, indicating that cluster 1 is enriched in cells with a transcriptional signature similar to the T_FH_ in VSV infection.

Flow cytometry analyses were performed to corroborate the scRNA-seq findings, revealing two distinct populations within the T-bet^+^ CD4^+^ Smarta T cells five days after rLCMV infection. These were classified based on the expression of GzmB and Tcf-1 (encoded by *Tcf7*) proteins, paralleling the scRNA-seq identified subsets (**Fig. 1F,G and S3A-C**). The GzmB^+^ population was predominant, while the Tcf-1^+^ subset constituted a smaller, yet significant fraction. Further examination of the GzmB^+^ cells showed elevated CXCR6 and Ly6C expression, along with marginally increased T-bet levels compared to the Tcf-1^+^ group (**Fig. S3D**). The Tcf-1^+^ cells instead showed slightly higher levels of Bcl-6, CXCR5, and CXCR3 (**Fig. S3D**). Also the endogenous CD4^+^ T cell compartment seven days post rLCMV infection contained these two subsets, with comparable frequencies to the Smarta cells (**Fig. 1H,I and S4A,B**). Functionally, the GzmB^+^ subset distinguished itself with a robust IFN-γ response upon GP61 peptide re-stimulation, a trait not as pronounced in the Tcf-1^+^ cells (**Fig. 1J**). The differential IFN-γ expression was further quantified, revealing significantly higher mean fluorescence intensity (MFI) in GzmB^+^ cells compared to their Tcf-1^+^ counterparts (**Fig. 1K**).

These findings underscore the existence of two functionally and phenotypically distinct subsets within the T-bet^+^ CD4^+^ T cell population upon s.c. rLCMV infection: the GzmB^+^ population probably representing the classical IFN-γ-producing T_H_1 cells, and the Tcf-1^+^ population as a putative precursor of *bona fide* T_FH_.

### IFN-γ suppresses T follicular helper differentiation and antibody production

Next, we sought to elucidate the molecular determinants driving the T_H_1 polarization and heterogeneity induced by LCMV infection. We initially investigated the role of IL-12, a well-known T_H_1-polarizing cytokine (Heufler *et al*., 1996; Athie-Morales *et al*., 2004). In line with previous studies on viral infections (Schijns *et al*., 1998; Oxenius *et al*., 1999; Krueger *et al*., 2021), neutralization of IL-12 did not significantly alter CD4^+^ T cell differentiation (**Fig. S5A-D**), prompting us to explore the influence of another cytokine known for reinforcing T_H_1 identity through positive feedback loops, IFN-γ (Wakil *et al*., 1998; Lighvani *et al*., 2001; Schulz *et al*., 2009).

We examined the effects of IFN-γ by transferring Smarta CD4^+^ T cells into wild-type mice treated with IFN-γ blocking antibodies (**Fig. 2A**). IFN-γ blockade affected the expansion of Smarta CD4^+^ T (**Fig. S6A**), in line with the previously reported role for IFN-γ in expansion of antiviral CD4^+^ T cells (Whitmire *et al*., 2005). Intriguingly, blocking IFN-γ shifted the balance in favor of the Tcf-1^+^ subset at the expense of the GzmB^+^ population, both in terms of percentages and absolute numbers (**Fig. 2B and S6B**). This was accompanied by an upregulation of the canonical T_FH_ marker CXCR5 in the Tcf-1^+^ subset (**Fig. 2C** and **D**). In addition, Tcf-1^+^ cells showed lower levels of T-bet and higher levels of PD-1, Bcl-6, CD95 and CXCR3 when IFN-γ was blocked (**Fig. S6C**). As a result, IFN-γ blockade resulted in a decrease in T-bet^+^ T_H_1 cells while promoting the emergence of CXCR5^+^ T_FH_ cells (**Fig. 2E, F, and S6D**). Notably, the effect of IFN-γ blockade in CD4^+^ T cell polarization was not a result of higher viral replication, since viral titers in mice receiving the α-IFN-γ neutralizing Ab were lower at day 3 and similar to control mice at day 5 after LCMV infection (**Fig. S6E**). In addition, we have previously shown that CD4^+^ T cells differentiate into T_H_1 but not T_FH_ even when animals are infected with 100-fold higher or lower viral loads (De Giovanni *et al*., 2020).

**Figure 2.**
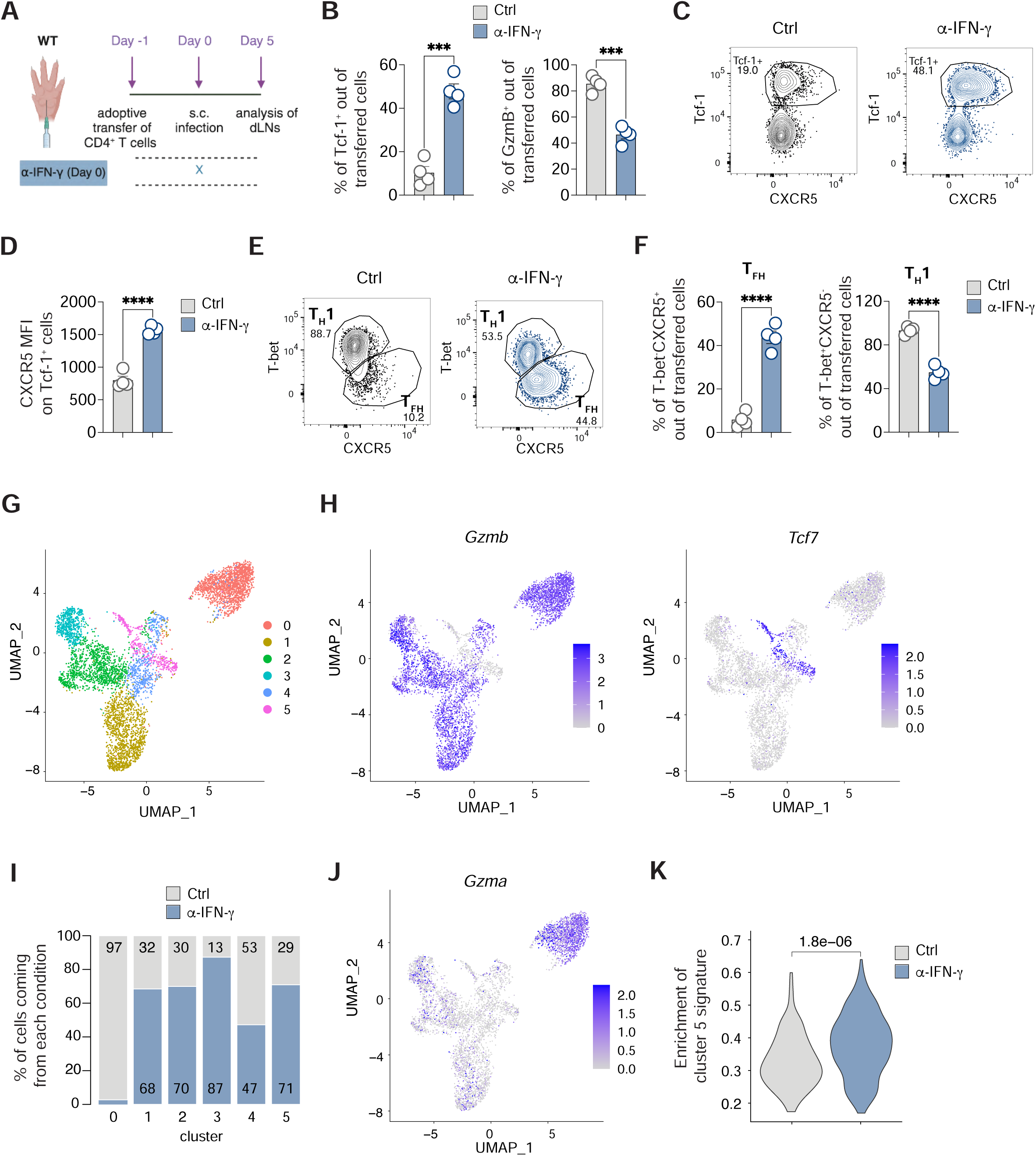
*IFN-γ suppresses T follicular helper cell differentiation.* **A)** 0.5*10^6^ purified CD45.1^+^ Smarta CD4^+^ T cells were transferred into CD45.2^+^ WT recipients 1 day before s.c. rLCMV infection (1*10^5^ FFU /footpad). CD45.2^+^ WT recipient mice were also treated with α-IFN-γ blocking antibody (or isotype Ctrl) at day 0. dLNs were analyzed 5 days post infection. **B)** Quantification of Tcf-1^+^ (left) and GzmB^+^ (right) in dLNs of mice described in (A) expressed as percentages out of transferred Smarta CD4^+^ T cells in. *n=4*. Mean ± SEM is shown. Data are representative of at least three independent experiments. An unpaired two-tailed t test was applied. \*\*\**p-value* <= 0.001. **C)** Representative flow cytometry plots showing CXCR5 expression on Tcf-1^+^ and Tcf-1^-^ Smarta CD4^+^ T cells in dLNs of mice described in (A). Numbers represent the percentage of cells within the indicated gate. **D)** Quantification of the MFI of CXCR5 on Tcf-1^+^ Smarta CD4^+^ T cells in dLNs of mice described in (A). *n=4*. Mean ± SEM is shown. Data are representative of at least three independent experiments. An unpaired two-tailed t test was applied. \*\*\*\**p-value* <= 0.0001. **E)** Representative flow cytometry plots showing T_H_1 (T-bet^+^CXCR5^-^) and T_FH_ (T-bet^-^CXCR5^+^) cells among Smarta CD4^+^ T cells in dLNs of mice described in (A). Numbers represent the percentage of cells within the indicated gate. **F)** Quantification of T_FH_ (left) and T_H_1 (right), expressed as percentages out of transferred Smarta CD4^+^ T cells in dLNs of mice described in (A). *n=4*. Mean ± SEM is shown. Data are representative of at least three independent experiments. An unpaired two-tailed t test was applied. \*\*\*\**p-value* <= 0.0001. **G)** UMAP projection of 5,746 sorted and sequenced LCMV-specific CD4^+^ T cells. Each dot corresponds to a single cell, colored according to unbiased clusters identified using the Louvain algorithm. **H)** Feature plot representation of the natural-log normalized expression level of *Gzmb* (left panel) and *Tcf7* (right panel) on the dataset in (G). **I)** Barplot representation of the frequencies of the two conditions (control in grey and α-IFN-γ in blue) in each cluster. **J)** Feature plot representation of the natural-log normalized expression level of *Gzma* on the dataset in (G). **K)** Violin plot showing the enrichment of a signature composed by marker genes of cluster 5 versus all the other clusters, comparing control (grey) and α-IFN-γ (blue) cells in cluster 5 only. Two-tailed Mann-Whitney test has been performed and the resulting p-value is shown.

To delve deeper into the transcriptional changes induced by IFN-γ, we conducted single-cell RNA sequencing (scRNA-seq) on Smarta cells from mice treated with either PBS or anti-IFN-γ antibody. This analysis unveiled six distinct clusters within LCMV-specific CD4^+^ T cells (**Fig. 2G** and **S7A and Table 3**). Among these, five expressed *Gzmb*, while one cluster (cluster 5) was *Gzmb*-negative but displayed high levels of *Tcf7* (**Fig. 2H and Table 3**). *Gzmb*^+^ clusters showed lower *Cxcr5* expression and higher *Tbx21* (which encodes for T-bet) levels (**Fig. S7B**), consistent with previous protein expression findings (**Fig. S3D**). These *Gzmb*^+^ cells also exhibited slightly higher expression of *Ifngr1*, suggesting heightened sensitivity to IFN-γ (**Fig. S7C**). Notably, cluster 0, characterized by *Gzma* expression, was almost exclusively present in PBS-treated animals, indicating a strong dependence on IFN-γ for its development (**Fig. 2I** and **S7D**).

Further analysis of the *Tcf7*-expressing cluster revealed a significant enrichment of cells from the anti-IFN-γ-treated group, constituting 71% of the cluster, compared to 29% from PBS-treated mice. This confirms our observation of Tcf-1^+^ cell increase following IFN-γ blockade (**Fig. 2I** and **S6B**). Additionally, Tcf7-expressing cells in the anti-IFN-γ group displayed markedly higher levels of the gene signature characteristic of cluster 5 compared to all other clusters (**Fig. 2K**). This finding suggests that the T_FH_ precursor gene signature in LCMV-infected mice is amplified in the absence of IFN-γ.

All in all, our data reveal substantial heterogeneity among T-bet^+^ cells in the context of LCMV infection. Specifically, one cluster’s development relies exclusively on IFN-γ, while the *Tcf7*-expressing cluster is numerically and phenotypically suppressed by IFN-γ, highlighting the complexity of CD4^+^ T cell differentiation in this scenario.

We further investigated whether Tcf-1^+^ CXCR5^+^ cells, generated in the absence of IFN-γ, functionally resemble T_FH_ cells. Confocal microscopy of dLNs from rLCMV-infected mice confirmed a marked increase in Tcf-1^+^ antigen-specific CD4^+^ T cells (Smarta) without IFN-γ (**Fig. 3A** and **B**). Notably, these Tcf-1^+^ cells were located not only in interfollicular regions but also within B cell follicles, indicating their transition to fully differentiated and functional CXCR5^+^ T_FH_ cells (**Fig. 3A**-**C**). This observation strongly suggests that IFN-γ inhibits T_FH_ cell differentiation by preventing the maturation of Tcf-1^+^ cells into CXCR5^+^ T_FH_ cells.

**Figure 3.**
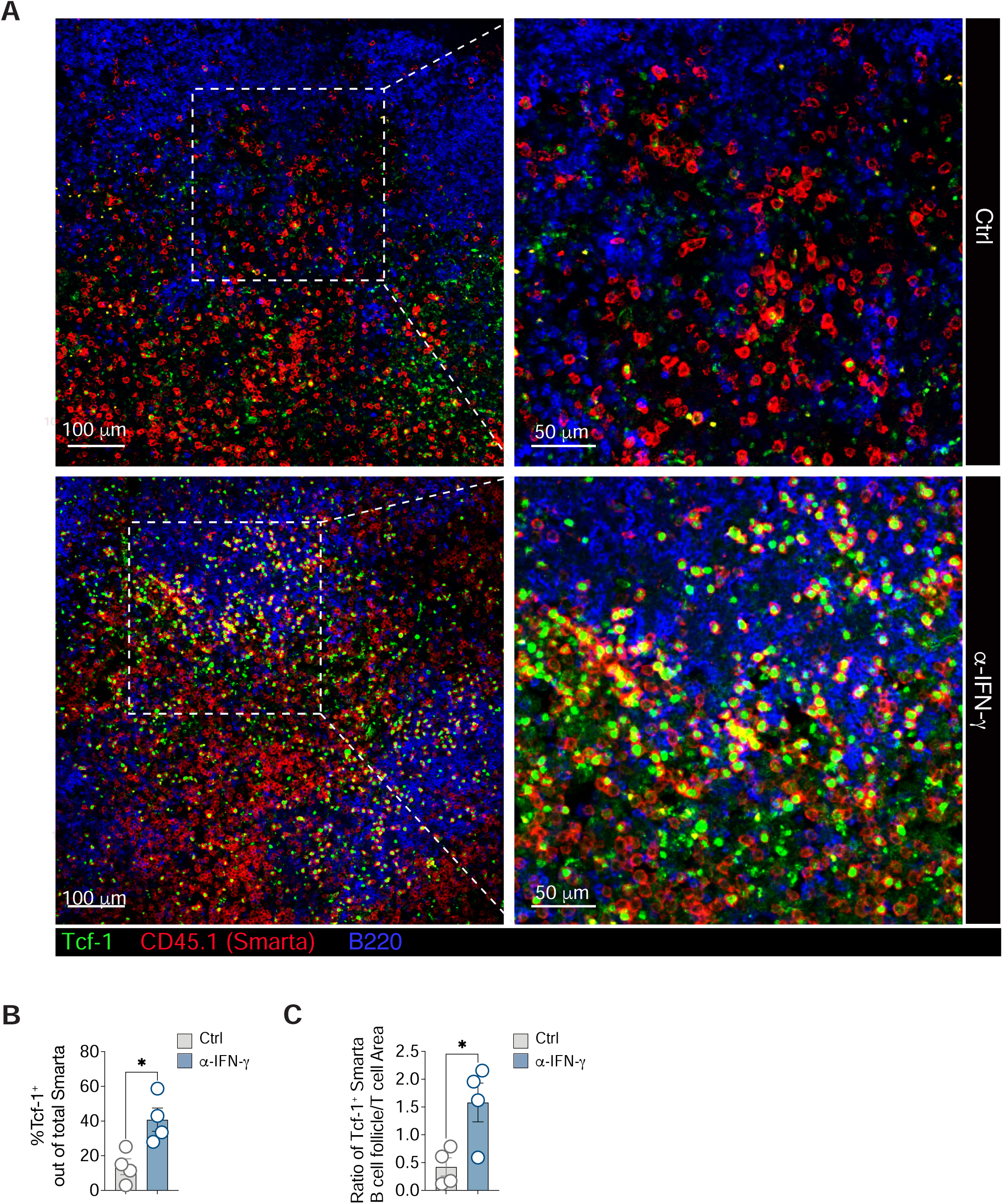
*IFN-γ suppresses T follicular helper cell responses upon viral infection.* **A)** 0.5*10^6^ purified CD45.1^+^ Smarta CD4^+^ T cells were transferred into CD45.2^+^ WT recipients one day before s.c. rLCMV infection (1*10^5^ FFU /footpad). CD45.2^+^ WT recipient mice were also treated with α-IFN-γ blocking antibody (or isotype Ctrl) at day 0. dLNs were analyzed by confocal microscopy 5 days post infection. CD45.1^+^ (Smarta CD4^+^ T cells) are depicted in red, Tcf-1^+^ cells in green and B220^+^ cells (B cell follicles) in blue. Images on the right panels are a magnification of the dotted square areas on the left. Data are representative of two independent experiments. Scale bars represent 100 (left) and 50 (right) μm. **B)** Quantification of Tcf-1^+^ cells out of total Smarta CD4^+^ T cells in the images (left panels) in (A). *n=4.* Mean ± SEM is shown. Data are representative of two independent experiments. An unpaired two-tailed t test was applied. * *p-value* <= 0.05. **C)** Calculation of the ratio of the density of Tcf-1^+^Smarta^+^ in the B cell follicle versus the T cell area in the images (left panels) in (A). *n=4.* Mean ± SEM is shown. Data are representative of two independent experiments. An unpaired two-tailed t test was applied. * *p-value* <= 0.05.

Interestingly, IFN-γ blockade resulted in a notable rescue of T_FH_ cells even in an endogenous setting where T cells were analyzed seven days upon s.c. infection (**Fig. 4A-B**). These T_FH_ generated upon IFN-γ blockade expressed Bcl-6 hinting to their complete differentiation (**Fig. 4C-D**). Moreover, IFN-γ was found to significantly impact B cell activation (**Fig. 4E** and **F**). Approximately 15-20% of B cells in the dLNs of rLCMV-infected mice expressed high levels of T-bet seven days post-infection, compared to a meager 1% expressing Bcl-6. Blocking IFN-γ, however, led to an increase in Bcl-6^+^ B cells and a decrease in T-bet^+^ B cells (**Fig. 4E** and **F**). This shift in B cell populations was in line with higher levels of total LCMV GP-binding IgG antibodies in the absence of IFN-γ (**Fig. 4G**). Collectively, these findings underscore the inhibitory role of IFN-γ in the development of T_FH_ cells, germinal center B cells, and antibody responses during viral infection.

**Figure 4.**
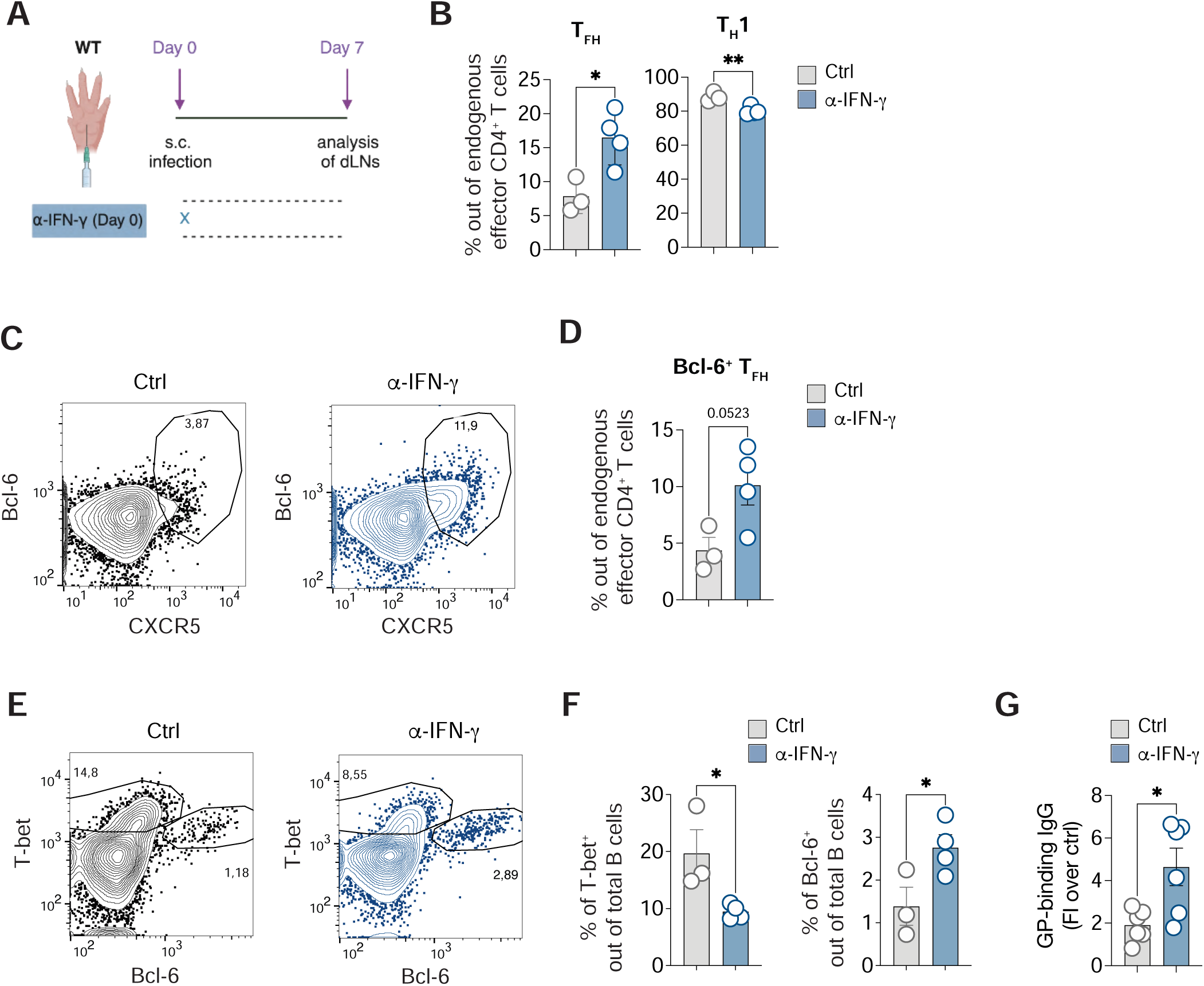
*IFN-γ suppresses T follicular helper cell and B cell responses in an endogenous setting.* **A)** CD45.2^+^ WT mice were infected s.c. with rLCMV (1*10^5^ FFU /footpad) and treated with α-IFN-γ blocking antibody (or isotype Ctrl) at day 0. dLNs were analyzed 7 days post infection. **B)** Quantification of T_FH_ (left) and T_H_1 (right), expressed as percentages out of endogenous effector CD4^+^ T cells in dLNs of s.c. infected mice described in (A). *n*=3-4. Mean ± SEM is shown. Data are representative of three independent experiments. An unpaired two-tailed t test was applied. * *p-value* <= 0.05, ** *p-value* <= 0.01. **C)** Representative flow cytometry plots showing CXCR5^+^Bcl-6^+^ cells expressed as percentages among endogenous effector CD4^+^ T cells in the dLNs. Numbers represent the percentage of cells within the indicated gate. **D)** Quantification of Bcl-6^+^ T_FH_, expressed as percentages out of endogenous effector CD4^+^ T cells in dLNs of s.c. infected mice described in (A). *n*=3-4. Mean ± SEM is shown. Data are representative of three independent experiments. An unpaired two-tailed t test was applied. **E)** Representative flow cytometry plots showing T-bet^+^ and Bcl-6^+^ B cells expressed as percentages among total B cells (B220^+^) in the dLNs. Numbers represent the percentage of cells within the indicated gate. **F)** Quantification of T-bet^+^ cells (left) and Bcl-6^+^ cells (right) expressed as percentages out of total B cells in dLNs of mice described in (A). *n=3* (Ctrl), 4 (α-IFN-γ). Mean ± SEM is shown. Data are representative of three independent experiments. An unpaired two-tailed t test was applied. * *p-value* <= 0.05. **G)** LCMV GP–binding IgG Abs were measured in the sera of mice 7 days upon infection as described in (A) and expressed as fold induction over uninfected controls. *n*=6. Mean ± SEM is shown. Data were pooled from two independent experiments. An unpaired two-tailed t test was applied. * *p-value* <= 0.05.

### Timing of IFN-γ-mediated suppression of T_FH_ differentiation

To elucidate the molecular mechanisms and timing of IFN-γ-mediated suppression of T_FH_ differentiation, we explored the effects of IFN-γ blockade initiated at different stages post-infection (**Fig. 5A**). Blocking IFN-γ at day 3 post-infection did not significantly alter the frequencies of Tcf-1^+^ and GzmB^+^ populations (**Fig. 5B**). Moreover, while IFN-γ inhibition at day 0 significantly increased CXCR5 expression on Tcf-1^+^ cells, initiating blockade at day 3 produced only minimal, non-significant changes (**Fig. 5C and S8A**). Consequently, an increase in functional T_FH_ frequencies was observed only when IFN-γ blockade was implemented from the onset of infection (**Fig. 5D**). This suggests that IFN-γ produced in the initial days post-infection is crucial for impeding T_FH_ differentiation.

**Figure 5.**
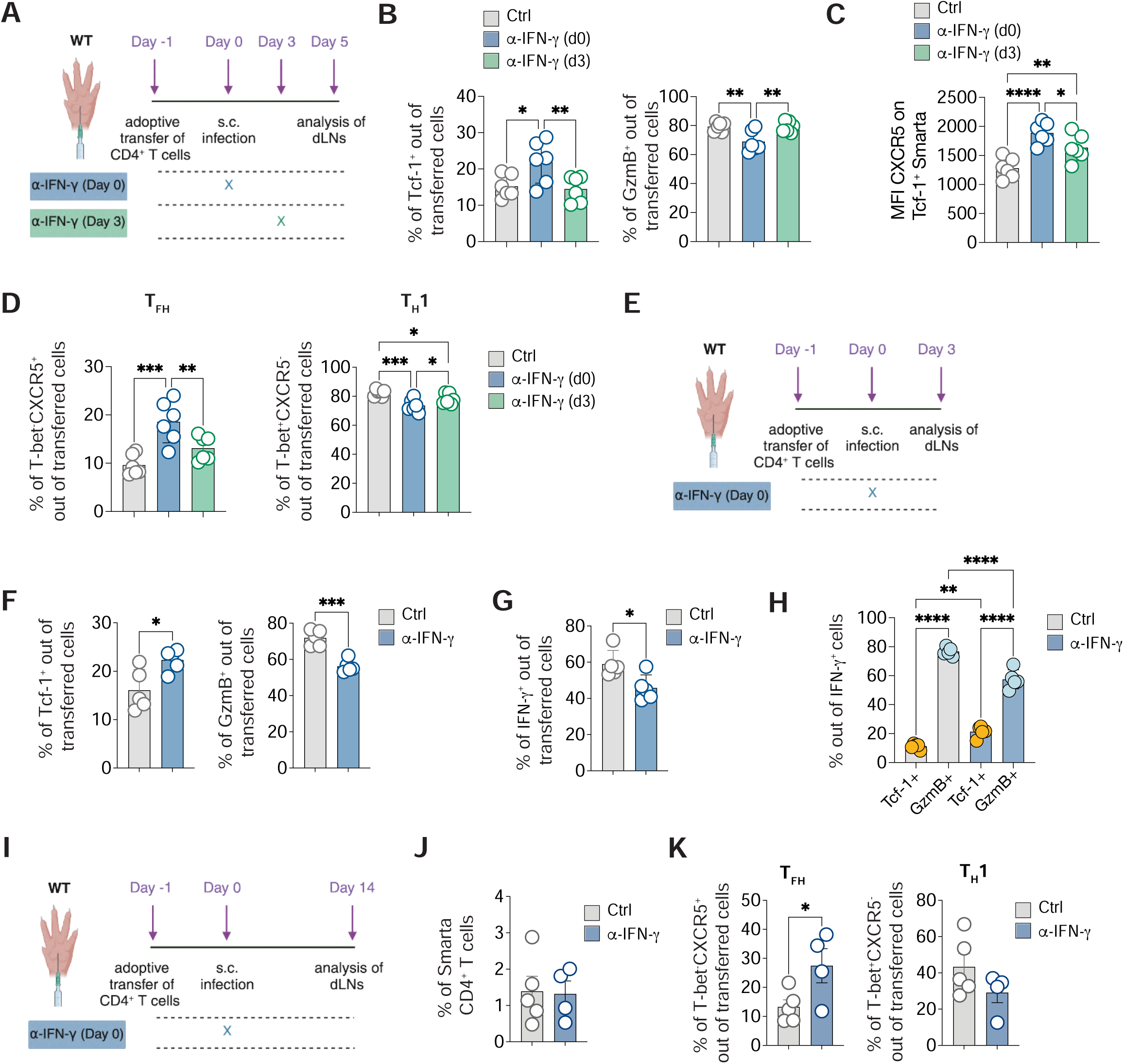
*The IFN-γ responsible for T_FH_ suppression is produced in the first days upon infection.* **A)** 0.5*10^6^ purified CD45.1^+^ Smarta CD4^+^ T cells were transferred into CD45.2^+^ WT recipients 1 day before s.c. rLCMV infection (1*10^5^ FFU /footpad). CD45.2^+^ WT recipient mice were also treated with α-IFN-γ blocking antibody (or isotype Ctrl) at day 0 or d3 after infection. dLNs were analyzed 5 days post infection. **B**) Quantification of Tcf-1^+^ (left) and GzmB^+^ (right), expressed as percentages out of transferred Smarta CD4^+^ T in dLNs of mice described in (A). *n=6*. Mean ± SEM is shown. Data are representative of three independent experiments. One-way ANOVA with uncorrected Fisher’s LSD was applied. * *p-value* <= 0.05, ** *p-value* <= 0.01. **C)** Quantification of the MFI of CXCR5 on Tcf-1^+^ Smarta CD4^+^ T cells in dLNs of mice described in (A). *n=6*. Mean ± SEM is shown. Data are representative of three independent experiments. One-way ANOVA with uncorrected Fisher’s LSD was applied. * *p-value* <= 0.05, ** *p-value* <= 0.01, **** *p-value* <= 0.0001. **D**) Quantification of T_FH_ (left) and T_H_1 (right), expressed as percentages out of transferred Smarta CD4^+^ T cells in dLNs of mice described in (A). *n=6*. Mean ± SEM is shown. Data are representative of three independent experiments. One-way ANOVA with uncorrected Fisher’s LSD was applied. * *p-value* <= 0.05, ** *p-value* <= 0.01, *** *p-value* <= 0.001. **E)** 0.5*10^6^ purified CD45.1^+^ Smarta CD4^+^ T cells were transferred into CD45.2^+^ WT recipients 1 day before s.c. rLCMV infection (1*10^5^ FFU /footpad). CD45.2^+^ WT recipient mice were also treated with α-IFN-γ blocking antibody (or isotype Ctrl) at day 0. dLNs were analyzed 3 days post infection. **F)** Quantification of Tcf-1^+^ and GzmB^+^ cells, expressed as percentages out of transferred Smarta CD4^+^ T cells in dLNs of mice described in (E). *n=5*. Mean ± SEM is shown. Data are representative of two independent experiments. An unpaired two-tailed t test was applied. * *p-value* <= 0.05, *** *p-value* <= 0.001. **G)** Quantification of IFN-γ^+^ cells out of ex-vivo restimulated Smarta CD4^+^ T cells in dLNs of mice described in (E). *n=5*. Mean ± SEM is shown. Data are representative of two independent experiments. An unpaired two-tailed t test was applied. * *p-value* <= 0.05. **H)** Quantification of GzmB^+^ and Tcf-1^+^ cells expressed as percentages among IFN-γ^+^ Smarta CD4^+^ T cells in dLNs of mice described in (E). *n=5*. Mean ± SEM is shown. Data are representative of two independent experiments. One-way ANOVA with uncorrected Fisher’s LSD was applied. ** *p-value* <= 0.01, **** *p-value* <= 0.0001. **I)** 0.5*10^6^ purified CD45.1^+^ Smarta CD4^+^ T cells were transferred into CD45.2^+^ WT recipients 1 day before s.c. rLCMV infection (1*10^5^ FFU /footpad). CD45.2^+^ WT recipient mice were also treated with α-IFN-γ blocking antibody (or isotype Ctrl) at day 0. dLNs were analyzed 14 days post infection. **J)** Quantification of Smarta CD4^+^ T cells, expressed as percentages out of total LNs cells of mice described in (D). **J**) Quantification of T_FH_ (left) and T_H_1 (right), expressed as percentages out of transferred Smarta CD4^+^ T cells in dLNs of mice described in (D). *n=5* (Ctrl), *4* (α-IFN-γ). Mean ± SEM is shown. Data are representative of two independent experiments. An unpaired two-tailed t test was applied. \**p-value* <= 0.05.

Further analyses were conducted to ascertain if CD4^+^ T cell polarization influenced by IFN-γ could be observed earlier than day 5 post-infection. Upon examining the LCMV-specific CD4^+^ T cells at day 3 post s.c. infection, we noted an increase in the Tcf-1^+^ population coupled with a decrease in GzmB^+^ cells when IFN-γ was blocked (**Fig. 5E, F, and S8B**). These observations indicate that IFN-γ begins shaping CD4^+^ T cell differentiation early during infection, influencing both T_FH_ and T_H_1 precursor populations. Interestingly, at this early stage, LCMV-specific CD4^+^ T cells were already capable of producing IFN-γ upon antigen re-stimulation, indicating their functional maturity just three days post-infection (**Fig. 5G**). The production of IFN-γ was reduced in mice treated with the IFN-γ-blocking antibody, aligning with its role in stabilizing the T_H_1 phenotype (**Fig. 5G and S8C**). Although both T cell subsets could produce IFN-γ, most of the cytokine was derived from GzmB^+^ cells (**Fig. 5H**).

We next asked for how long could differences in T cell polarization upon IFN-γ blockade be detected. To this end, Smarta CD4^+^ T cells were analyzed 14 days upon infection (**Fig. 5I**). The frequency of Smarta cells at this time-point was highly variable and significantly lower than day five, due to the contraction phase all T cells undergo after viral clearance (Fig. **5J**). Nonetheless, IFN-γ blockade still resulted in increased T_FH_ frequencies (Fig. **5K**) indicating that a single-dose of blocking antibody at the beginning of infection was sufficient to affect T cell polarization for weeks.

Overall, our findings demonstrate that the presence of IFN-γ during early priming events crucially influences CD4^+^ T cell differentiation, favoring T_H_1 cell development at the expense of T_FH_ cells.

### IFN-γ Produced by T Cells Is Key in T_FH_ Suppression

To investigate which might be the cellular source of the early IFN-γ crucial for T_FH_ suppression, we utilized the IFN-γ-Yellow Fluorescent Protein (YFP) mouse reporter model (Reinhardt *et al*., 2009) to identify cells expressing this cytokine during the initial stages of LCMV infection. Analysis of IFN-γ-YFP mice revealed diverse immune cell types (identified through gating strategy in **Figure S9**) expressing IFN-γ as early as 24 hours post-infection, including NK1.1^+^ group 1 innate lymphoid cells (ILCs) (comprising both NK cells and ILC1s), T lymphocytes, monocytes, and DCs (**Fig. 6A**, left panel). By 48 hours, group 1 ILCs emerged as the predominant IFN-γ-expressing subset, followed by CD8^+^ T cells (**Fig. 6A**, middle panel). Three days post-infection, group 1 ILCs, CD8^+^ T cells, CD4^+^ T cells, and monocytes were the main IFN-γ producers (**Fig. 6A**, right panel).

**Figure 6.**
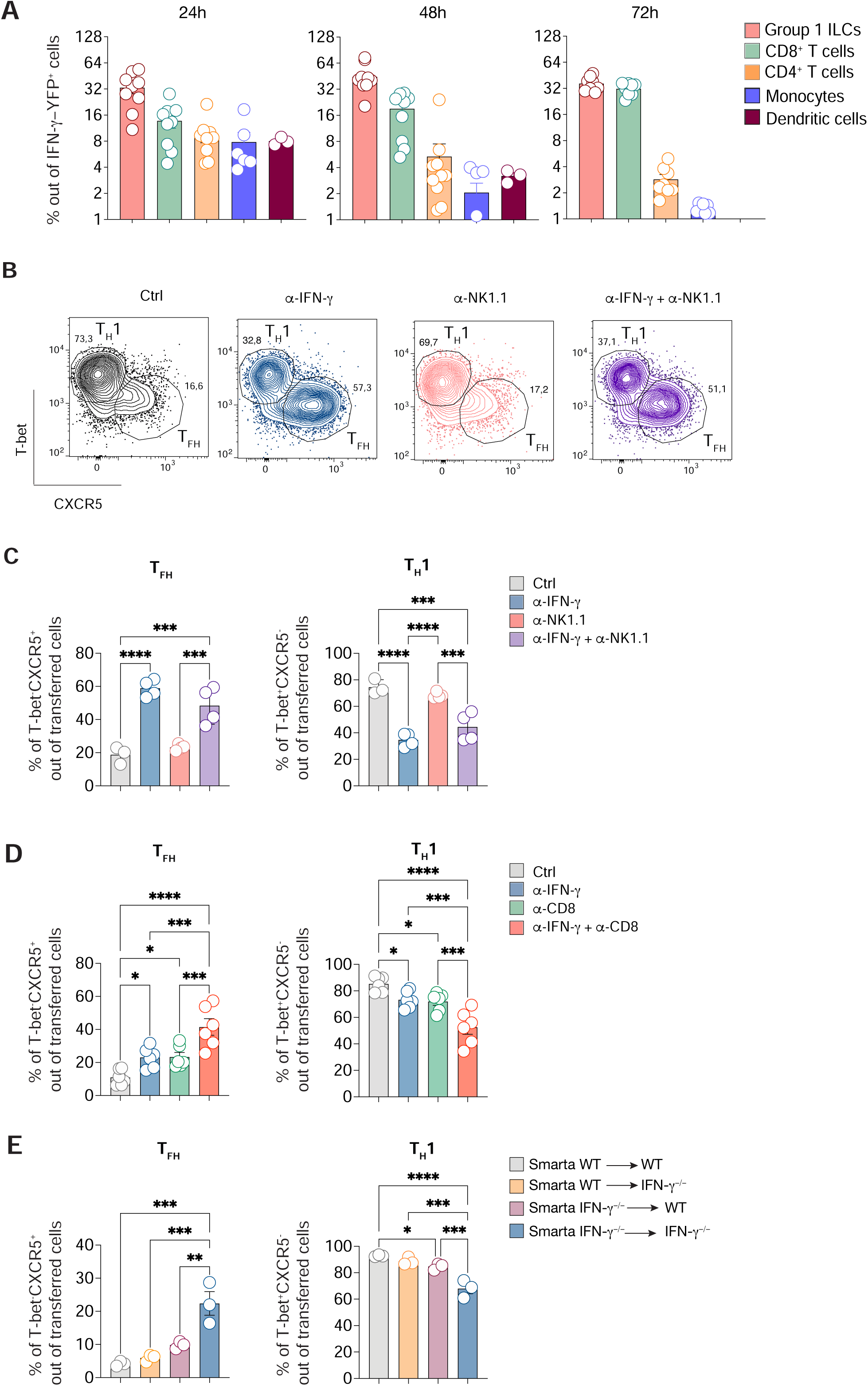
*IFN-γ produced by T cells is responsible for T_FH_ suppression **A**)* dLNs of IFN-γ-YFP mice were analyzed at 24h, 48h and 72 hours upon s.c. rLCMV infection (1*10^5^ FFU /footpad). Quantification of Group 1 ILCs (NK1.1^+^ NKp46^+^), CD8^+^, CD4^+^, B cells, monocytes (CD11b^+^ Ly6C^hi^), DC (CD11c^+^ MHC-II^+^) expressed as percentages out of the total YFP^+^ cells is shown. *n=3-9* (24, 48h), 7 (72h). Mean ± SEM is shown. Data are representative of two independent experiments. **B)** 0.5*10^6^ purified CD45.1^+^ Smarta CD4^+^ T cells were transferred into CD45.2^+^ WT recipients 1 day before s.c. rLCMV infection (1*10^5^ FFU /footpad). In some conditions CD45.2^+^ WT recipient mice were also treated with α-IFN-γ blocking antibody at day 0, α-NK1.1 antibody (d-1, d0) or both antibodies in combination. dLNs were analyzed 5 days post infection. Representative flow cytometry plot showing T_FH_ and T_H_1 cells among Smarta CD4^+^ T cells in dLNs. Numbers represent the percentage of cells within the indicated gate. **C)** Quantification of T_FH_ and T_H_1 cells, expressed as percentages out of transferred Smarta CD4^+^ T cells, in dLNs of mice described in (B). *n=3* (Ctrl), 4 (α-IFN-γ, α-NK1.1, α-IFN-γ + α-NK1.1). Mean ± SEM is shown. Data are representative of three independent experiments. One-way ANOVA with uncorrected Fisher’s LSD was applied. *** *p-value* <= 0.001, **** *p-value* <= 0.0001. **D)** 0.5*10^6^ purified CD45.1^+^ Smarta CD4^+^ T cells were transferred into CD45.2^+^ WT recipients 1 day before s.c. rLCMV infection (1*10^5^ FFU /footpad). In some conditions CD45.2^+^ WT recipient mice were also treated with α-IFN-γ blocking antibody at day 0, α-CD8 antibody (d-1, d2) or both antibodies in combination. dLNs were analyzed 5 days post infection. Quantification of T_FH_ and T_H_1 cells, expressed as percentages out of transferred Smarta CD4^+^ T cells. *n=6*. Mean ± SEM is shown. Data are representative of two independent experiments. One-way ANOVA with uncorrected Fisher’s LSD was applied. * *p-value* <= 0.05, *** *p-value* <= 0.001, **** *p-value* <= 0.0001. **E**) 0.5*10^6^ purified CD45.1^+^ Smarta CD4^+^ T cells from WT or Smarta-IFN-γ^-/-^ were transferred into CD45.2^+^ WT or IFN-γ^-/-^ recipients 1 day before s.c. rLCMV infection (1*10^5^ FFU /footpad). dLNs were analyzed 5 days post infection. Quantification of T_FH_ and T_H_1 cells, expressed as percentages out of transferred Smarta CD4^+^ T cells. *n=3*. Mean ± SEM is shown. Data are representative of three independent experiments. One-way ANOVA with uncorrected Fisher’s LSD was applied. * *p-value* <= 0.05, ** *p-value* <= 0.01, *** *p-value* <= 0.001, **** *p-value* <= 0.0001.

To explore the significance of IFN-γ produced by group 1 ILCs in CD4^+^ T cell polarization, we adoptively transferred Smarta CD4^+^ T cells into WT mice that were treated with neutralizing antibodies against IFN-γ or NK1.1, or a combination of both. Five days after LCMV infection group 1 ILCs were still efficiently depleted (**Fig. S10A**). In line with previous experiments, IFN-γ blocking led to a shift towards the T_FH_ cell subset (**Fig. 6B** and **C**) with a substantial increase of the Tcf-1^+^ population, a relative decrease of GzmB^+^ cells, and an increase of CXCR5 levels on Tcf-1^+^ cells (**Fig. S10B and C**). Notably, treatment with anti-NK1.1 antibody alone did not impact the polarization of T helper cell subsets. Moreover, the polarization pattern in mice treated with the combination of both antibodies mirrored the one observed with anti-IFN-γ alone (**Fig. 6B** and **C, and S10B and C**). Taken together, these findings suggest that, at least in this infection setting, IFN-γ derived from group 1 ILCs is not directly involved in CD4^+^ T cell polarization.

With group 1 ILCs ruled out, we explored other potential sources of IFN-γ. Monocytes and DCs, detected to produce IFN-γ 24 hours post-infection (**Fig. 6A**), were examined. Focusing on DCs, we employed a mixed bone marrow chimera model combining CD11c-DTR (for DC depletion) and IFN-γ^-/-^ mice. Depletion of WT DCs, leaving only IFN-γ^-/-^ DCs for T cell priming, did not impair CD4^+^ T cell differentiation (**Fig. S11A and B**). Similarly, the lack of monocytes in CCR2-deficient mice did not significantly impact CD4^+^ T cell polarization post-infection (**Fig. S12A-C**).

Subsequently, we focused on T cells, as they were identified as early IFN-γ producers (**Fig. 6A** and **Fig. 5G**). Depletion of CD8^+^ T cells slightly increased T_FH_ and Tcf-1^+^ subsets while reducing T_H_1 and GzmB^+^ subsets (**Fig. 6D** and **Fig. S13A and B**). A more pronounced effect was observed when combining CD8^+^ T cell depletion with IFN-γ blockade, suggesting that CD8^+^ T cells likely contribute to CD4^+^ T cell polarization through mechanisms other than IFN-γ (**Fig. 6D** and **Fig. S13A and B**).

Lastly, the role of CD4^+^ T cell-derived IFN-γ was examined. We transferred Smarta WT or IFN-γ^-/-^ CD4^+^ T cells into WT or IFN-γ^-/-^ recipients and assessed Smarta polarization post-infection. Consistent with earlier findings, the absence of IFN-γ in both donor and recipient led to an increase in T_FH_ cells and a marked decrease in T_H_1 cells (**Fig. 6E**). Smarta-derived IFN-γ effectively suppressed T_FH_ polarization in IFN-γ^-/-^ recipients, almost as efficiently as in WT recipients. However, when Smarta cells could not produce IFN-γ, a majority of CD4^+^ T cells still differentiated into T_H_1, implying that IFN-γ from endogenous CD4^+^ and CD8^+^ T cells might suffice for T_FH_ suppression (**Fig. 6E** and **Fig. S14**). These results collectively suggest that IFN-γ from adoptively transferred CD4^+^ T cells, as well as endogenous T cells, is responsible for T_FH_ suppression in the context of s.c. LCMV infection.

### IFN-γ Produced by T Cells Does Not Act in an Autocrine Fashion

Other groups have previously shown that IFN-γ produced by T_H_1 cells plays an instrumental role in maintaining and reinforcing the T_H_1 phenotype by acting in an autocrine fashion (Bradley *et al*., 1996; Zhang *et al*., 2001). To investigate whether early IFN-γ needed to suppress T_FH_ differentiation in the s.c. LCMV infection also targets CD4^+^ T cells, we applied Crispr/Cas9 technology to delete IFNGR1 from primary Smarta CD4^+^ T cells (**Fig. 7A**). Three control sgRNAs or three sgRNAs targeting IFNGR1 were transfected into naïve Smarta cells with high efficiency (**Fig. 7B**) prior to their transfer into WT recipient mice. Five days upon s.c. LCMV infection we found that Smarta cells had expanded similarly in the two groups of recipients (**Fig. 7C**) and Smarta cells receiving sgRNAs targeted to IFNGR1 displayed lower levels of the IFNGR1 protein (**Fig. 7D**). Notably, no difference at all was observed in CD4^+^ T cell polarization (**Fig. 7E**). To exclude any technical caveats we adopted an alternative approach by transferring into WT recipients genetically modified Smarta IFNGR1^-/-^ or Smarta WT mice (**Fig. 7F**). In addition, both experimental groups were treated or not with the IFN-γ blocking Ab. In this setting we detected significantly lower levels of IFNGR1 protein on Smarta IFNGR1^-/-^, as well as slightly lower levels of the receptor on WT Smarta in mice treated with the IFN-γ blocking Ab, suggesting that CD4^+^ T cells can upregulate the receptor when they sense IFN-γ (**Fig. 7G**). However, no significant differences were observed in the polarization between WT and IFNGR1^-/-^ cells (**Fig. 7H**). Instead, a notable T_FH_ rescue (expressing both CXCR5 and Bcl-6) was observed when mice were treated with the IFN-γ blocking Ab (**Fig. 7H**). All in all, these experiments show that IFN-γ released by T cells suppresses T_FH_ differentiation by acting on cells other than CD4^+^ T cells.

**Figure 7.**
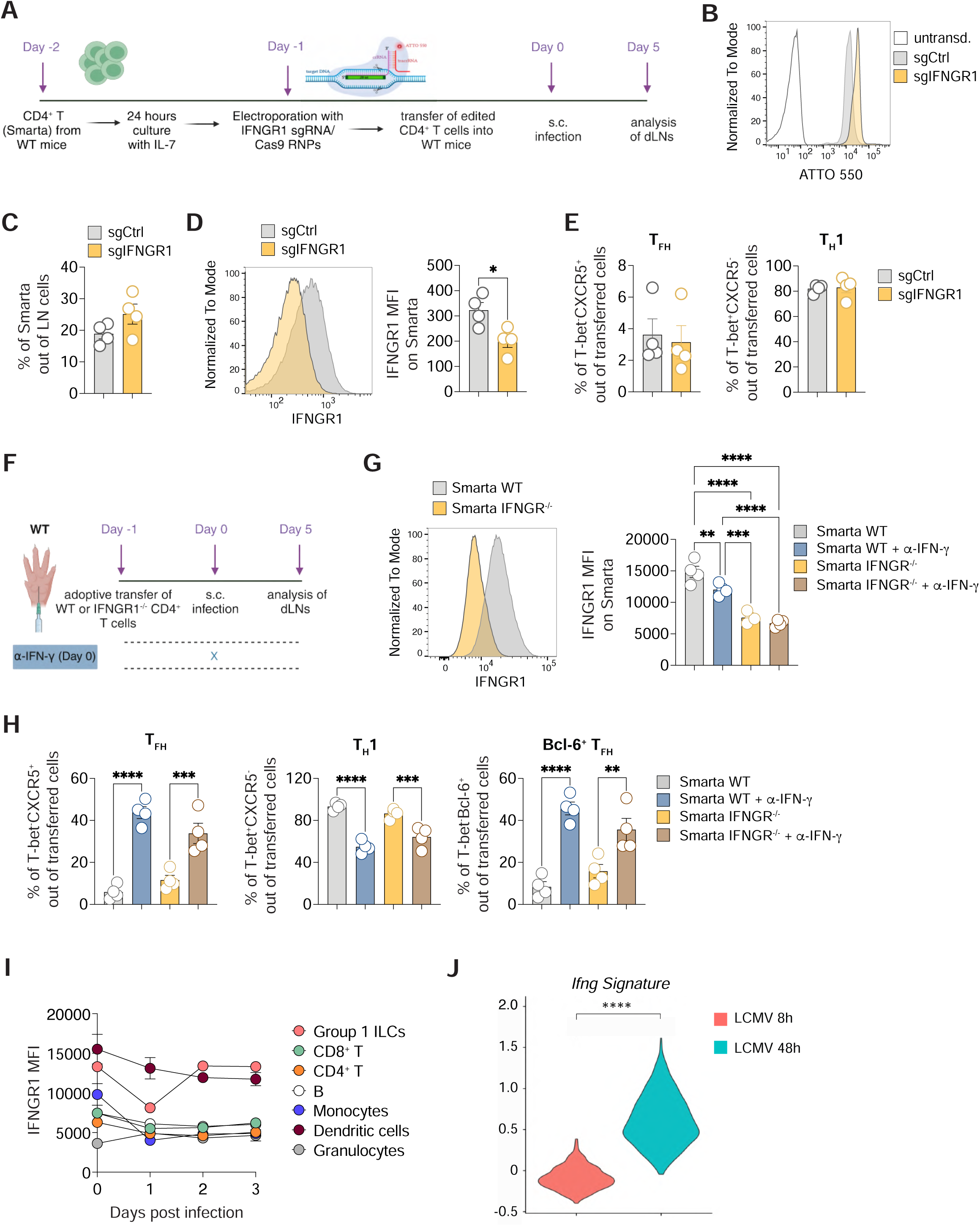
*IFN-γ Produced by T Cells Does Not Act in an Autocrine Fashion. **A**)* Total CD4^+^ T cells were isolated from spleens of naïve CD45.1^+^ Smarta Tg mice and cultured for 24 hrs with rIL-7. The day after, Smarta CD4^+^ T cells were transduced with control sgRNAs or with sgRNAs targeting IFNGR1 and with Cas9. After transduction, 0.5*10^6^ purified CD45.1^+^ Smarta CD4^+^ T cells were transferred into CD45.2^+^ WT 1 day before s.c. rLCMV infection (1*10^5^ FFU /footpad). dLNs were analyzed 5 days post infection. **B)** Representative plot of the transduction efficiency of Smarta CD4^+^ T cells is shown. **C)** Quantification of transferred Smarta CD4^+^ T cells expressed as percentages. *n=4*. Mean ± SEM is shown. Data are representative of two independent experiments. **D)** Representative flow cytometry plot (left panel) and quantification (right panel) showing the fluorescence intensity of IFNGR1 expression within Smarta CD4^+^ T cells of mice described in A. *n=4.* Mean ± SEM is shown. Data are representative of two independent experiments. An unpaired two-tailed t test was applied: \**p-value* <= 0.05. **E)** Quantification of T_FH_ (left) and T_H_1 (right), expressed as percentages out of transferred Smarta CD4^+^ T cells in dLNs of mice described in (A). *n=4*. Mean ± SEM is shown. Data are representative of two independent experiments. **F)** 0.5*10^6^ purified CD45.1^+^ WT or Thy1.1^+^ IFNGR1^-/-^ Smarta CD4^+^ T cells were transferred into CD45.2^+^ WT 1 day before s.c. rLCMV infection (1*10^5^ FFU /footpad). CD45.2^+^ WT recipient mice were also treated with α-IFN-γ blocking antibody (or isotype Ctrl) at day 0. dLNs were analyzed 5 days post infection. **G)** Representative flow cytometry plot (left panel) and quantification (right panel) showing the fluorescence intensity of IFNGR1 expression within Smarta CD4^+^ T cells of mice described in F. *n=4.* Mean ± SEM is shown. Data are representative of three independent experiments. An unpaired two-tailed t test was applied: \*\**p-value* <= 0.01, \*\*\**p-value* <= 0.001, \*\*\*\**p-value* <= 0.0001. **H)** Quantification of T_FH_ (left), T_H_1 (middle), and Bcl-6^+^ T_FH_ (right) expressed as percentages out of transferred Smarta CD4^+^ T cells in dLNs of mice described in (F). *n=4*. Mean ± SEM is shown. Data are representative of three independent experiments. \*\**p-value* <= 0.01, \*\*\**p-value* <= 0.001, \*\*\*\**p-value* <= 0.0001. **I)** Quantification of the mean fluorescence intensity of IFNGR1 expression on the indicated immune cells subsets in dLNs at the indicated time-points upon s.c. LCMV infection. *n=3.* Mean ± SEM is shown. **J)** Violin plot representation of the enrichment scores of the *Ifng* signature, comparing DCs sorted from mice infected with LCMV for 8 or 48 hours (published dataset in De Giovanni et al.). Two-tailed Mann-Whitney test has been performed, ****p < 0.0001.

To determine which cells of the LN might be the target for IFN-γ, we analyzed IFNGR1 protein levels on different immune cells in the first three days upon s.c. LCMV infection (**Fig. 7I**). As suggested by abovementioned results, CD4^+^ T cells did not express high levels of the receptor, and the same was for CD8^+^ T cells, B cells, and granulocytes. Instead, the highest levels of IFNGR1 were expressed by group 1 ILCs and DC (Fig. 7I). Since depletion of group 1 ILCs did not have any effect on CD4^+^ T cell polarization (**Fig. 6B-C**), we hypothesized that the possible target for IFN-γ suppressing T_FH_ differentiation might be DC. Analysis of a previous published dataset (De Giovanni *et al*., 2020) confirmed that DC sense IFN-γ during LCMV infection and express a signature of IFN-γ-stimulated genes 48 hours after infection (**Fig. 7J and S15A**). In addition, analysis of a published transcriptional dataset on WT and IFNGR2^-/-^ DCs during a parasitic infection (Lee *et al*., 2015) confirmed that DC that sense IFN-γ are characterized by a different transcriptional profile with respect to those that cannot sense this cytokine (**Fig. S15B**). In summary, our findings indicate that the IFN-γ responsible for T_FH_ suppression is T cell-derived but does not act on CD4^+^ T cells themselves. Instead, IFN-γ might be sensed by another cell type, likely DCs, which might then acquire a phenotype that interferes with T_FH_ differentiation.

### Exploring the role of IFN-γ Across Various Infection and Immunization Models

Diverging from s.c. infection dynamics, systemic LCMV infection notably induces a dual polarization of CD4^+^ T cells into both T_H_1 and T_FH_ subsets (Johnston *et al*., 2009; Hale *et al*., 2013; Ray *et al*., 2014; Weinstein *et al*., 2018). Echoing previous studies, we observed that approximately 40-50% of Smarta CD4^+^ T cells adopted a CXCR5^+^ profile indicative of T_FH_ cells on day 5 post systemic LCMV infection. The remaining population predominantly exhibited heightened T-bet expression (**Fig. S16A and B**). In this context, IFN-γ blockade distinctly influenced CD4^+^ T cell polarization, slightly increasing T_FH_ frequencies while decreasing T_H_1 cell proportions (**Fig. 8A** and **B**). However, this modulation only minimally altered Tcf-1^+^ and GzmB^+^ subsets, nor did it enhance CXCR5 expression within the Tcf-1^+^ subset, contrasting the patterns observed in s.c. infection routes (**Fig. 2B, C** and **Fig. S16C-E**).

**Figure 8.**
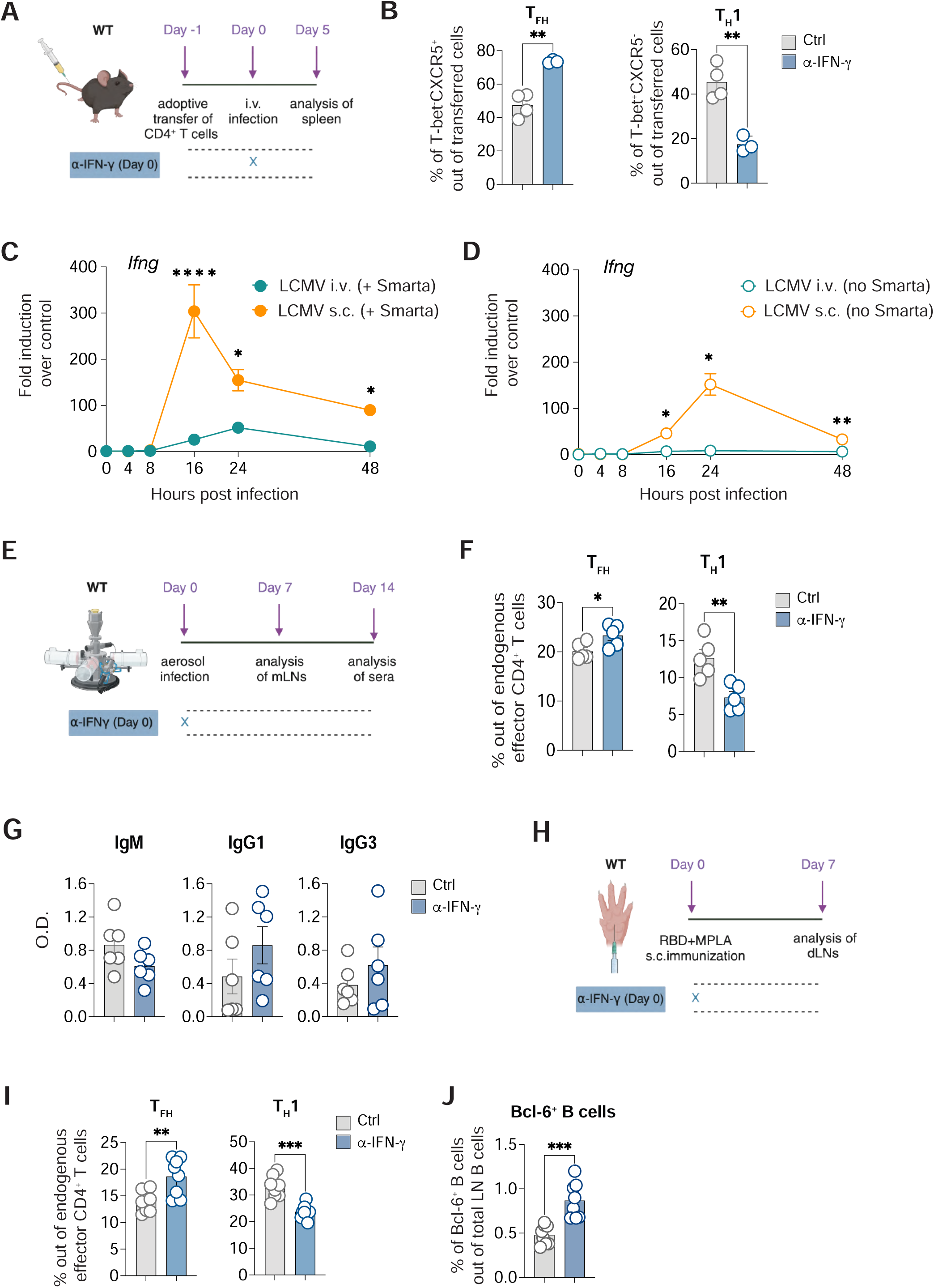
*Dissecting the role of IFN-γ in other infection and immunization settings. **A**)* 0.5*10^6^ purified CD45.1^+^ Smarta CD4^+^ T cells were transferred into CD45.2^+^ WT recipients 1 day before intravenous (i.v.) rLCMV infection (2*10^5^ FFU). CD45.2^+^ WT recipient mice were also treated with α-IFN-γ blocking antibody (or isotype Ctrl) at day 0. Spleens were analyzed 5 days post infection. **B)** Quantification of T_FH_ (left) and T_H_1 (right), expressed as percentages out of transferred Smarta CD4^+^ T cells in spleens of mice described in (A). *n=*4 (Ctrl), 3 (α-IFN-γ). Mean ± SEM is shown. Data are representative of three independent experiments. An unpaired two-tailed t test was applied. ** *p-value* <= 0.01. **C**) 0.5*10^6^ purified CD45.1^+^ Smarta CD4^+^ T cells were transferred into CD45.2^+^ WT recipients 1 day before i.v. (2*10^5^ FFU) or s.c. (1*10^5^ FFU /footpad) rLCMV infection. Analysis of *Ifng* gene expression at 0, 4, 8, 16, 24 and 48 hours in dLN (orange) and spleens (green) of mice infected s.c. (orange) and i.v. (green) with rLCMV is shown. *n=4-6*. Mean ± SEM is shown. Two-way ANOVA with uncorrected Fisher’s LSD was applied. * *p-value* <= 0.05, **** *p-value* <= 0.0001. **D)** WT mice were infected i.v. (2*10^5^ FFU) or s.c. (1*10^5^ FFU /footpad) with rLCMV. Analysis of *Ifng* gene expression at 0, 4, 8, 16, 24 and 48 hours in dLN (orange) and spleens (green) of mice infected s.c. (orange) and i.v. (green) with rLCMV is shown. *n=3-4*. Mean ± SEM is shown. Two-way ANOVA with uncorrected Fisher’s LSD was applied. * *p-value* <= 0.05, ** *p-value* <= 0.01. **E)** WT mice were infected via aerosol with a mouse-adapted strain of SARS-CoV-2 and mediastinal LNs (mLNs) were analyzed 7 days upon infection. **F)** Quantification of T_FH_ (left) and T_H_1 (right), expressed as percentages out of endogenous effector CD4^+^ T cells in mLNs of infected mice described in (G). *n*=5. Mean ± SEM is shown. Data are representative of two independent experiments. An unpaired two-tailed t test was applied. * *p-value* <= 0.05, ** *p-value* <= 0.01. **G)** RBD–binding IgM, IgG1 and IgG3 Abs were measured in the sera of mice 14 days upon aerosol infection as described in (E) and expressed as O.D. *n*=6. Mean ± SEM is shown. **H)** WT mice were immunized s.c. with MPLA+ RBD-S1 and treated with α-IFN-γ blocking antibody (or isotype Ctrl) at day 0. dLNs were analyzed 7 days post infection. **I)** Quantification of T_FH_ (left) and T_H_1 (right), expressed as percentages out of endogenous effector CD4^+^ T cells in dLNs of s.c. infected mice described in (H). **J)** Quantification of Bcl-6^+^ B cells expressed as percentages out of endogenous B cells in dLNs of s.c. infected mice described in (H).*n=*8. Mean ± SEM is shown. Data are representative of three independent experiments. An unpaired two-tailed t test was applied. ** *p-value* <= 0.01, *** *p-value* <= 0.001.

We hypothesized that these observed differences might stem from varying IFN-γ levels induced by different infection routes. Indeed, quantitative PCR analysis revealed that systemic infection elicited lower *Ifng* levels compared to s.c. infection (**Fig. 8C**). Considering potential overestimation of *Ifng* expression due to the adoptive transfer of antigen-specific CD4^+^ T cells, we assessed endogenous *Ifng* expression in non-transferred mice. This confirmed significant *Ifng* induction only in s.c.ly infected mice (**Fig. 8D**). This finding aligns with previous reports suggesting that systemic LCMV infection’s low *Ifng* expression could be a result of type I IFNs inhibition (Cousens *et al*., 1997b; Nguyen *et al*., 2000; Pien and Biron, 2000; Nguyen *et al*., 2002; Miyagi *et al*., 2007). Consistently, endogenous CD4^+^ T cell polarization post systemic infection remained unchanged despite IFN-γ blockade (**Fig. S16F-H)**, whereas as shown before s.c. infection saw a notable rescue of T_FH_ cells (**Fig. 4**).

To probe for a possible role of IFN-γ in affecting CD4 T cells responses in other infection settings, we decided to analyze CD4^+^ T cell polarization in the context of a mouse model for SARS-CoV-2 infection. To this end we used a mouse-adapted SARS-CoV-2 strain, rSARS-N501YMA30 (Wong *et al*., 2022) which was administrated to WT mice via aerosol (Fumagalli *et al*., 2021, 2024) (**Fig. 8E**). We found that SARS-CoV-2 infection induced a balanced CD4^+^ T cell response consisting of both T_H_1 and T_FH_ cells: however, when blocking IFN-γ, T_H_1 cells decreased significantly, whereas T_FH_ frequency increased (**Fig. 8F**), confirming the role of IFN-γ as a molecular switch shifting the balance towards cellular responses. Moreover, two weeks after infection, we observed a trend in the increase of IgG1 and IgG3 antibodies, which have been reported to be the only T_FH_-dependent isotypes (Chen *et al*., 2022) (**Fig. 8G**).

Broadening our investigation, we explored the role of IFN-γ in a translational context using a monophosphoryl lipid A (MPLA)-based immunization model known to foster T_H_1 differentiation (Mata-Haro *et al*., 2007; Komai-Koma *et al*., 2021). Mice were immunized with MPLA-conjugated SARS-CoV-2 RBD protein, and the effects of IFN-γ blockade were analyzed (**Fig. 8I**). Immunization resulted in increased expression of *Ifng* especially at 16h (**Fig. S16I**). Intriguingly, inhibiting IFN-γ in this setting also led to reduced T_H_1 cells, and increased T_FH_ and Bcl-6^+^ B cells (**Fig. 8I** and **J**). These findings collectively suggest that both in viral infection and immunization scenarios, IFN-γ functions to suppress T_FH_ responses, and its blockade could enhance humoral vaccine responses.

## Discussion

CD4^+^ T cell polarization plays a critical role in shaping effector immune responses (Tuzlak *et al*., 2021). Previously, we demonstrated that s.c. LCMV infection is characterized by a stark compartmentalization of CD4^+^ T cell responses, leading to almost exclusive T_H_1 polarization but severely impaired T_FH_ cell differentiation (De Giovanni *et al*., 2020). Building upon these findings, our current research reveals that IFN-γ, known for supporting type I responses and maintaining the T_H_1 phenotype, plays a pivotal role in suppressing T_FH_ differentiation and B cell responses during viral infections (**Fig. S17**). Importantly, we found that the IFN-γ responsible for T_FH_ suppression is T cell-derived but does not act on CD4^+^ T cells themselves, unlike to what previously reported for T cell expansion and T_H_1 maintenance (Bradley *et al*., 1996; Lighvani *et al*., 2001; Whitmire *et al*., 2005). Instead, IFN-γ might be sensed by DCs, which might then acquire a phenotype that interferes with T_FH_ differentiation. Thus, we propose that T_H_1 antagonize T_FH_ not only through competition of cell-intrinsic transcription factors as proposed by others (Nakayamada *et al*., 2011), but also through a cell-extrinsic effect on the surrounding microenvironment.

The inhibitory effect of IFN-γ on T_FH_ differentiation and humoral immunity is not unprecedented. For instance, in severe malaria infection, a setting where T_H_1-like dysfunctional T_FH_ cells expressing T-bet are observed, concomitant blockade of IFN-γ and TNF-α resulted in enhanced T_FH_ differentiation and improved antibody responses (Ryg-Cornejo *et al*., 2016). Nevertheless, in this context, the cells responsible for producing or sensing IFN-γ were not identified, and the underlying mechanism remained unexplored. In another study performed with bacterial infection, IFN-γ produced by B cells upon IL-12 sensing was shown to act in an autocrine fashion and to suppress GC reactions (Elsner *et al*., 2024). Our research indicates that, at least upon viral infection, IFN-γ specifically produced by T cells suppresses functional T_FH_ differentiation by likely acting on DC. Since IFN-γ was found to be produced in the first three days upon infection, we hypothesize that the first differentiated T_H_1 cells might antagonize the arising T_FH_ cells by possibly modifying the dLNs microenvironment.

Notably, we found that IFN-γ does not influence only CD4^+^ T cell polarization but also the B cell phenotype, as shown by restriction of the development of Bcl-6^+^ B cells, commonly identified as GC B cells. A role for IFN-γ on B cell activation has been reported by others although with different outcomes depending on the context (Abed *et al*., 1994; Myles *et al*., 2017; Obeng-Adjei *et al*., 2017; Unger *et al*., 2018; Stone *et al*., 2019; Zumaquero *et al*., 2019; Chodisetti *et al*., 2020; Arroyo-Díaz *et al*., 2023). In particular previous literature has suggested that IFN-γ can act directly on B cells to induce T-bet expression (Obeng-Adjei *et al*., 2017; Stone *et al*., 2019; Zumaquero *et al*., 2019; Chodisetti *et al*., 2020). T-bet-expressing B cells, observed in various chronic infections, severe malaria, and autoimmune diseases, are often characterized as ‘atypical’ due to their dysfunctional traits and pathogenicity (Obeng-Adjei *et al*., 2017; Rubtsov *et al*., 2017; Rubtsova *et al*., 2017; Burton *et al*., 2018). However, in the context of respiratory viral infections, T-bet-expressing T and B lymphocytes function collaboratively, facilitating optimal antiviral immunity (Mendoza *et al*., 2021). In our experimental setting IFNGR1 expression on B cells was not very high in the first three days upon infection, therefore we believe it is likely that the increase in Bcl6^+^ B cells may result from enhanced T_FH_ differentiation following early IFN-γ blockade, whereas the decrease in T-bet^+^ B cells could be due to IFN-γ action directly on B cells at later time-points.

Recently, a population of Tcf-1^+^Bcl-6^low^PD-1^+^ CD4^+^ T cells that can give rise to both effector cells and T_FH_ cells has been described in a setting of chronic LCMV infection and antigen persistence (Xia *et al*., 2022). We cannot formally exclude that this population functionally resembles Tcf-1^+^T-bet^+^ CD4^+^ T identified in our setting: however, it is worth mentioning that the s.c. LCMV infection is cleared within one week and therefore represents an acute infection setting (Sammicheli *et al*., 2016).

Notably, IL-12, a well-established T_H_1-polarizing cytokine (Hsieh *et al*., 1993; Heufler *et al*., 1996; Athie-Morales *et al*., 2004) was not essential for T_H_1 differentiation in LCMV infection, as echoed by other studies (Schijns *et al*., 1998; Oxenius *et al*., 1999). This may be explained by the poor induction of IL-12 in certain viral infections due to type I IFNs’ inhibitory effects (Cousens *et al*., 1997a; Pien and Biron, 2000). IFN-γ, while known to promote T_H_1 phenotype survival and even act as a polarizing cytokine alongside IL-12 in some contexts (Bradley *et al*., 1996; Heufler *et al*., 1996; Wakil *et al*., 1998; Lighvani *et al*., 2001; Miro *et al*., 2006; Schulz *et al*., 2009), was crucial for the development of at least one identified T_H_1 subset expressing both *Gzma* and *Gzmb*. The detailed characterization of this subset warrants further investigation.

Our study indicates that the pronounced CD4^+^ T cell compartmentalization observed in s.c. LCMV infection, leading to a dominant T_H_1 response, is likely attributed to the high levels of IFN-γ induced in this infection route. Indeed, we suggest that reduced IFN-γ induction in systemic infections (Cousens *et al*., 1997b; Nguyen *et al*., 2000; Pien and Biron, 2000; Nguyen *et al*., 2002; Miyagi *et al*., 2007) permits a coexistence of T_H_1 and T_FH_ cells (Johnston *et al*., 2009; Hale *et al*., 2013; Ray *et al*., 2014; Weinstein *et al*., 2018). In scenarios where IFN-γ is overexpressed, such as upon transfer of antigen-specific CD4^+^ T cells, blocking IFN-γ results in increased T_FH_ cell frequencies regardless of the infection route. However, IFN-γ was shown to restrict T_FH_ development also in endogenous settings of not only s.c. LCMV infection, but also of SARS-CoV-2 infection or with an immunization approach. These findings imply a universal role for IFN-γ in suppressing humoral responses across various contexts. Crucially, this insight could guide the development of more effective vaccination strategies. Future studies delving deeper into the spatiotemporal mechanisms employed by this cytokine could provide further clarity and direction for innovative therapeutic approaches.

## Materials and Methods

### Mice

Mice were housed under specific pathogen-free conditions and used at 8-10 weeks of age, unless otherwise indicated. In all experiments female mice matched for age were used. All experimental animal procedures were approved by the Institutional Animal Committee of the San Raffaele Scientific Institute and by the Italian Ministry of Health. C57BL/6 and C57BL/6-Ly5.1 (CD45.1) (inbred C57BL/6) mice were purchased from Charles River. CCR2^-/-^ (B6.129S4-Ccr2tm1Ifc/J), IFNg-YFP (129S4(B6)-Ifnγtm3-1Lky/J), IFNGR1^-/-^ (B6.129S7-Ifngr1tm1Agt/J) and IFN-γ^-/-^ (B6.129S7-Ifngtm1Ts/J) mice were purchased from The Jackson Laboratory. Mice bearing LCMV-specific transgenic CD4^+^ T cells (Smarta) were obtained through the Swiss Immunological Mouse Repository (SwImMR). CD11c-DTR mice have been described previously (Jung *et al*., 2002) and were obtained from M. De Palma and L. Naldini (San Raffaele Scientific Institute). Bone marrow (BM) chimeras were generated by irradiation of C57BL/6 mice with ∼900 rad and reconstitution with the indicated bone marrow; mice were supplied with antibiotic-supplemented water and allowed to reconstitute for at least 8 weeks prior to use.

### Infections and immunizations

Mice were infected s.c.ly (s.c.) in the footpad with 1×10^5^ Plaque-Forming Unit (PFU) of rVSV (a recombinant VSV expressing a GP derived from the LCMV WE strain and recognized by Smarta TCR-transgenic instead of the VSV GP) or with 1×10^5^ Focus-Forming Unit (FFU) of rLCMV (a recombinant LCMV clone 13 expressing a GP derived from the LCMV WE strain and recognized by Smarta TCR-transgenic instead of the LCMV Cl13 GP) (Fallet *et al*., 2016; De Giovanni *et al*., 2020). In indicated experiments, mice were infected intravenously (i.v.) with 2×10^5^ FFU of rLCMV. Viruses were propagated and quantified as described in previous studies (Kuka *et al*., 2012; Sammicheli *et al*., 2016; De Giovanni *et al*., 2020) and were diluted in 25μl of HBBS before s.c. footpad injection. Viral titers from dLNs of LCMV-infected mice were measured by focus assay. Infection of C57BL/6 mice with aerosolized SARS-CoV-2 was performed as described (Fumagalli *et al*., 2021, 2024). Briefly, non-anesthetized mice were placed in a nose-only Allay restrainer on the inhalation chamber (DSI Buxco respiratory solutions; DSI). To reach a target accumulated inhaled aerosol (also known as delivered dose), C57BL/6 mice were exposed to a target accumulated inhaled aerosol of the mouse-adapted SARS-CoV-2 strain (rSARS-N501YMA30) kindly provided by Stanley Perlman. Primary inflows and pressure were controlled and set to 0.5 l min−1 per port and −0.5 cmH2O, respectively. Infected mice were monitored daily to record body weight and clinical and respiratory parameters. All infectious work was performed in designated Biosafety Level 2 (BSL-2) and BSL-3 workspaces in accordance with institutional guidelines.

In immunization settings, mice were injected s.c. in the footpad with 5 μg of rRBD (Sars Cov-2 (2019-nCoV) Spike RBD Protein (S1 Subunit,FC Tag) Sino Biological #40592-V02H) conjugated to 10 μg of MPLA-SM* VacciGrade™ (vac-mpla2) in a volume of 30 μl/footpad.

### T cell isolation, adoptive transfer, and in vivo treatments

CD4^+^ T cells were negatively selected from spleens of naïve Smarta CD45.1^+^ transgenic mice by magnetic isolation (Miltenyi Biotec), with purity always above 98% as determined by flow cytometry. Unless otherwise indicated, 0.5×10^6^ Smarta T cells were injected i.v. into indicated recipients one day before infection. In indicated experiments, mice were treated with: InVivoMab α-IFN-γ blocking antibody (BioXcell Clone XMG1.2 #BE0055) or rat IgG1 isotype control (BioXcell Clone HRPN #BE0088): 250 μg intraperitoneally (i.p.) at day 0 (or in selected experiments at day 3); InVivoMab α-IL-12 blocking antibody (BioXcell Clone R2-9A5 #BE0233) or rat IgG2b isotype control, (BioXcell Clone LFT-2 #BE0090): 1 mg i.p. at day 0 and 3 after infection; InVivoMab α-NK1.1 depleting antibody (BioXcell Clone PK136 #BE0036): 1 mg i.p. at day 0 and day 1 after infection; InVivoMab α-CD8 depleting antibody (BioXcell Clone YTS 169.4 #BE0117): 200 μg i.p. at day-1 and day 2 after infection.

To deplete DCs 500 ng of diphtheria toxin (DTX, Millipore, #322326) diluted in 200μl of PBS was administered i.p. one day before the infection and every other day thereafter to CD11c-DTR/IFN-γ^-/-^ and CD11c-DTR/WT BM chimeras, respectively.

### CRISPR/Cas9-Mediated IFNGR1 Knockout in Primary CD4^+^ T Cells

CRISPR/Cas9-Mediated IFNGR1 knockout was performed following the protocol in (Oh *et al*., 2019). Briefly, CD4^+^ T cells were negatively selected from spleens of naïve Smarta CD45.1^+^ transgenic mice by magnetic isolation (Miltenyi Biotec), with purity always above 98% as determined by flow cytometry. Cells were then resuspended at a concentration of 10^6^/ml and cultured with recombinant IL-7 (5 ng/ml) overnight at 37°C. Cas9/gRNA RNP complexes containing sgRNAs (either control or targeted to IFNGR1) and cas9 were assembled following the protocol provided by the supplier (Integrated DNA Technologies, IDT). CD4^+^ T cells were resuspended in nucleofection buffer and transfected with Cas9/RNP complexes following the instructions of the Primary Cell 4D-Nucleofector X Kits (Lonza). Cells were then analyzed by flow cytometry to evaluate the transfection efficiency (through fluorescence conferred by ATTO550).

### Single cell suspensions and flow cytometry

Single-cell suspensions of spleens and LNs were obtained by mechanical dissection as previously described (Kuka *et al*., 2012; Sammicheli *et al*., 2016; De Giovanni *et al*., 2020; Fiore *et al*., 2023). Red blood cells were lysed with ammonium chloride (ACK) lysis buffer. In selected experiments, cell suspensions were plated in round-bottom 96-well plates (1*10^6^ cells/well) and restimulated for 4 hours with 2 μM GP61–80 peptide from LCMV (GLKGPDIYKGVYQFKSVEFD) in the presence of Brefeldin A (GolgiPlug, 1 ml/ml), in RPMI supplemented with 10% fetal bovine serum.

All flow cytometry stainings of surface-expressed markers were performed in FACS Buffer containing PBS and 2% FBS at 4°C, while intracellular molecule staining was performed using Foxp3/Transcription Factor Staining Buffer set (eBioscience, #00-5523-00), following the manufacturer’s instructions at room temperature. Anti-CD16/32 antibody (Invitrogen # 14-0161-82) was added to cell pellets prior to staining with fluorochrome-conjugated antibodies to block Fc receptors. Antibodies (Abs) used were purchased from BD Bioscience, Invitrogen, Biolegend or Cell Signalling and are indicated in **Table 1**. Flow cytometry analyses were performed on BD FACSCanto II, BD FACSymphony A5 or Cytek Aurora and analysed with FlowJo software (Treestar).

| Antibodies |  |
| --- | --- |
| Rat anti-mouse CD185 (CXCR5) (2G8) | Rat anti-mouse CD8a (53-6.7) |
| Rat anti-mouse T-bet (4B10) | Rat anti-mouse CD4 (RM4-5) |
| Mouse anti-mouse/human Granzyme B (GB11) | Mouse anti-mouse CD45.1 (A20) |
| Rabbit anti-mouse TCF1/TCF7 (C63D9) | Rat anti-mouse/human CD11b (M1/70) |
| Rat anti-mouse IFN-γ (XMG1.2) | Rat anti-mouse Ly-6G (1A8) |
| Mouse anti-mouse Bcl-6 (K112-91) | Hamster anti-mouse CD11c (HL3) |
| Rat anti-mouse CD186 (CXCR6) (SA051D1) | Mouse anti-mouse I-A<sup>b</sup> (AF6-120.1) |
| Hamster anti-mouse CD183 (CXCR3-173) | Rat anti-mouse CD44 (IM7) |
| Rat anti-mouse Ly-6C (HK1.4) | Rat anti-mouse CD62L (MEL-14) |
| Rat anti-mouse CD45R/B220 (RA3-6B2) | Rat anti-mouse TCR V alpha 2 (B20.1) |
| Mouse anti-mouse NK-1.1 (PK136) | Hamster anti-mouse Vβ 8.3 TCR (1B3.3) |
| Rat anti-mouse CD335 (NKp46) (29A1.4) | Rat anti-mouse IFNGR1 (GR20) |

### Confocal Immunofluorescence Staining

For confocal microscopy analysis LNs were directly collected and incubated for 90 min at room temperature in Antigen Fix (Diapath #P0016), and then washed in DPBS and dehydrated in 30% Sucrose at 4°C. LNs were embedded in OCT freezing media (Killik Bio-Optica #05-9801) and 20 μm cryosections were prepared on a CM1520 cryostat (Leica), adhered to Superfrost Plus slides (Thermo Scientific) and stored at −20°C. Sections were permeabilized and blocked with Blocking Buffer composed of DPBS, 10% FCS and 0.3% Triton X-100 (Sigma-Aldrich) and stained in the same buffer. Anti-CD16/32 antibody (Invitrogen # 14-0161-82) was added to cell pellets prior to staining with fluorochrome-conjugated antibodies to block Fc receptors. Before staining with fluorochrome-conjugated antibodies, slides were stained with Anti-mouse Fc Block antibody to block non-specific binding sites. The following fluorochrome-conjugated antibodies were used for cryosections staining: rat αB220 (RA3-6B2), mouse αCD45.1 (A20) and rabbit αTCF1/TCF7 (C63D9). Stained slides were mounted with FluorSave^TM^ Reagent (Merck Millipore, #345789) and Images were acquired on an inverted Leica microscope (SP8, Leica Microsystems) with a motorized stage for tiled imaging using an HC PL APO CS 20X (NA 0.7) Dry or HCX PL APO λ blue 40X (NA 1.25) Oil objectives. To minimize fluorophore spectral spillover, we used the Leica sequential laser excitation and detection modality. B cell follicles were defined on the basis of B220 staining. For three-dimensional image acquisition, 6-10 xy stacks (1,024×1,024 pixels) sampled with 2-μm z spacing were acquired to provide image volumes that were 20μm in depth.

### ELISA

The GP-1-IgG ELISA was carried out in 96-well half-volume polystyrene plates (Corning #3690). Plates were coated overnight at 4°C with goat α-human IgG Fc capturing Ab (Jackson Immunoresearch #109005098) diluted 1:1000 in 0.1 M sodium carbonate buffer (pH 9.6). Afterward, the plates were blocked for 1 h with 5% milk diluted in PBS-T. Thereafter, the plates were incubated with 50 μl per well of GP-1-IgG-containing cell supernatant for 1 h. Sera were diluted 1:4 in 5% milk in the first 96-well row and then 1:2 serial dilution were carried out in the GP-1-IgG-saturated plates, followed by incubation for 1 h. Finally, the plates were incubated for 1h with HRP Goat anti-mouse IgG (PerkinElmer NEF822001EA) diluted 1:500 in 0,5% BFA. HRP was detected by using TMB Substrate Reagent set (BD Bioscience #555214). All steps were carried out at room temperature. Between each step the plates were washed five times with PBS-

### T. Titers represent double-above-background values

The SARS-CoV-2 S1 RBD-specific ELISA was carried out by coating plates with recombinant SARS-CoV-2 S1 subunit protein (RayBiotech, 230-30162) at a concentration of 2 μg ml/1 in PBS and incubated overnight (O/N) at 4 °C. Subsequently, the plates were blocked with PBS containing 1% bovine serum albumin (PBS-1%BSA) for 1 h at room temperature. The sera were then added at a dilution of 1/20 (sera from day 7) or 1/500 (sera from days 14, 21 and 28) and diluted 1:10 up to 1/1,280 or 1/32,000, respectively, in duplicate, and the plates were incubated for 2 h at room temperature. After five washes with 0.05% Tween 20 in PBS, the secondary anti-mouse IgM, IgG1 or IgG3 conjugated with horseradish peroxidase (SouthernBiotech # 5300-05B) was added and the plates were incubated for 1 h at room temperature. After washing, the binding of the secondary antibody was detected by adding the substrate 3,3′,5,5′-tetramethylbenzidine (BD Biosciences). The reaction was blocked with 0.5 M H2SO4 and the absorbance at 450 nm and reference 630 nm was measured.

### Single-cell RNA sequencing (1) – library preparation

Single cell populations from rVSV-infected mice (CD4^+^CD45.1^+^or CD4^+^ CD45.1^+^PD-1^+^ICOS^+^405 cells), rLCMV-infected mice or not infected SMARTA mice (CD4^+^CD45.1^+^) were sorted by using the following flow cytometry antibodies: APC-CXCR5 (2G8; BD Biosciences), APC-Cy7-CD45.1 (A20; Biolegend), eFluor450-CD4 (RM4-5; eBioscience), BV605-ICOS (C398.4A; Biolegend), PE-PD-1 (J43; eBioscience), AxFl488-B220 (RA3-6B2; Biolegend), AxFl488-NK1.1 (PK136; Biolegend), PE-Cy7-CD8a (53-6.7; Biolegend). LIVE/DEAD Fixable Aqua Dead Cell Stain (ThermoFisher Scientific) was used to exclude dead cells. Sorting was performed following exclusion of doublets, dead cells, and B220^+^B cells, NK1.1^+^NK cells, CD8a^+^T cells, and Ter119^+^erythrocytes.

Single-cell libraries were prepared as previously described (Jaitin *et al*., 2014). Briefly, mRNA from cells sorted into cell capture plates was barcoded, converted to cDNA and pooled with an automated pipeline. The pooled sample was then linearly amplified by T7 in vitro transcription, and the resulting RNA was fragmented and converted to a sequencing-ready library by tagging the samples with pool barcodes and Illumina sequences during ligation, reverse transcription and PCR. Each pool of cells was tested for library quality, and the concentration was assessed as described (Jaitin *et al*., 2014). RNA-seq libraries were sequenced on an Illumina NextSeq 500 platform, at a median sequencing depth of 15,054 reads per cell. Sequences were mapped to the mouse genome (mm10), demultiplexed, and filtered as previously described (Berglund *et al*., 2018), with the following adaptations. Mapping of reads was done using HISAT (version 0.1.6); reads with multiple mapping positions were excluded. Reads were associated with genes if they were mapped to an exon, using the UCSC genome browser for reference. We estimated a median of 2% spurious UMI in the data using statistics on empty MARS-seq wells.

### Single-cell RNA sequencing (2) – library preparation

CD4^+^ Smarta T cells from rLCMV-infected mice (day 5) treated or not with α-IFN-γ blocking antibody were sorted on MACSQuant Tyto Cell Sorter (Miltenyi Biotec) by using the following flow cytometry antibodies: PE-CD45.1 (A20; Biolegend), eFluor450-CD4 (RM4-5; eBioscience). LIVE/DEAD Fixable Near-IR Dead Cell Stain (ThermoFisher Scientific) was used to exclude dead cells. Sorting was performed following exclusion of doublets and dead cells.

Single cells were processed on the Chromium platform (10x) using the Chromium Next GEM Single Cell 3’ v3.1 (Dual Index). After quality controls and quantification on TapeStation instrument (Agilent), libraries were sequenced on NovaSeq 6000 platform (Illumina) generating around 18,000 reads/cell. Raw sequencing data were demultiplexed with the mkfastq application (Cell Ranger v.6.0.2). UMI-Tools (v.1.0.0) whitelist and extract commands were used to identify and select the number of cell barcodes to use in downstream analysis. Reads were mapped to the reference genome using STAR v.2.5.3a and assigned to genes with featureCounts v.1.6.4. GRCm38 was used as the reference genome. Bam files were sorted with samtools software (v1.9). Finally, Umi-Tools count was used to processing the UMIs aligned to each gene in each cell to find the number of distinct, error-corrected UMIs mapping to each gene. The UMI count tables of each cellular barcode were used for further analysis.

### Single-cell RNA sequencing bioinformatics analysis

Single cell data analysis was performed using Seurat (v4.0.2). Cells with sufficient bioinformatic quality were obtained after applying a filter of at least 200 genes expressed per cell and only genes expressed in at least 5 cells were retained. Moreover, cells with more than 10% of reads mapped to mitochondrial genes were also excluded from the analysis. UMI count matrix was further normalized and scaled following the standard Seurat workflow and Umap reduction was then applied on first 30 Principal Components after running PCA. Unbiased clustering was computed using the FindClusters function in Seurat with default parameters and a resolution value of 0.4. Specific markers for the different unbiased clusters were found using the function FindAllmarkers or FindMarkers in Seurat with default parameters. The plots showing normalized expression values with a color scale on top of Umap plots (on Fig. 1E, 2H, 2J, S2A) and the Violin plots of specific genes were produced with FeaturePlot and VlnPlot Seurat functions, respectively. The max.cutoff parameter is set to “q95”. The gene signature in Fig. 2K was calculated with the AddModuleScore function in Seurat.

### qPCR

Total RNA was isolated from frozen LNs or spleens with the ReliaPrep RNA Miniprep system (Promega), following the manufacturer’s instructions. One microgram of total RNA was reverse transcribed before qPCR analyses for *Ifng* (Mm01168134_m1) in a QuantStudio 5 Real-Time PCR System (all from Thermo Fisher Scientific). All experiments were done in duplicate, and data were normalized to the housekeeping gene *Gapdh* (Mm99999915_g1, Thermo Fisher Scientific).

### Statistical analyses

Flow and imaging data were collected using FlowJo Version 10.5.3 (Treestar) and Imaris (Bitplane), respectively. Statistical analyses were performed with GraphPad Prism software version 9.5 (GraphPad). Results are expressed as mean ± SEM. Means between two groups were compared with unpaired two-tailed t test. Means among three or more groups were compared with one-way or two-way ANOVA. Uncorrected Fisher LSD post-test was used for multiple comparisons. Significance is indicated as follows: * *p-value* < 0.05; ** *p-value* < 0.01; *** *p-value* < 0.001; **** *p-value* < 0.0001. Comparisons are not statistically significant unless indicated.

## Data and materials availability

The data that support the findings of this study are openly available in the San Raffaele Open Research Data Repository at DOI: 10.17632/c9r2fwjhr4.1 (reserved link until acceptance). The scRNA-seq data are available in the Gene Expression Omnibus (GEO) database under accession no. GSE239968.

## Author contributions

Conceptualization: M.K.; Investigation: E.S., M.N., C.L., P.D.L., E.B.B., M.M., D.M., V.S., M.G., L.G., J.N., D.K., G.F., C.M., E.C.; Single cell RNA-seq experiment: C.G.B., E.D., M.C., A.G.; Resources: M.K., M.I., I.A., B.B., R.B., L.G.G.; Formal Analysis, E.S., M.N., M.K.; Bioinformatic analysis: C.L., F.T.; Writing: M.K. with input from E.S., C.L., and M.I.; Editing: all authors.; Visualization, E.S, M.K.; Project supervision, M.K.; M.I.; Funding Acquisition, M.K., M.I., L.G.G.

## Acknowledgments

We thank M. Silva and M. Tinelli for secretarial assistance and the members of the Kuka and Iannacone Laboratory for helpful discussions. Flow cytometry was carried out at FRACTAL, a flow cytometry resource and advanced cytometry technical applications laboratory established by the San Raffaele Scientific Institute. Confocal immunofluorescence histology was carried out at Alembic, an advanced microscopy laboratory established by the San Raffaele Scientific Institute and the Vita-Salute San Raffaele University. We thank D. Pinschewer for rLCMV and Stanley Perlman for the mouse-adapted SARS-CoV-2 strain (rSARS-N501YMA30). E.S., C.L., and M.N. conducted this study as partial fulfillment of their PhD in the PhD program in Basic and Applied Immunology and Oncology at Vita-Salute San Raffaele University.

This research was supported by the Italian Ministry of University and Research grants PRIN-P2022YKH9R, PRIN-2017ZXT5WR, and SIR-RBSI14BAO5 to M.K. M.K. is further supported by the Italian Ministry of University and Research grants PRIN-20209Y5YFZ and PE00000007 (INF-ACT), and by the Italian Ministry of Health (MoH) Grant GR-2021-12372615. M.I. is supported by European Research Council (ERC) Consolidator Grant 725038, ERC Proof of Concept Grant 957502, Italian Association for Cancer Research (AIRC) Grants 19891 and 22737, Italian Ministry of Health (MoH) Grant RF-2018-12365801, Italian Ministry for University and Research (Project no. PE00000007, INF-ACT), and sponsored research agreements from Gilead Sciences, Asher Biotherapeutics and VIR Biotechnology. C.L. is supported by Fondazione Prossimo Mio. L.G.G. is supported by the Italian Ministry of University and Research grants PRIN-20224NMLXK, PRIN-P2022Z8HNC, Italian Ministry for University and Research Grants PE00000007 (INF-ACT), Italian Association for Cancer Research (AIRC) Grants 22737 and a donation from FONDAZIONE SAME.

---

**Table 1** List of genes differentially expressed in rVSV (logFC > 0.25) vs rLCMV (logFC < −0.25) comparison, related to Figure S2B. Absolute logFC > 0.25 and adjusted p-value <0.05 filtering was applied.

**Table 2** List of genes differentially expressed among the two rLCMV unbiased clusters: cluster 0 (logFC > 0.25) vs cluster 1 (logFC < −0.25) comparison, related to Figure 1D. Absolute logFC > 0.25 and adjusted p-value <0.05 filtering was applied.

**Table 3** List of genes upregulated in each unbiased cluster identified in the scRNA-seq experiment, related to Figure 2G. A filter of absolute logFC > 0.25 was applied.

## Supplementary figure legends

**Figure S1.**
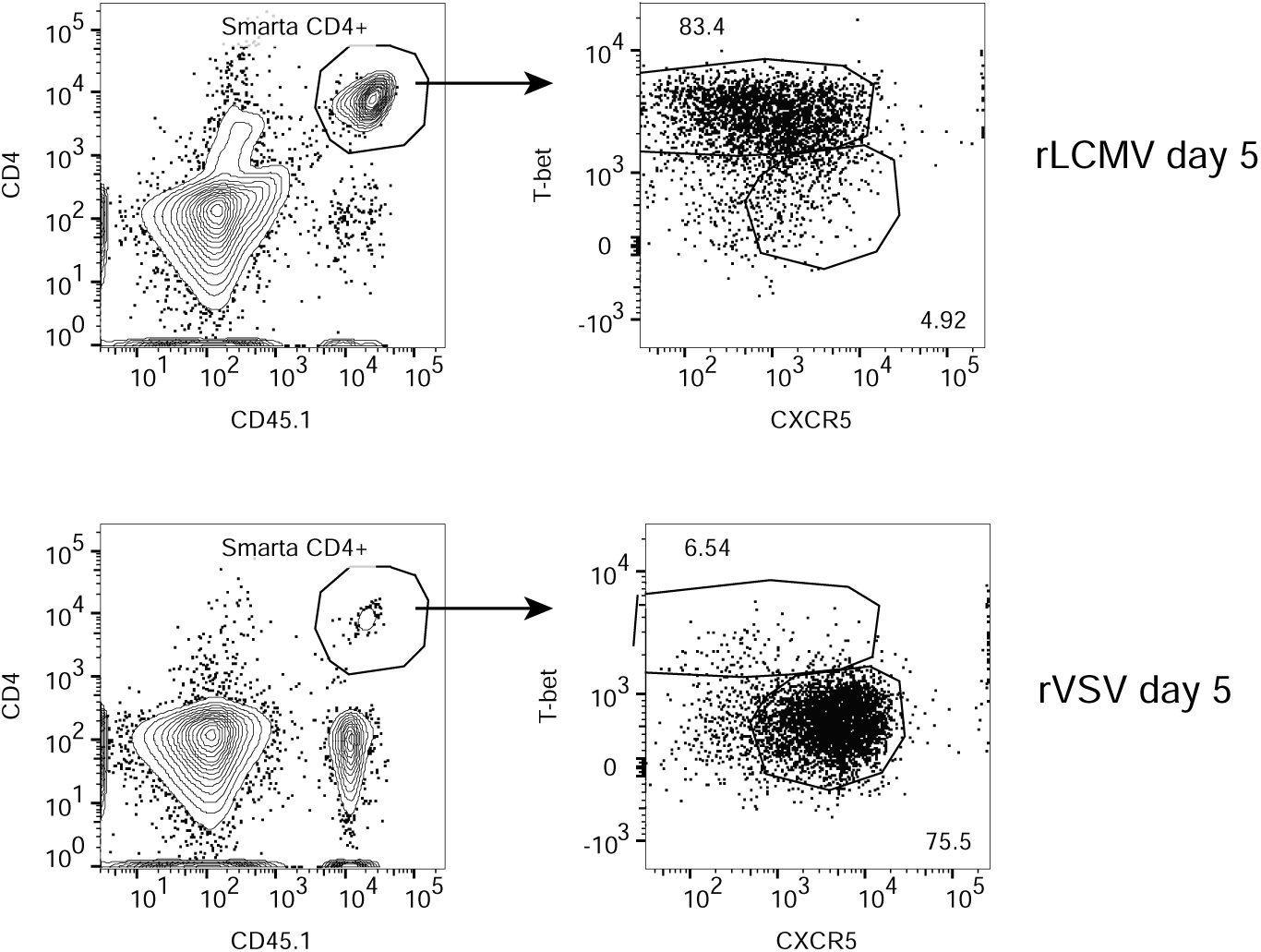
*VSV and LCMV induce distinct antiviral CD4+ T cell polarization.* 0.5*10^6^ purified CD45.1^+^ Smarta CD4+ T cells were transferred into CD45.2^+^ WT recipients 1 day before s.c. rLCMV or rVSV infection (1*10^5^ FFU /footpad). dLNs were analyzed 5 days post infection. Representative flow cytometry plots showing T_H_1 (T-bet^+^CXCR5^-^) and T_FH_ (T-bet^-^ CXCR5^+^) cells among Smarta CD4^+^ T cells in dLNs. Numbers represent the percentage of cells within the indicated gate.

**Figure S2.**
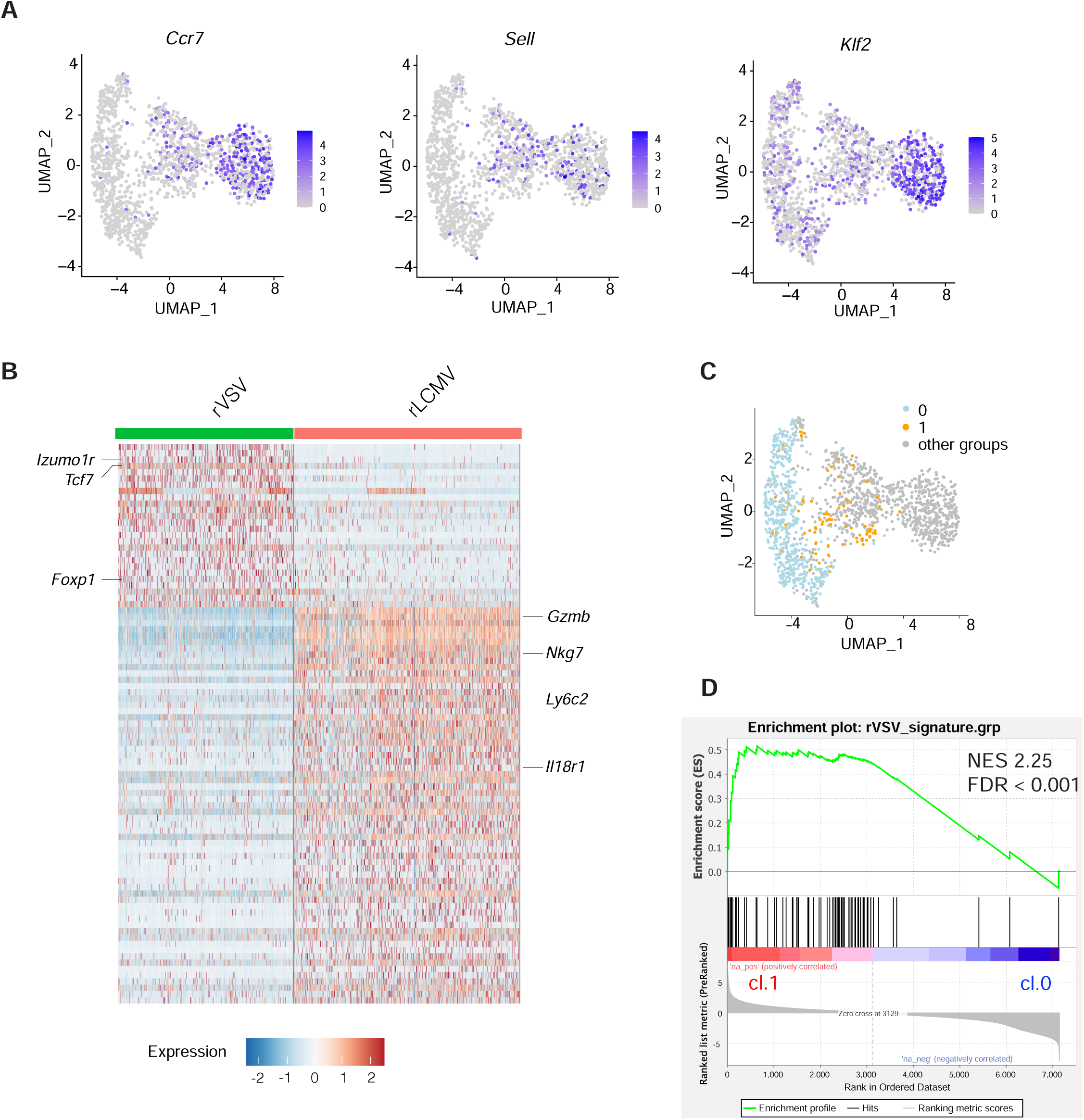
*CD4^+^ T cells from the infected hosts are clustered based on the infective agent.* **A)** Feature plot representation of the natural-log normalized expression level of *Ccr7* (left panel), *Sell* (middle panel) and *Klf2* (right panel) on the dataset in Fig. 1B. **B)** Heatmap of normalized and scaled expression values of the 561 marker genes identifying the two groups (logFC threshold: ±0.25 and adjusted p value < 0.05 filters were applied). Color coding of the bar on the top of the heatmap as in Fig. 1B. Noteworthy genes representative of each group are indicated. **C)** UMAP projection of dataset in Fig. 1B. Each dot corresponds to a single cell, colored according to the two unbiased clusters identified among the rLCMV cells (cluster 0 in light blue and cluster 1 in orange. Naïve and rVSV cells are colored in grey). **D)** Gene Set Enrichment Analysis (GSEA) relative to the rVSV signature enrichment in cluster 1 (Tcf7+) versus cluster 0 (Gzmb+) of Figure 1C. Genes were pre-ranked on the basis of the log2 fold change between the two clusters using the GseaPreranked Java tool (https://doi.org/10.1073/pnas.0506580102). The rVSV signature has been defined as the genes upregulated in rVSV compared to rLCMV in dataset in Figure 1B, with adj. pvalue < 0.05. NES: Normalized Enrichment Score.

**Figure S3.**
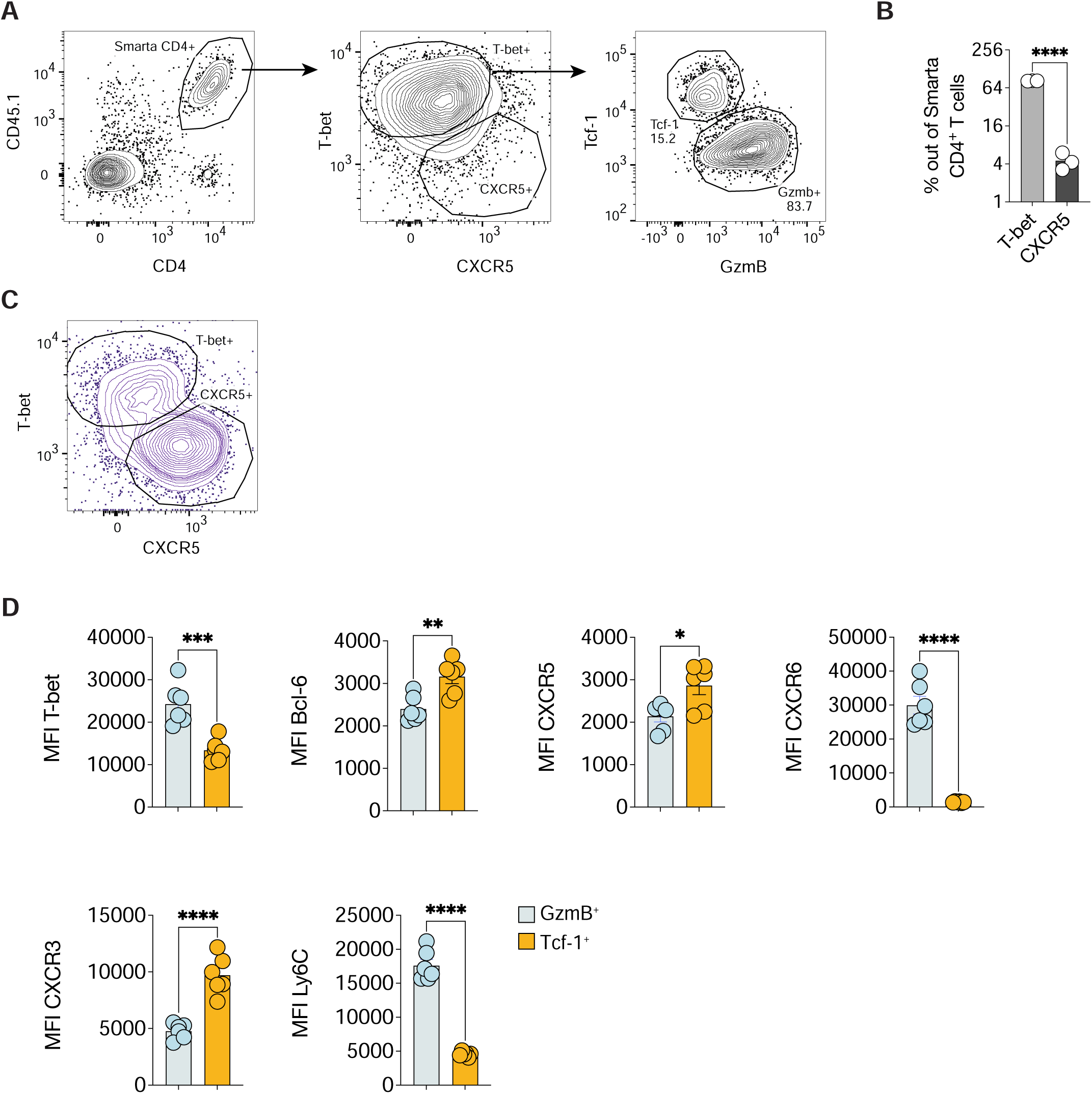
*Gating strategy of Tcf-1 and GzmB subsets among T-bet^+^ Smarta CD4^+^ T cells.* **A)** 0.5*10^6^ purified CD45.1^+^ Smarta CD4^+^ T cells were transferred into CD45.2^+^ WT recipients 1 day before s.c. rLCMV infection (1*10^5^ FFU /footpad). dLNs were analyzed 5 days post infection. The gating strategy for Fig. 1F is shown. First, Smarta CD4^+^ T were identified as CD4^+^CD45.1^+^ cells among all LN cells. Next, T-bet^+^CXCR5^-^ and T-bet^-^CXCR5^+^ among all Smarta CD4^+^ T cells are shown. Finally, Tcf-1^+^ and GzmB^+^ cells among T-bet^+^CXCR5^-^ are shown. Numbers represent the percentage of cells within the indicated gate. **B)** Quantification of T-bet^+^ and CXCR5^+^ cells expressed as percentages of Smarta CD4^+^ T cells in dLNs of mice described in A. *n=3*. Mean ± SEM is shown. Data are representative of at least five independent experiments. An unpaired two-tailed t test was applied. **** *p-value* <= 0.0001. **C)** Representative plot showing T-bet^+^CXCR5^-^ and T-bet^-^CXCR5^+^ cells among all cells of the LN, shown as controls for T-bet and CXCR5 staining. **D)** 0.5*10^6^ purified CD45.1^+^ Smarta CD4^+^ T cells were transferred into CD45.2^+^ WT recipients 1 day before s.c. rLCMV infection (1*10^5^ FFU /footpad). dLNs were analyzed 5 days post infection. Quantification of the MFI of T-bet, Bcl-6, CXCR5, CXCR6, CXCR3 and Ly6C on GzmB^+^ versus Tcf-1^+^ Smarta CD4^+^ T cells in dLNs of mice described above. *n=6.* Mean ± SEM is shown. Data are representative of at least three independent experiments. An unpaired two-tailed t test was applied. * *p value* <= 0.05, ** *p value* <= 0.01, *** *p value* <= 0.001, **** *p value* <= 0.0001.

**Figure S4.**
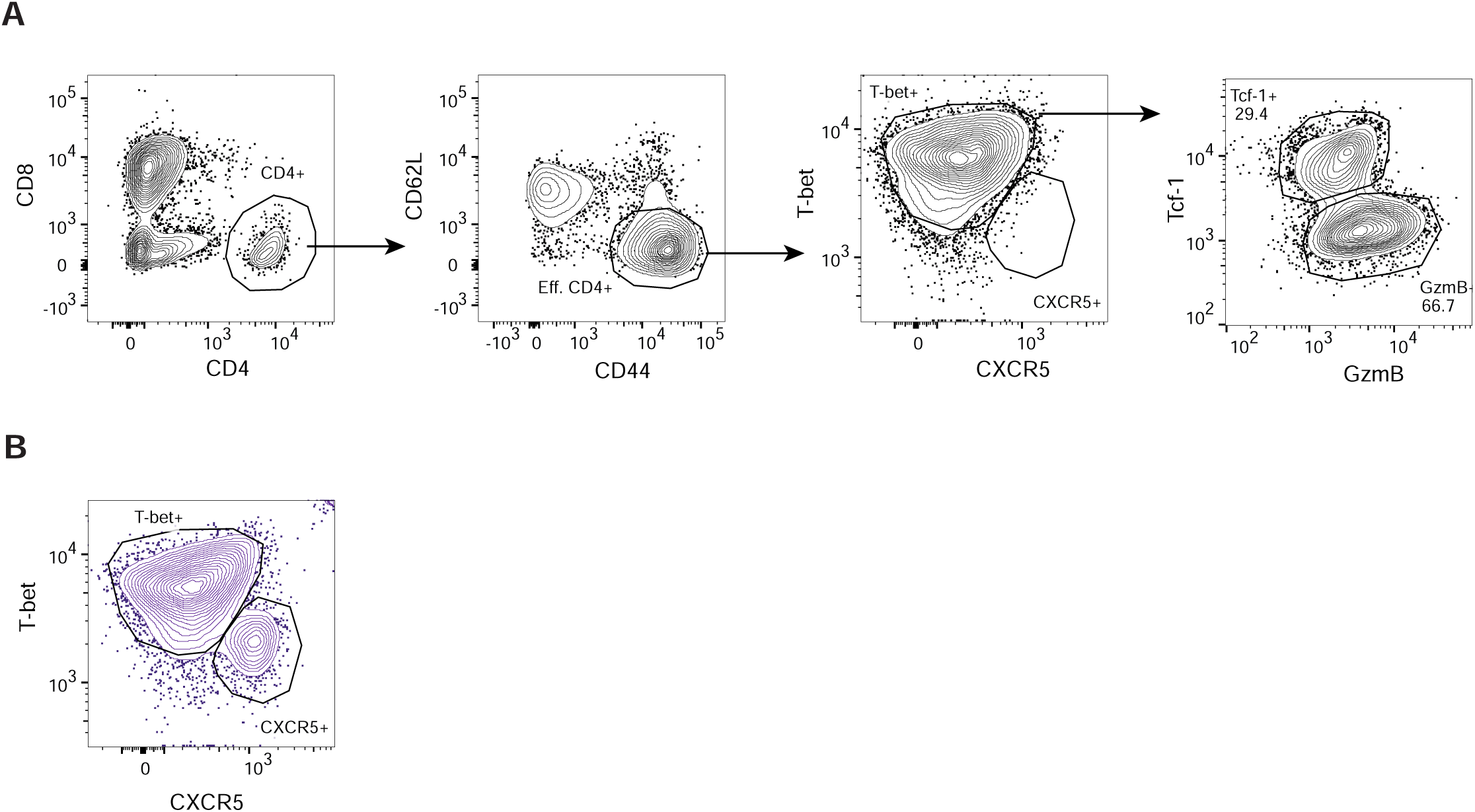
*Gating strategy of Tcf-1 and GzmB subsets among T-bet^+^ endogeneous CD4^+^ T cells.* **A)** CD45.2^+^ WT mice were infected s.c. with rLCMV (1*10^5^ FFU /footpad) and dLNs were analyzed 7 days post infection. The gating strategy for Fig. 1H is shown. First, endogenous CD4^+^ T cells were identified as CD4^+^CD8^-^ cells among all LN cells. Second, effector CD4^+^ T cells were identified as CD44^+^CD62L^-^ cells among CD4^+^ T cells. Then, T-bet^+^CXCR5^-^ and T-bet^-^CXCR5^+^ among all effector CD4^+^ T cells are shown. Finally, Tcf-1^+^ and GzmB^+^ cells among T-bet^+^CXCR5^-^ are shown. Numbers represent the percentage of cells within the indicated gate. **B)** Representative plot showing T-bet^+^CXCR5^-^ and T-bet^-^CXCR5^+^ cells among all cells of the LN, shown as controls for T-bet and CXCR5 staining.

**Figure S5.**
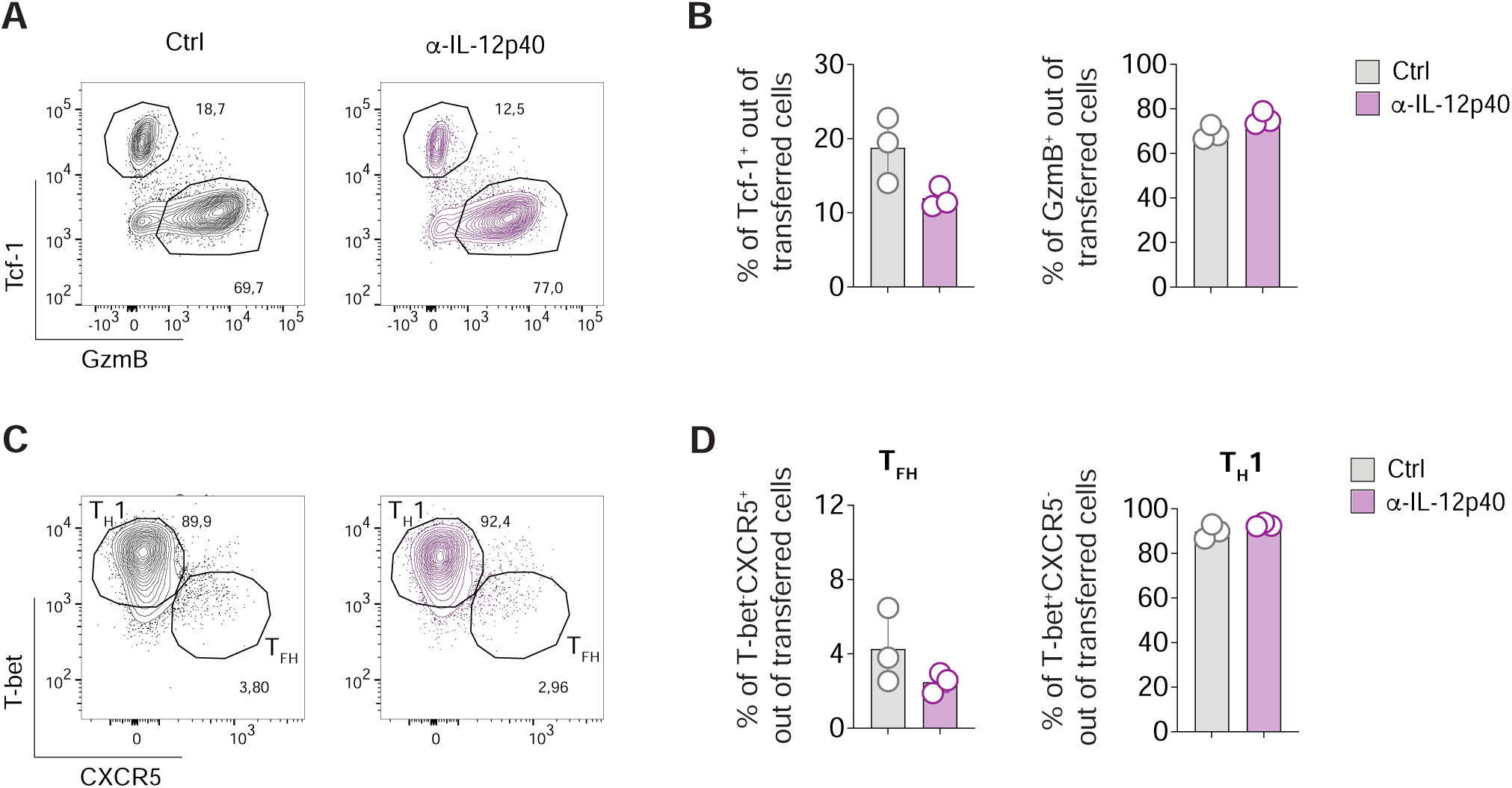
*IL-12 is not involved in CD4^+^ T cell differentiation upon LCMV infection.* **A)** 0.5*10^6^ purified CD45.1^+^ Smarta CD4^+^ T cells were transferred into CD45.2^+^ WT recipients 1 day before s.c. rLCMV infection (1*10^5^ FFU /footpad). CD45.2^+^ WT recipient mice were also treated with α-IL-12p40 blocking antibody (or Isotype Ctrl) at d0 and d3 after infection. dLNs were analyzed 5 days post infection. Representative flow cytometry plot showing Tcf-1^+^ versus GzmB^+^ cells among Smarta CD4^+^ T cells in dLNs. Numbers represent the percentage of cells within the indicated gate. **B**) Quantification of Tcf-1^+^ and GzmB^+^ cells expressed as percentages out of transferred Smarta CD4^+^ T cells in dLNs of mice described in (A). *n=3.* Mean ± SEM is shown. Data are representative of three independent experiments. An unpaired two-tailed t test was applied. Statistics is not shown since there are no statistically significant differences between conditions. **C**) Representative flow cytometry plots showing T_H_1 (T-bet^+^CXCR5^-^) versus T_FH_ (T-bet^-^CXCR5^+^) cells among Smarta CD4^+^ T cells in dLNs of mice described in (A). Numbers represent the percentage of cells within the indicated gate. **D**) Quantification of T_FH_ and T_H_1, expressed as percentages of transferred Smarta CD4^+^ T cells, in dLNs of mice described in (A). *n=3.* Mean ± SEM is shown. Data are representative of three independent experiments. An unpaired two-tailed t test was applied. Statistics is not shown since there are no statistically significant differences between conditions.

**Figure S6.**
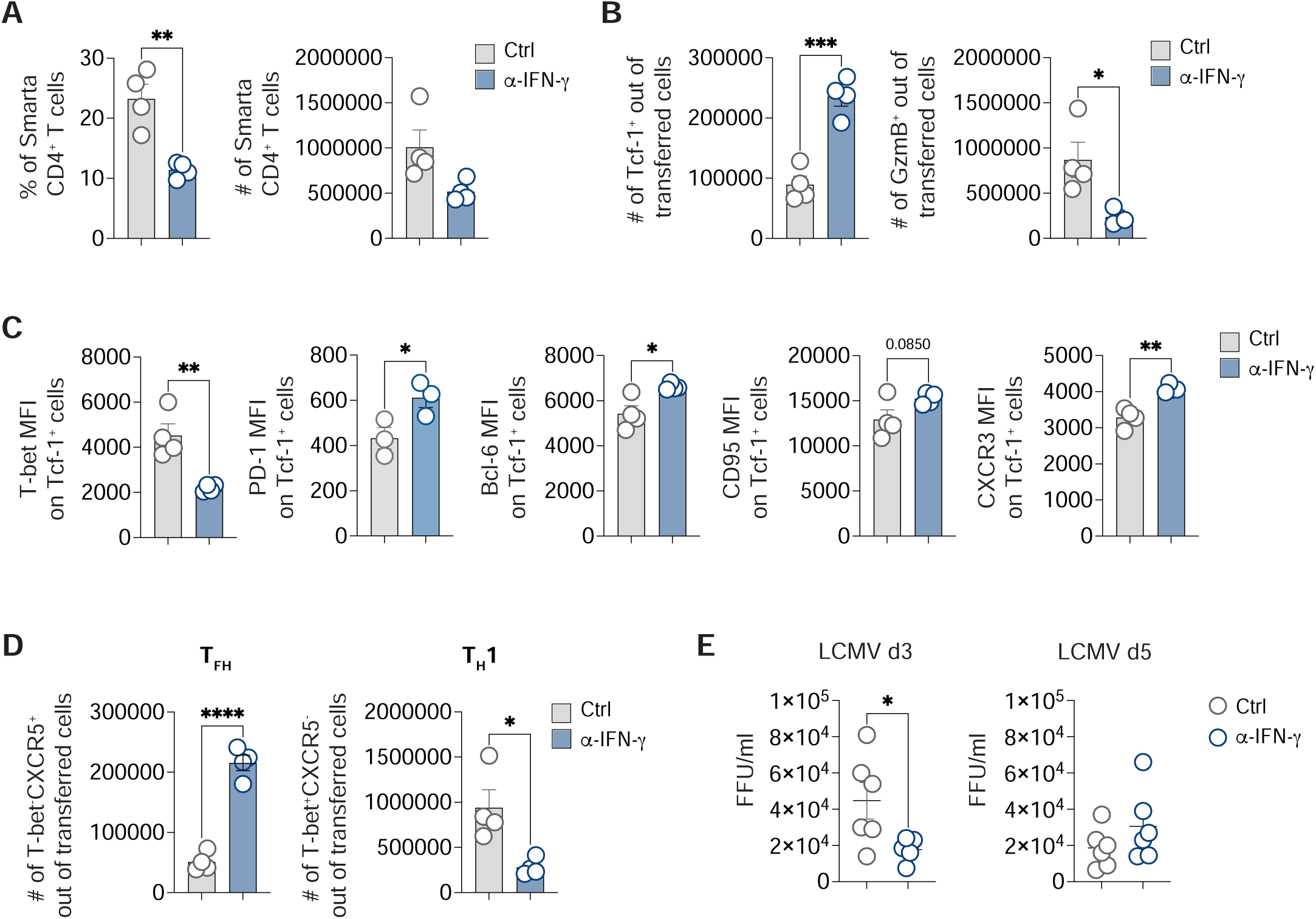
*IFN-γ suppresses T follicular helper cell differentiation.* 0.5*10^6^ purified CD45.1^+^ Smarta CD4+ T cells were transferred into CD45.2^+^ WT recipients 1 day before s.c. rLCMV infection (1*10^5^ FFU /footpad). CD45.2^+^ WT recipient mice were also treated with α-IFN-γ blocking antibody (or isotype Ctrl) at day 0. dLNs were analyzed 5 days post infection. **A)** Quantification of Smarta CD4^+^ T cells in dLNs of mice described in (A) expressed as percentages (left) and absolute numbers (right) out of total LN cells. *n=4*. Mean ± SEM is shown. Data are representative of at least three independent experiments. An unpaired two-tailed t test was applied. \**p-value* <= 0.05. **B)** Quantification of Tcf-1^+^ (left) and GzmB^+^ (right) cells in dLNs of mice described in (A) expressed as absolute numbers out of total LN cells. *n=4*. Mean ± SEM is shown. Data are representative of at least three independent experiments. An unpaired two-tailed t test was applied. \**p-value* <= 0.05, \*\*\**p-value* <= 0.001. **C)** Quantification of the MFI of T-bet, PD-1, Bcl-6, CD95, and CXCR3 on Tcf-1^+^ Smarta CD4^+^ T cells in dLNs of mice is shown. *n=3,4.* Mean ± SEM is shown. Data are representative of two independent experiments. An unpaired two-tailed t test was applied. * *p value* <= 0.05, \*\**p-value* <= 0.01. **D)** Quantification of T_FH_ (left) and T_H_1 (right), expressed as absolute numbers out of total LN cells. *n=4*. Mean ± SEM is shown. Data are representative of at least three independent experiments. An unpaired two-tailed t test was applied. \**p-value* <= 0.05, \*\*\*\**p-value* <= 0.0001.**E)** Quantification of viral titers in dLNs at day 3 and day 5 upon LCMV infection. An unpaired two-tailed t test was applied. * *p value* <= 0.05

**Figure S7.**
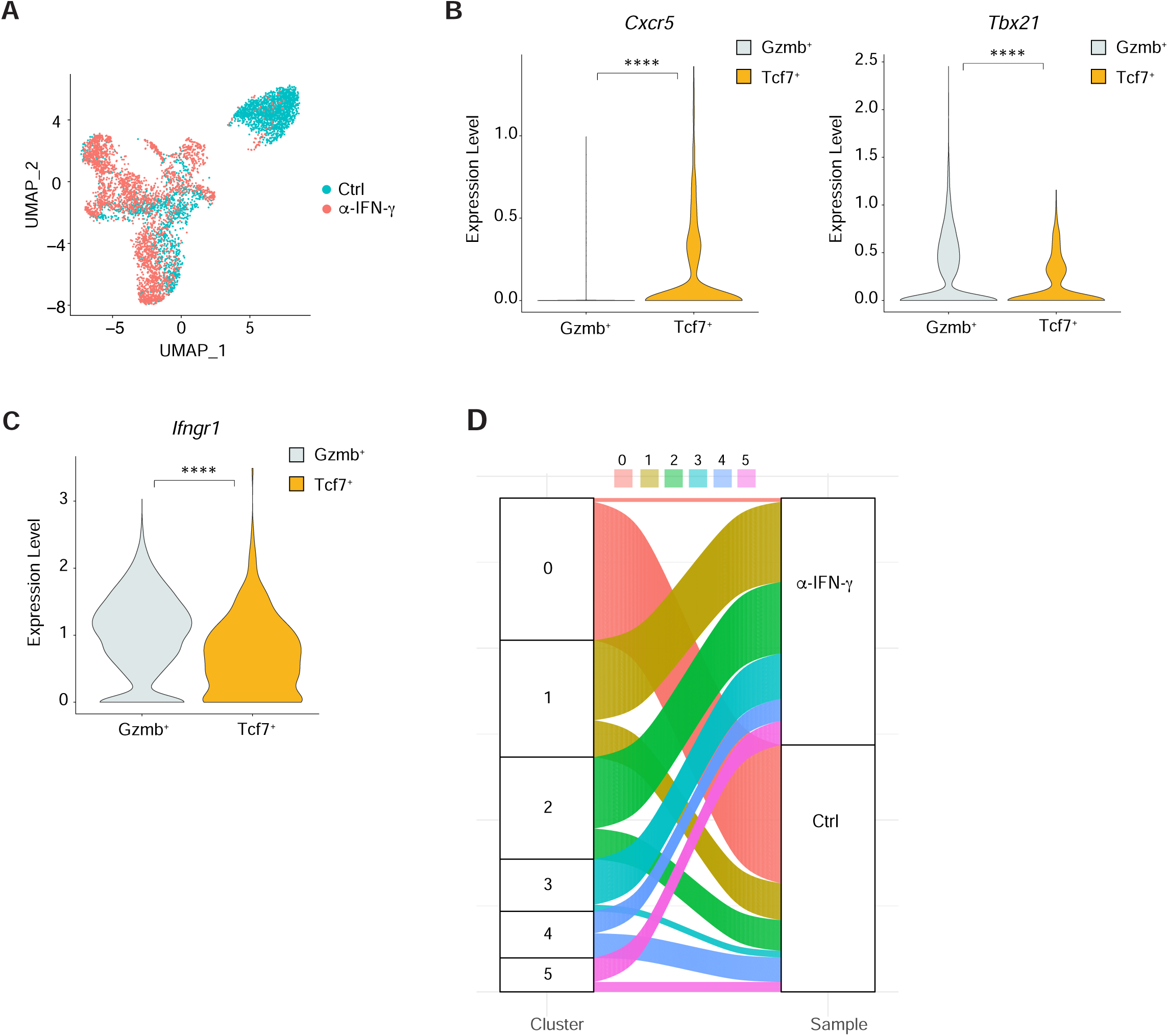
*scRNA-seq analysis of Gzmb- and Tcf7-expressing clusters.* **A)** UMAP projection of 5,746 sorted and sequenced LCMV-specific CD4^+^ T cells (dataset in Fig. 2). Each dot corresponds to a single cell, colored according to the condition (control cells in cyan, α-IFN-γ treated cells in red). The same number of cells for both samples has been considered (2,873 cells). **B)** Violin plot representation of the natural-log normalized expression level of *Cxcr5* (left panel) and *Tbx21* (right panel), comparing Gzmb^+^ and Tcf7^+^ cells belonging to dataset in A and Fig. 2. Two-tailed Mann-Whitney test has been performed, ****p < 0.0001. **C)** Violin plot representation of the natural-log normalized expression level of *Ifngr1.* Two-tailed Mann-Whitney test has been performed, ****p < 0.0001. **D)** Alluvial plot showing the composition of unbiased clusters represented in Fig. 2G according to the condition of each cell. Cluster colors as in Fig. 2G.

**Figure S8.**
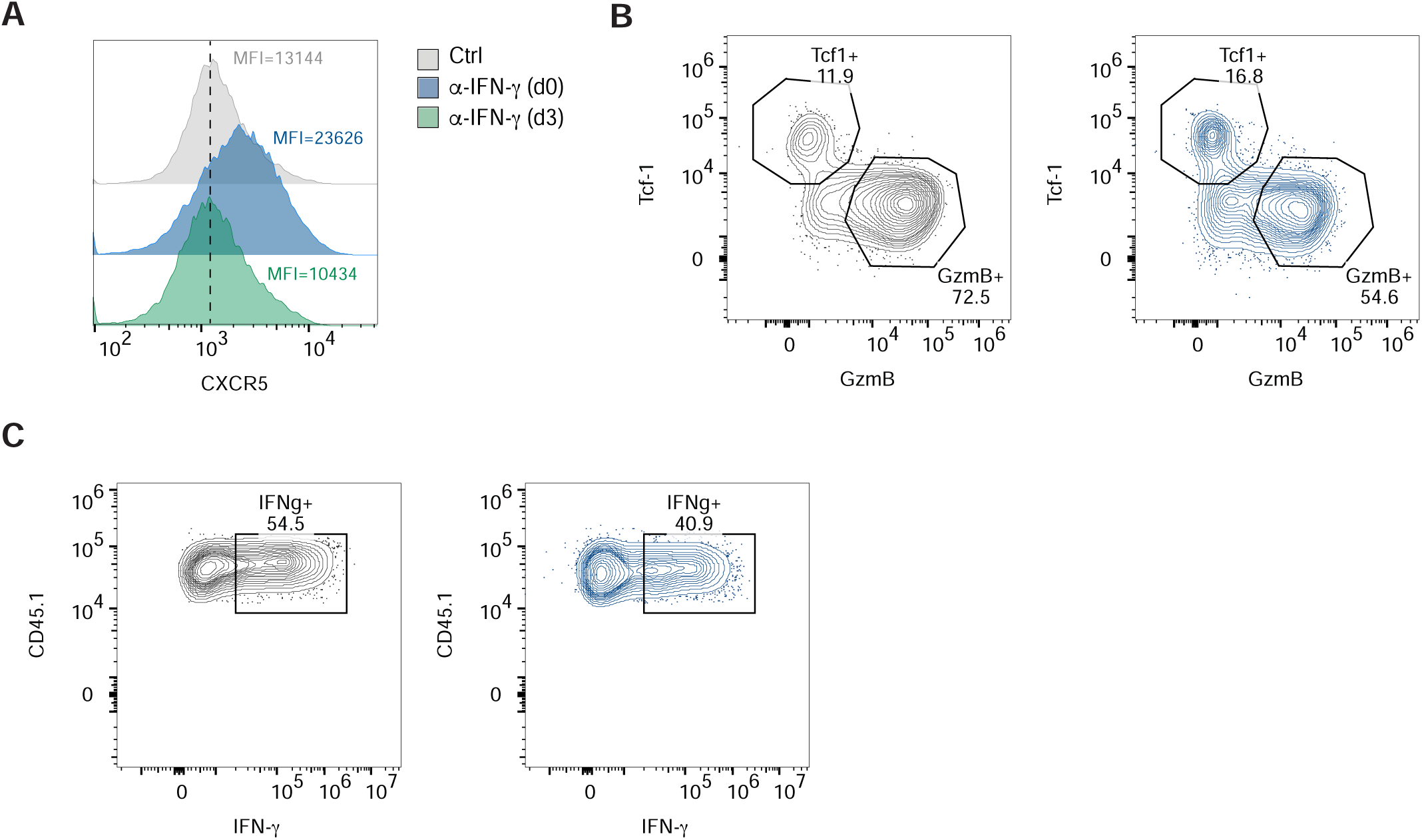
*The IFN-γ responsible for T_FH_ suppression is produced in the first days upon infection.* **A)** 0.5*10^6^ purified CD45.1^+^ Smarta CD4^+^ T cells were transferred into CD45.2^+^ WT recipients 1 day before s.c. rLCMV infection (1*10^5^ FFU /footpad). CD45.2^+^ WT recipient mice were also treated with α-IFN-γ blocking antibody (or isotype Ctrl) at day 0 or d3 after infection. dLNs were analyzed 5 days post infection. Representative plots of the MFI of CXCR5 on Tcf-1^+^ Smarta CD4^+^ T cells are shown. **F)** 0.5*10^6^ purified CD45.1^+^ Smarta CD4^+^ T cells were transferred into CD45.2^+^ WT recipients 1 day before s.c. rLCMV infection (1*10^5^ FFU /footpad). CD45.2^+^ WT recipient mice were also treated with α-IFN-γ blocking antibody (or isotype Ctrl) at day 0. dLNs were analyzed 3 days post infection. Representative plots of Tcf-1^+^ and GzmB^+^ cells out of transferred Smarta CD4^+^ T cells are shown. **C)** Representative plots of IFN-γ^+^ cells out of ex-vivo restimulated Smarta CD4^+^ T cells in dLNs of mice described in (F) are shown.

**Figure S9.**
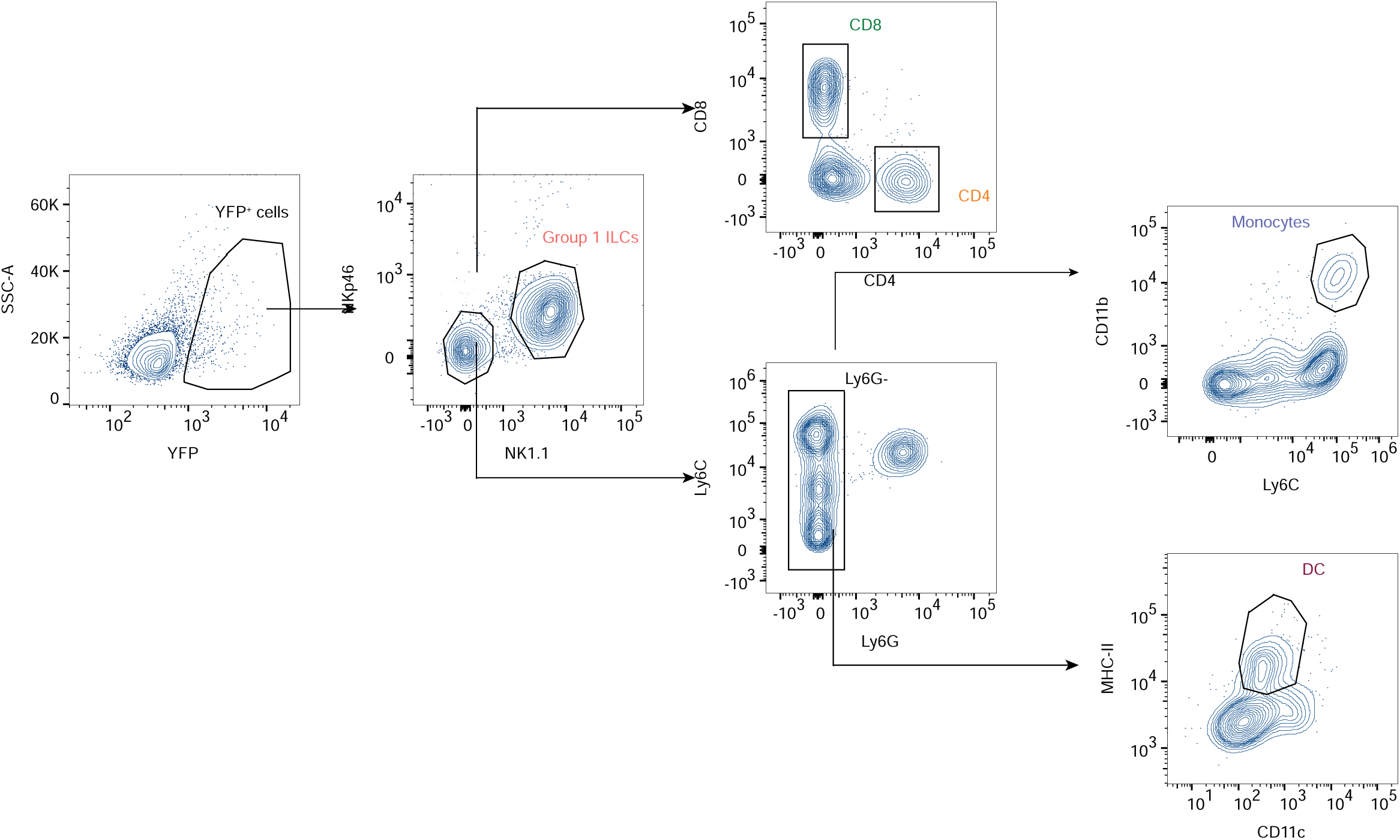
*Gating strategy for immune cell subsets analyzed for production of IFN-γ.* dLNs of IFN-γ-YFP mice were analyzed at 24h, 48h and 72 hours upon s.c. rLCMV infection (1*10^5^ FFU /footpad). The gating strategy used to identify the immune cell subsets analyzed for production of IFN-γ is shown. First, group 1 ILCs were identified as NK1.1^+^NKp46^+^ cells out of all YFP^+^ cells in the dLNs. Then, CD8^+^ and CD4^+^ T cells were identified among the NK1.1^-^ NKp46^-^ cells. Monocytes (CD11b^+^ Ly6C^hi^) and DC (CD11c^+^ MHC-II^+^) were identified out of Ly6G^-^ and NK1.1^-^NKp46^-^ cells.

**Figure S10.**
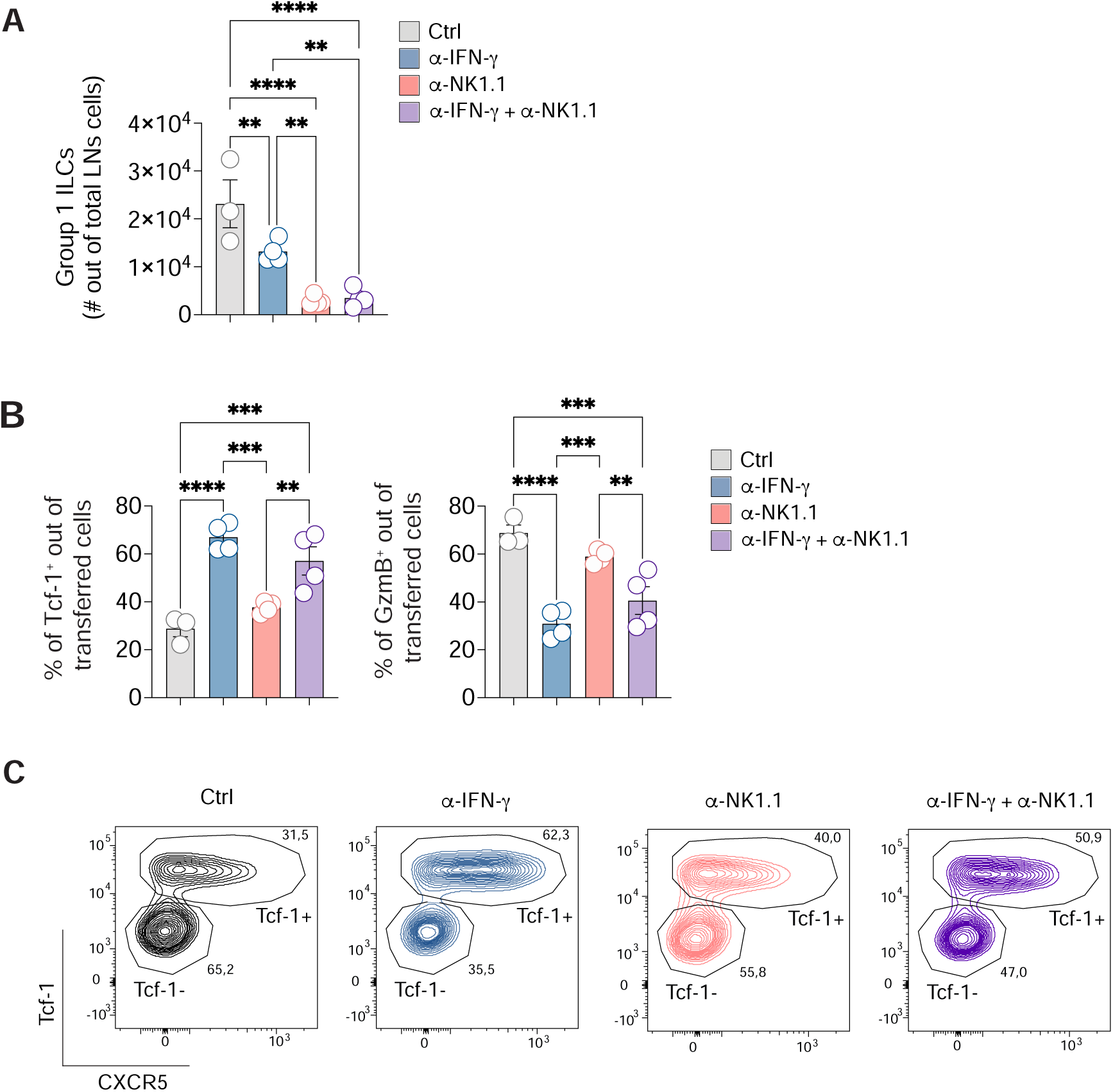
*IFN-γ derived from group 1 ILCs is not involved in CD4^+^ T cell polarization.* **A)** 0.5*10^6^ purified CD45.1^+^ Smarta CD4^+^ T cells were transferred into CD45.2^+^ WT recipients 1 day before s.c. rLCMV infection (1*10^5^ FFU /footpad). In some conditions CD45.2^+^ WT recipient mice were also treated with α-IFN-γ blocking antibody at day 0, α-NK1.1 antibody (d-1, d0) or both antibodies in combination. dLNs were analyzed 5 days post infection. Quantification of group 1 ILCs, expressed as absolute numbers in dLNs of described mice. *n=3* (Ctrl), 4 (α-IFN-γ, α-NK1.1, α-IFN-γ + α-NK1.1). Mean ± SEM is shown. Data are representative of three independent experiments. One-way ANOVA with uncorrected Fisher’s LSD was applied. ** *p value* <= 0.01, **** *p value* <= 0.0001. **B)** Quantification of Tcf-1^+^ and GzmB^+^ cells, expressed as percentages out of transferred Smarta CD4^+^ T cells, in dLNs of mice described in (A). *n=3* (Ctrl), 4 (α-IFN-γ, α-NK1.1, α-IFN-γ + α-NK1.1). Mean ± SEM is shown. Data are representative of three independent experiments. One-way ANOVA with uncorrected Fisher’s LSD was applied. ** *p value* <= 0.01, *** *p value* <= 0.001. **C)** Representative flow cytometry plots showing CXCR5 expression on Tcf-1^+^ and Tcf-1^-^ Smarta CD4^+^ T cells in dLNs of mice described in (A). Numbers represent the percentage of cells within the indicated gate.

**Figure S11.**
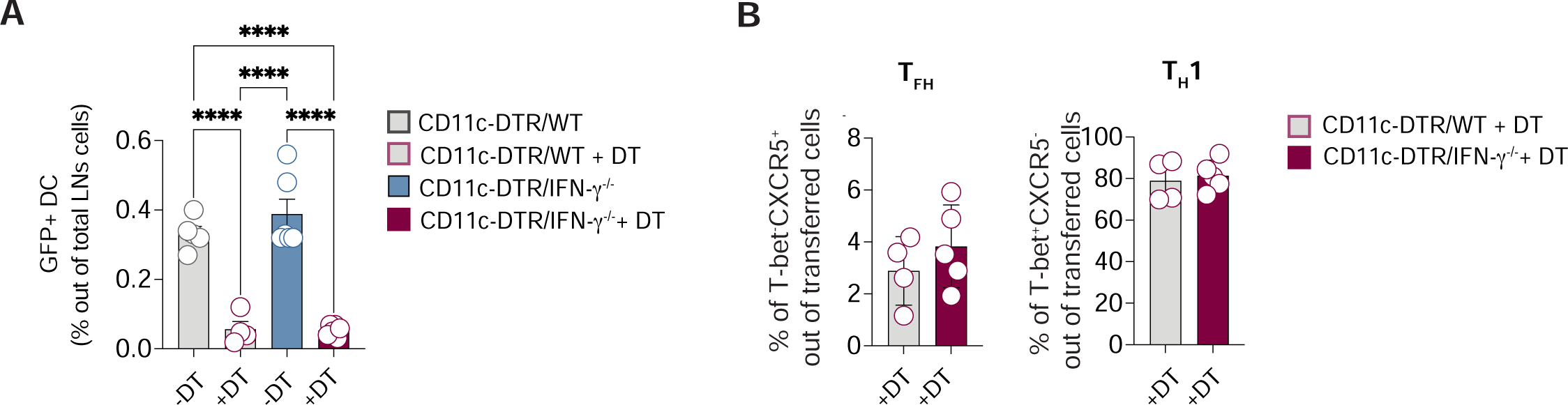
*IFN-γ derived from DCs is not involved in CD4^+^ T cell polarization* **A)** CD11c-DTR/WT or CD11c-DTR/IFN-γ^-/-^ BM chimeras were injected with PBS or DT at day-1, 1,3 and 5 and infected s.c. with LCMV (1*10^5^ FFU /footpad) at day 0. dLNs were analyzed 7 days post infection. Quantification of dendritic cells expressing GFP (CD11c^+^MHC-II^+^GFP^+^) expressed as percentages out of total LNs cells is shown. *n=*4-6. Mean ± SEM is shown. Data are representative of three independent experiments. One-way ANOVA with uncorrected Fisher’s LSD was applied. **** *p value* <= 0.0001. **C)** Quantification of T_FH_ and T_H_1, expressed as percentages out of CD44^+^CD62L^-^ effector CD4^+^ T cells, in dLNs of mice described in (A). *n=*4-6. Mean ± SEM is shown. Data are representative of three independent experiments. One-way ANOVA with uncorrected Fisher’s LSD was applied. Statistics is not shown since there are no statistically significant differences between conditions.

**Figure S12.**
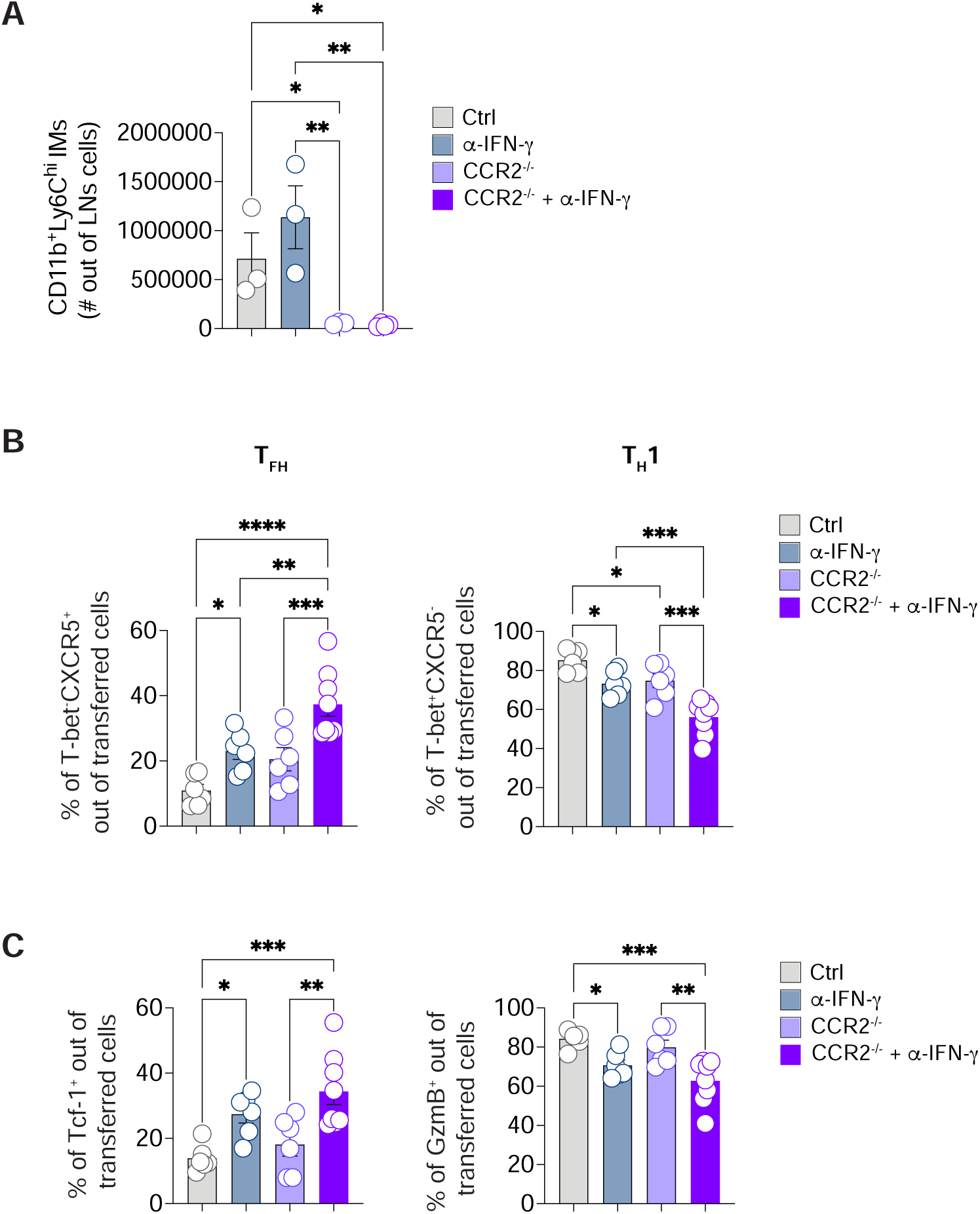
*IFN-γ derived from inflammatory monocytes is not involved in CD4^+^ T cell polarization.* **A)** 0.5*10^6^ purified CD45.1^+^ Smarta CD4^+^ T cells were transferred into CD45.2^+^ WT or CCR2^-/-^ recipients 1 day before s.c. rLCMV infection (1*10^5^ FFU /footpad). In some conditions recipient mice were also treated with α-IFN-γ blocking antibody at day 0. dLNs were analyzed 5 days post infection. Quantification of inflammatory monocytes (CD11b^+^Ly6C^+^), expressed as absolute numbers in dLNs of described mice. *n*=3-4. Mean ± SEM is shown. Data are representative of three independent experiments. One-way ANOVA with uncorrected Fisher’s LSD was applied. * *p value* <= 0.05, ** *p value* <= 0.01. **B)** Quantification of T_FH_ and T_H_1, expressed as percentages of transferred Smarta CD4^+^ T cells in dLNs. *n*=6-8. Mean ± SEM is shown. Data are representative of three independent experiments. One-way ANOVA with uncorrected Fisher’s LSD was applied. * *p value* <= 0.05, ** *p value* <= 0.01, *** *p value* <= 0.001, **** *p value* <= 0.0001. **C)** Quantification of Tcf-1^+^ and GzmB^+^ cells, expressed as percentages out of transferred Smarta CD4^+^ T cells in dLNs. *n*=6-8. Mean ± SEM is shown. Data are representative of three independent experiments. One-way ANOVA with uncorrected Fisher’s LSD was applied. * *p value* <= 0.05, ** *p value* <= 0.01, *** *p value* <= 0.001.

**Figure S13.**
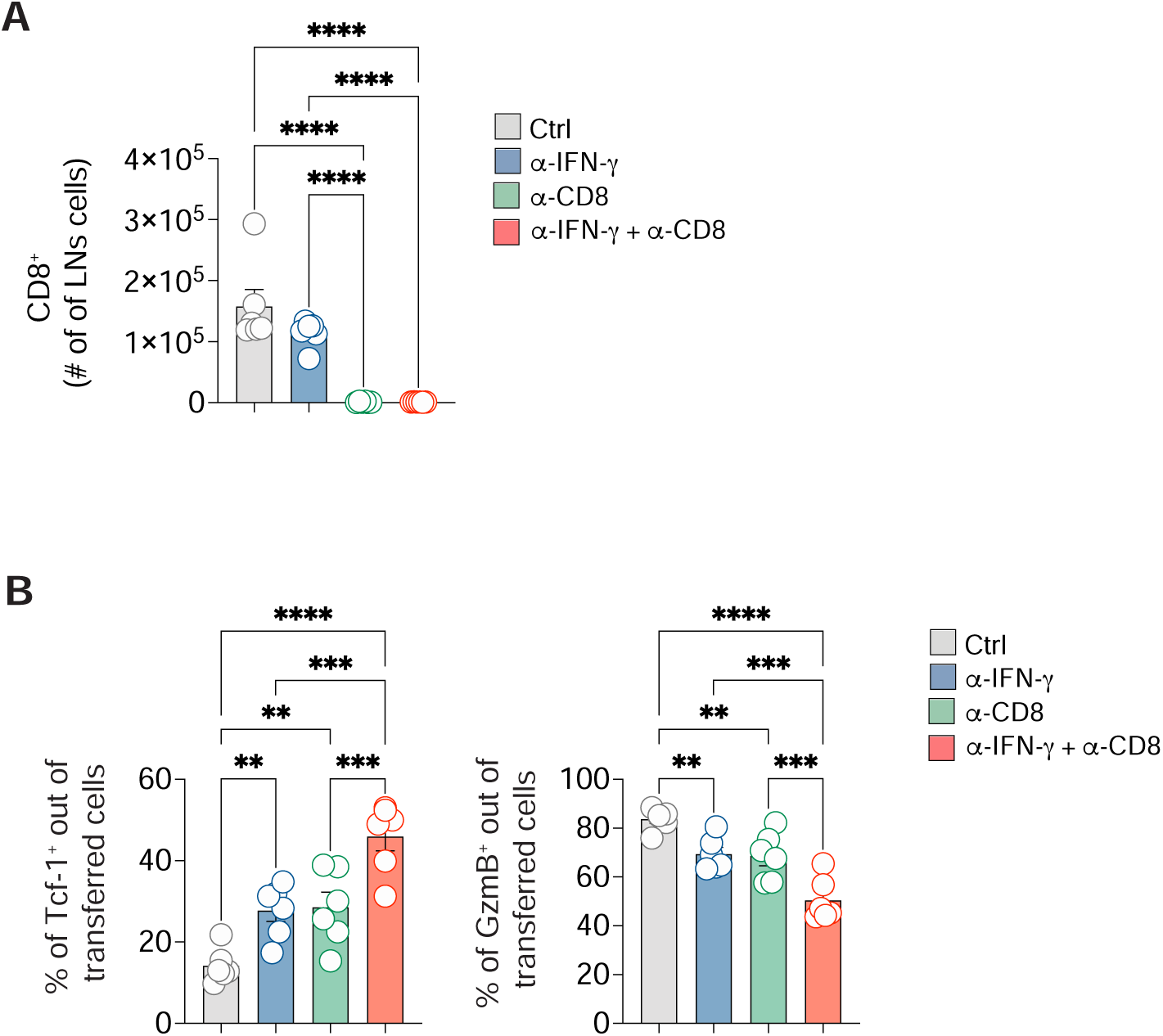
*CD8^+^ T cells contribute to CD4^+^ T cell polarization through IFN-γ and other mechanisms. **A**)* 0.5*10^6^ purified CD45.1^+^ Smarta CD4^+^ T cells were transferred into CD45.2^+^ WT recipients 1 day before s.c. rLCMV infection (1*10^5^ FFU /footpad). In some conditions CD45.2^+^ WT recipient mice were also treated with α-IFN-γ blocking antibody at day 0, α-CD8 antibody (d-1, d2) or both antibodies in combination. dLNs were analyzed 5 days post infection. Quantification of CD8^+^ T cells, expressed as absolute numbers in dLNs of described mice. *n=6*. Mean ± SEM is shown. Data are representative of two independent experiments. One-way ANOVA with uncorrected Fisher’s LSD was applied. **** *p value* <= 0.0001. **B)** Quantification of Tcf-1^+^ and GzmB^+^ cells, expressed as percentages out of transferred Smarta CD4^+^ T cells, in dLNs of mice described in (A). *n=6*. Mean ± SEM is shown. Data are representative of two independent experiments. One-way ANOVA with uncorrected Fisher’s LSD was applied. * *p value* <= 0.05, **** *p value* <= 0.0001.

**Figure S14.**
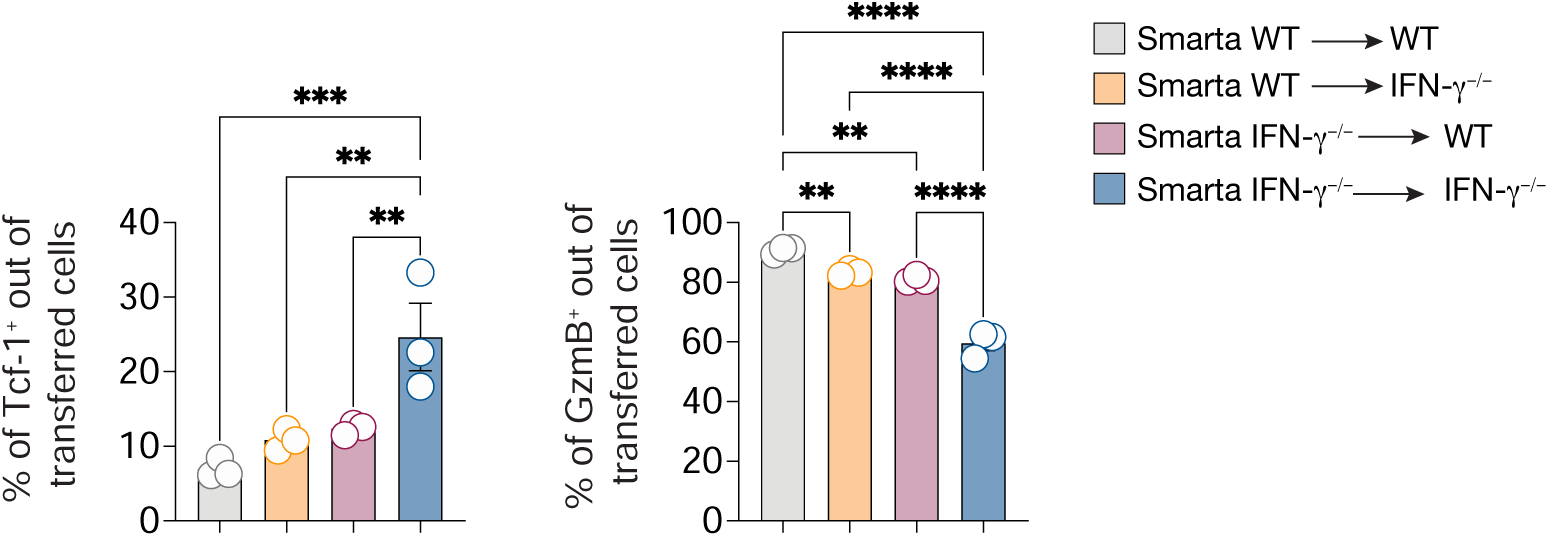
*IFN-γ from adoptively transferred CD4^+^ T cells is sufficient for T_FH_ suppression.* 0.5*10^6^ purified CD45.1^+^ Smarta CD4^+^ T cells from WT or Smarta-IFN-γ^-/-^ were transferred into CD45.2^+^ WT or IFN-γ^-/-^ recipients 1 day before s.c. rLCMV infection (1*10^5^ FFU /footpad). dLNs were analyzed 5 days post infection. Quantification of Tcf-1^+^ and GzmB^+^ cells, expressed as percentages out of transferred Smarta CD4^+^ T cells. *n=3*. Mean ± SEM is shown. Data are representative of three independent experiments. One-way ANOVA with uncorrected Fisher’s LSD was applied. ** *p value* <= 0.01, *** *p value* <= 0.001, **** *p value* <= 0.0001.

**Figure S15.**
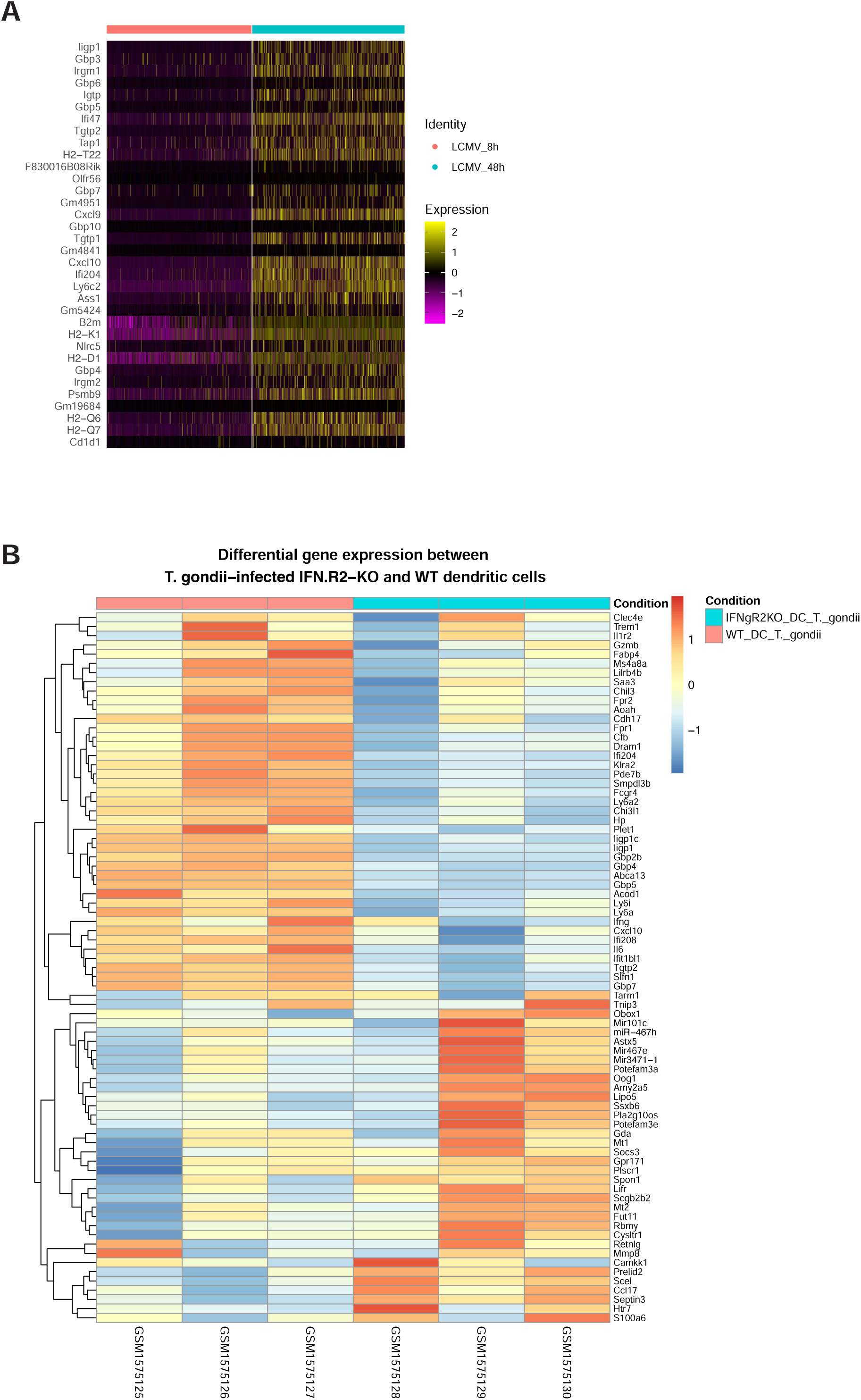
DCs sense IFN-γ *upon LCMV infection*. **A)** Heatmap of normalized and scaled expression values of the marker genes identifying the Ifng signature as reported in {Singhania et al., 2019 #10618}. DCs sorted from mice infected with LCMV for 8 or 48 hours (published dataset in {De Giovanni et al., 2020 #29499}) were compared. **B)** Heatmap visualization of differentially expressed genes comparing T. Gondii-infected conditions, with three biological replicates per condition shown in columns. The analysis includes the top 50 upregulated genes from both IFNyR2-KO infected vs. control and WT infected vs. control comparisons, showing their expression patterns in T. gondii-infected IFNyR2-KO versus T. gondii-infected wild-type samples. Expression values are represented as z-scores, where red indicates upregulation (1) and blue indicates downregulation (−1).

**Figure S16.**
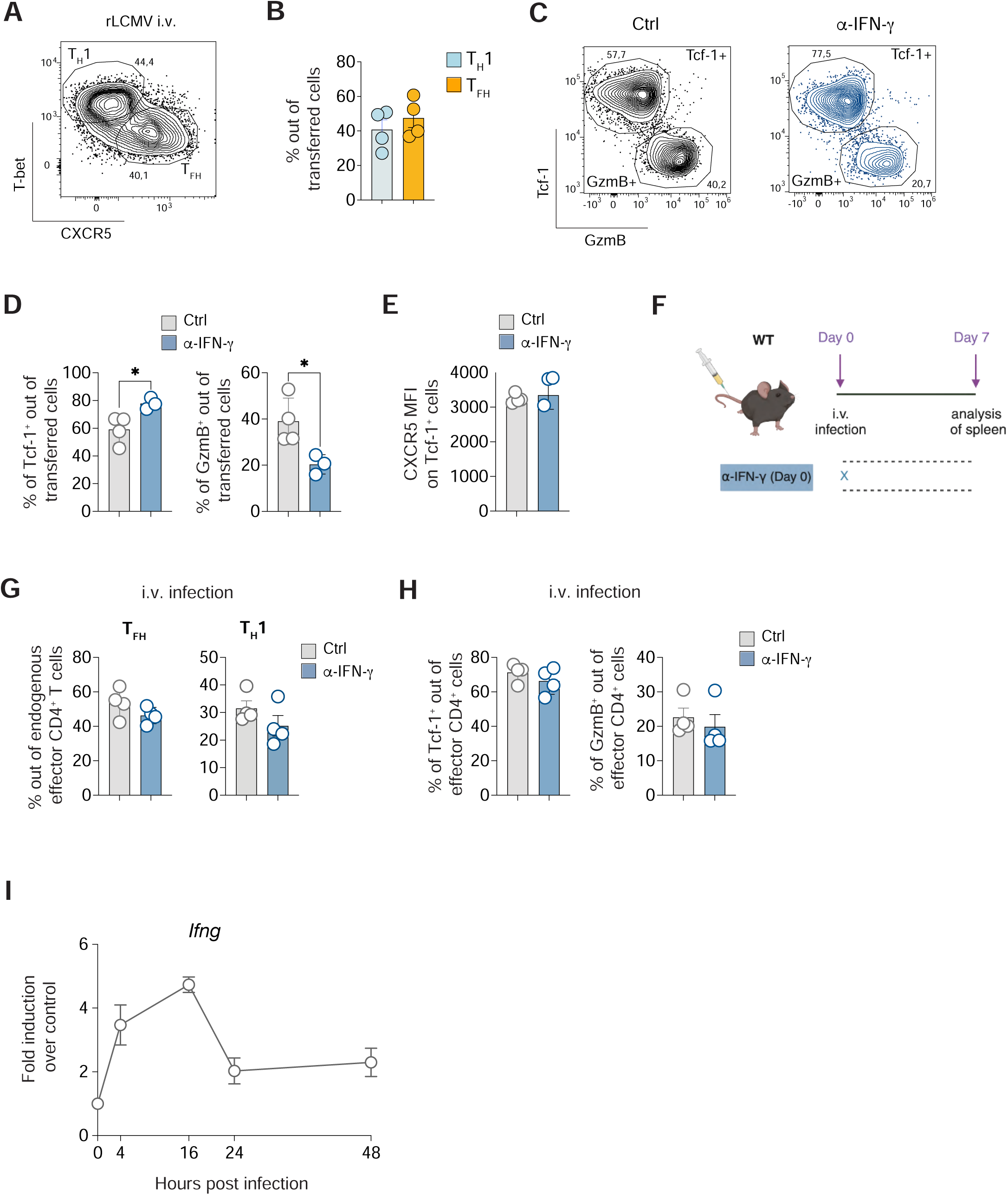
Exploring the role of IFN-γ across different routes of infection. **A)** 0.5*10^6^ purified CD45.1^+^ Smarta CD4+ T cells were transferred into CD45.2^+^ WT recipients 1 day before intravenous (i.v.) rLCMV infection (2*10^5^ FFU). CD45.2^+^ WT recipient mice were also treated with α-IFN-γ blocking antibody (or isotype Ctrl) at day 0. Spleens were analyzed 5 days post infection. Representative flow cytometry plots showing T_H_1 (T-bet^+^CXCR5^-^) and T_FH_ (T-bet^-^CXCR5^+^) cells among Smarta CD4^+^ T cells in dLNs. Numbers represent the percentage of cells within the indicated gate. **B)** Quantification of T_FH_ and T_H_1, expressed as percentages out of transferred Smarta CD4^+^ T cells in spleens of mice described in (A). *n=*4. Mean ± SEM is shown. Data are representative of three independent experiments. An unpaired two-tailed t test was applied. Statistics is not shown since there are no statistically significant differences between conditions. **C)** Representative flow cytometry plots showing Tcf-1^+^ versus GzmB^+^ cells among Smarta CD4^+^ T cells in spleens. Numbers represent the percentage of cells within the indicated gate. **D)** Quantification of Tcf-1^+^ and GzmB^+^ cells, expressed as percentages out of transferred Smarta CD4^+^ T cells in spleens of mice described in (A). *n=*4 (Ctrl), 3 (α-IFN-γ). Mean ± SEM is shown. Data are representative of three independent experiments. An unpaired two-tailed t test was applied. * *p value* <= 0.05. **E)** Quantification of the MFI of CXCR5 on Tcf-1^+^ Smarta CD4^+^ T cells in spleens of mice described in (A). *n=*4 (Ctrl), 3 (α-IFN-γ). Mean ± SEM is shown. Data are representative of three independent experiments. An unpaired two-tailed t test was applied. Statistics is not shown since there are no statistically significant differences between conditions. F) WT mice were infected i.v. (2*10^5^ FFU) with rLCMV and spleens were analyzed 7 days upon infection. WT mice were also treated with α-IFN-γ blocking antibody (or isotype Ctrl) at day 0. G) Quantification of T_FH_ (left) and T_H_1 (right), expressed as percentages out of endogenous effector CD4^+^ T cells in spleens of i.v. infected mice described in (E). *n=*4. Mean ± SEM is shown. Data are representative of three independent experiments. An unpaired two-tailed t test was applied but no statistically significant differences were detected. **H)** Quantification of Tcf-1^+^ and GzmB^+^ cells, expressed as percentages out of endogenous effector CD4^+^ T cells in spleens of i.v. infected mice. *n=*4. Mean ± SEM is shown. Data are representative of three independent experiments. An unpaired two-tailed t test was applied but no statistically significant differences were detected. **I)** WT mice were immunized s.c. with MPLA+ RBD-S1. Analysis of *Ifng* gene expression at 0, 4, 8, 16, 24 and 48 hours in dLN of immunized mice is shown. *n=6.* Mean ± SEM is shown.

**Figure S17.**
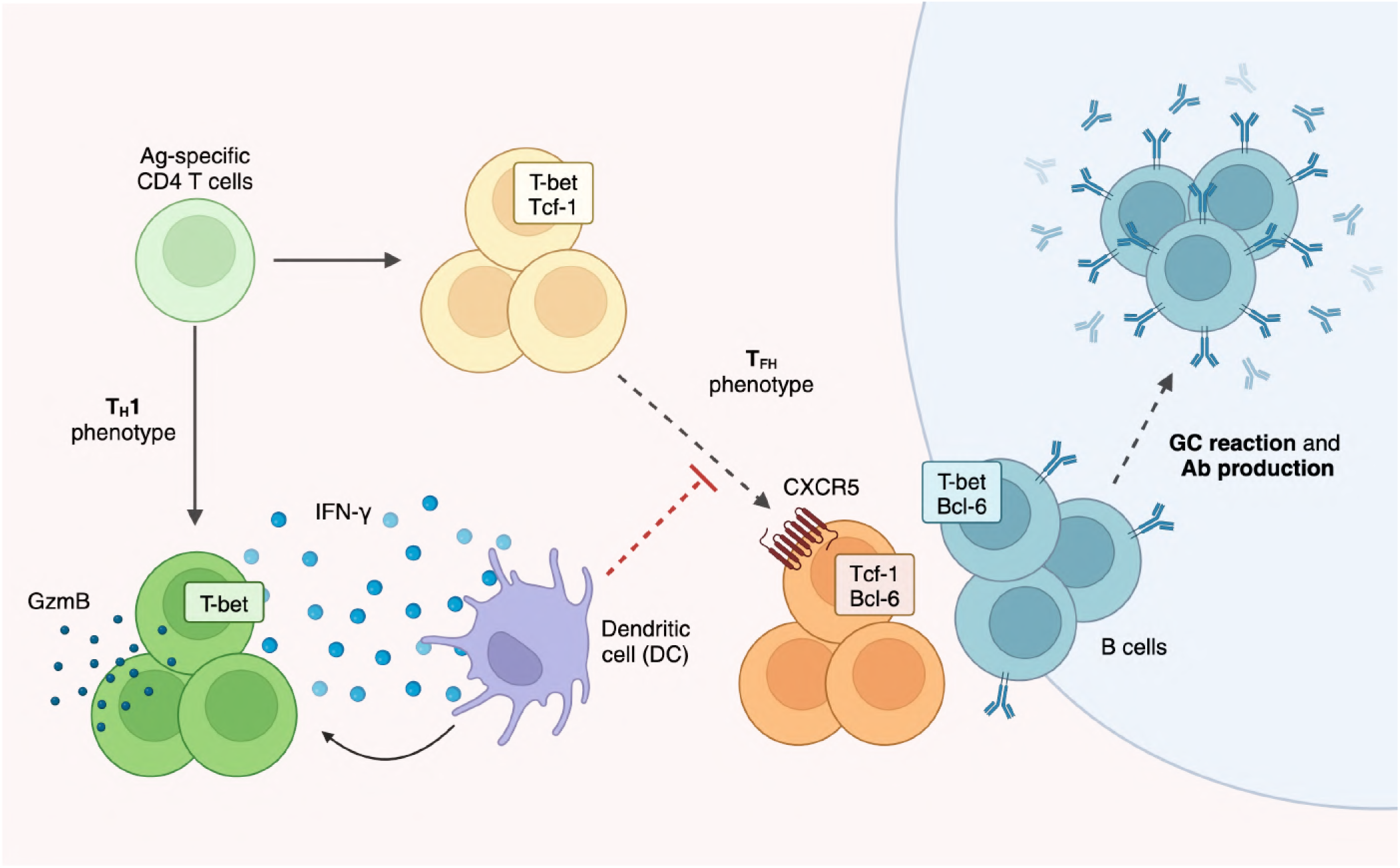
*Graphical abstract illustrating the proposed model.* Early upon subcutaneous LCMV infection Ag-specific CD4^+^ T cells are induced to differentiate into either T-bet^+^GzmB^+^ or T-bet^+^Tcf-1^+^ cells. The former subset releases high levels of IFN-γ as early as three days upon infection. IFN-γ acts to further expand the T-bet^+^GzmB^+^ subset, as well as it suppresses the development of T-bet^+^Tcf-1^+^ cells into fully differentiated CXCR5^+^Bcl-6^+^ CD4^+^ T cells. The suppression of T_FH_ cells by IFN-γ is not directly mediated through CD4^+^ T cells but rather involves another cell type, likely dendritic cells. Consistently, inhibition of IFN-γ enables robust T_FH_ differentiation, formation of germinal centers and increased antibody production.

## References

Abed NS, Chace JH, Cowdery JS (1994) T cell-independent and T cell-dependent B cell activation increases IFN-gamma R expression and renders B cells sensitive to IFN-gamma-mediated inhibition. J Immunol, 153: 3369–3377

Arroyo-Díaz NM, Bachus H, Papillion A, Randall TD, Akther J, Rosenberg AF, León B, Ballesteros-Tato A (2023) Interferon-γ production by Tfh cells is required for CXCR3^+^ pre-memory B cell differentiation and subsequent lung-resident memory B cell responses. Immunity, 56: 2358–2372.e5

Athie-Morales V, Smits HH, Cantrell DA, Hilkens CM (2004) Sustained IL-12 signaling is required for Th1 development. J Immunol, 172: 61–69

Berglund E, Maaskola J, Schultz N, Friedrich S, Marklund M, Bergenstråhle J, Tarish F, Tanoglidi A, Vickovic S, Larsson L, Salmén F, Ogris C, Wallenborg K, Lagergren J, Ståhl P, Sonnhammer E, Helleday T, Lundeberg J (2018) Spatial maps of prostate cancer transcriptomes reveal an unexplored landscape of heterogeneity. Nat Commun, 9: 2419

Bradley LM, Dalton DK, Croft M (1996) A direct role for IFN-gamma in regulation of Th1 cell development. J Immunol, 157: 1350–1358

Burton AR, Pallett LJ, McCoy LE, Suveizdyte K, Amin OE, Swadling L, Alberts E, Davidson BR, Kennedy PT, Gill US, Mauri C, Blair PA, Pelletier N, Maini MK (2018) Circulating and intrahepatic antiviral B cells are defective in hepatitis B. J Clin Invest, 128: 4588–4603

Chen JS, Chow RD, Song E, Mao T, Israelow B, Kamath K, Bozekowski J, Haynes WA, Filler RB, Menasche BL, Wei J, Alfajaro MM, Song W, Peng L, Carter L, Weinstein JS, Gowthaman U, Chen S, Craft J, Shon JC, Iwasaki A, Wilen CB, Eisenbarth SC (2022) High-affinity, neutralizing antibodies to SARS-CoV-2 can be made without T follicular helper cells. Sci Immunol, 7: eabl5652

Chodisetti SB, Fike AJ, Domeier PP, Singh H, Choi NM, Corradetti C, Kawasawa YI, Cooper TK, Caricchio R, Rahman ZSM (2020) Type II but Not Type I IFN Signaling Is Indispensable for TLR7-Promoted Development of Autoreactive B Cells and Systemic Autoimmunity. J Immunol, 204: 796–809

Choi YS, Kageyama R, Eto D, Escobar TC, Johnston RJ, Monticelli L, Lao C, Crotty S (2011) ICOS receptor instructs T follicular helper cell versus effector cell differentiation via induction of the transcriptional repressor Bcl6. Immunity, 34: 932–946

Cousens LP, Orange JS, Su HC, Biron CA (1997a) Interferon-α/β inhibition of interleukin 12 and interferon-γ production in vitro and endogenously during viral infection. Proceedings of the National Academy of Sciences, 94: 634–639

Cousens LP, Orange JS, Su HC, Biron CA (1997b) Interferon-α/β inhibition of interleukin 12 and interferon-γ production in vitro and endogenously during viral infection. Proceedings of the National Academy of Sciences, 94: 634–639

Crotty S (2011) Follicular helper CD4 T cells (TFH). Annu Rev Immunol, 29: 621–663

De Giovanni M, Cutillo V, Giladi A, Sala E, Maganuco CG, Medaglia C, Di Lucia P, Bono E, Cristofani C, Consolo E, Giustini L, Fiore A, Eickhoff S, Kastenmüller W, Amit I, Kuka M, Iannacone M (2020) Spatiotemporal regulation of type I interferon expression determines the antiviral polarization of CD4^+^ T cells. Nat Immunol, 21: 321–330

DiToro D, Winstead CJ, Pham D, Witte S, Andargachew R, Singer JR, Wilson CG, Zindl CL, Luther RJ, Silberger DJ, Weaver BT, Kolawole EM, Martinez RJ, Turner H, Hatton RD, Moon JJ, Way SS, Evavold BD, Weaver CT (2018) Differential IL-2 expression defines developmental fates of follicular versus nonfollicular helper T cells. Science, 361: eaao2933

Eisenbarth SC (2018) Dendritic cell subsets in T cell programming: location dictates function. Nat Rev Immunol,

Elsner RA, Smita S, Shlomchik MJ (2024) IL-12 induces a B cell-intrinsic IL-12/IFNγ feed-forward loop promoting extrafollicular B cell responses. Nature Immunology, 25: 1283–1295

Fallet B, Narr K, Ertuna YI, Remy M, Sommerstein R, Cornille K, Kreutzfeldt M, Page N, Zimmer G, Geier F (2016) Interferon-driven deletion of antiviral B cells at the onset of chronic infection. Science immunology, 1: eaah6817

Fiore A, Sala E, Laura C, Riba M, Nelli M, Fumagalli V, Oberrauch F, Mangione M, Cristofani C, Provero P, Iannacone M, Kuka M (2023) A fluorescent reporter model for the visualization and characterization of TDC. *Eur J Immunol*, e2350529

Fumagalli V, Ravà M, Marotta D, Di Lucia P, Bono EB, Giustini L, De Leo F, Casalgrandi M, Monteleone E, Mouro V, Malpighi C, Perucchini C, Grillo M, De Palma S, Donnici L, Marchese S, Conti M, Muramatsu H, Perlman S, Pardi N, Kuka M, De Francesco R, Bianchi ME, Guidotti LG, Iannacone M (2024) Antibody-independent protection against heterologous SARS-CoV-2 challenge conferred by prior infection or vaccination. Nat Immunol,

Fumagalli V, Ravà M, Marotta D, Di Lucia P, Laura C, Sala E, Grillo M, Bono E, Giustini L, Perucchini C, Mainetti M, Sessa A, Garcia-Manteiga JM, Donnici L, Manganaro L, Delbue S, Broccoli V, De Francesco R, D’Adamo P, Kuka M, Guidotti LG, Iannacone M (2021) Administration of aerosolized SARS-CoV-2 to K18-hACE2 mice uncouples respiratory infection from fatal neuroinvasion. *Sci Immunol*, eabl9929

Hale JS, Youngblood B, Latner DR, Mohammed AUR, Ye L, Akondy RS, Wu T, Iyer SS, Ahmed R (2013) Distinct memory CD4+ T cells with commitment to T follicular helper-and T helper 1-cell lineages are generated after acute viral infection. Immunity, 38: 805–817

Hangartner L, Zinkernagel RM, Hengartner H (2006) Antiviral antibody responses: the two extremes of a wide spectrum. Nat Rev Immunol, 6: 231–243

Hansen DS, Obeng-Adjei N, Ly A, Ioannidis LJ, Crompton PD (2017) Emerging concepts in T follicular helper cell responses to malaria. Int J Parasitol, 47: 105–110

Heufler C, Koch F, Stanzl U, Topar G, Wysocka M, Trinchieri G, Enk A, Steinman RM, Romani N, Schuler G (1996) Interleukin-12 is produced by dendritic cells and mediates T helper 1 development as well as interferon-gamma production by T helper 1 cells. Eur J Immunol, 26: 659– 668

Hsieh CS, Macatonia SE, Tripp CS, Wolf SF, O’Garra A, Murphy KM (1993) Development of TH1 CD4+ T cells through IL-12 produced by Listeria-induced macrophages. Science, 260: 547–549

Jaitin DA, Kenigsberg E, Keren-Shaul H, Elefant N, Paul F, Zaretsky I, Mildner A, Cohen N, Jung S, Tanay A, Amit I (2014) Massively parallel single-cell RNA-seq for marker-free decomposition of tissues into cell types. Science, 343: 776–779

Johnston RJ, Poholek AC, DiToro D, Yusuf I, Eto D, Barnett B, Dent AL, Craft J, Crotty S (2009) Bcl6 and Blimp-1 are reciprocal and antagonistic regulators of T follicular helper cell differentiation. Science, 325: 1006–1010

Jung S, Unutmaz D, Wong P, Sano G, De los Santos K, Sparwasser T, Wu S, Vuthoori S, Ko K, Zavala F, Pamer EG, Littman DR, Lang RA (2002) In vivo depletion of CD11c+ dendritic cells abrogates priming of CD8+ T cells by exogenous cell-associated antigens. Immunity, 17: 211–220

Komai-Koma M, Ji Y, Cao H, Liu Z, McSharry C, Xu D (2021) Monophosphoryl lipid A directly regulates Th1 cytokine production in human CD4+ T-cells through Toll-like receptor 2 and 4. Immunobiology, 226: 152132

Krueger PD, Goldberg MF, Hong SW, Osum KC, Langlois RA, Kotov DI, Dileepan T, Jenkins MK (2021) Two sequential activation modules control the differentiation of protective T helper-1 (Th1) cells. Immunity, 54: 687–701.e4

Kuka M, Iannacone M (2021) Heterogeneity in antiviral B cell responses: Lessons from the movies. Immunol Rev,

Kuka M, Munitic I, Ashwell JD (2012) Identification and characterization of polyclonal αβ-T cells with dendritic cell properties. Nature communications, 3: 1223

Lee H-M, Fleige A, Forman R, Cho S, Khan AA, Lin L-L, Nguyen DT, O’Hara-Hall A, Yin Z, Hunter CA, Muller W, Lu L-F (2015) IFNγ signaling endows DCs with the capacity to control type I inflammation during parasitic infection through promoting T-bet+ regulatory T cells. PLoS Pathog, 11: e1004635

Lighvani AA, Frucht DM, Jankovic D, Yamane H, Aliberti J, Hissong BD, Nguyen BV, Gadina M, Sher A, Paul WE, O’Shea JJ (2001) T-bet is rapidly induced by interferon-gamma in lymphoid and myeloid cells. Proc Natl Acad Sci U S A, 98: 15137–15142

Lönnberg T, Svensson V, James KR, Fernandez-Ruiz D, Sebina I, Montandon R, Soon MSF, Fogg LG, Nair AS, Liligeto U, Stubbington MJT, Ly L-H, Bagger FO, Zwiessele M, Lawrence ND, Souza-Fonseca-Guimaraes F, Bunn PT, Engwerda CR, Heath WR, Billker O, Stegle O, Haque A, Teichmann SA (2017) Single-cell RNA-seq and computational analysis using temporal mixture modelling resolves Th1/Tfh fate bifurcation in malaria. Sci Immunol, 2: eaal2192

Mata-Haro V, Cekic C, Martin M, Chilton PM, Casella CR, Mitchell TC (2007) The vaccine adjuvant monophosphoryl lipid A as a TRIF-biased agonist of TLR4. Science, 316: 1628–1632

McInnes L, Healy J, Melville J (2018) Umap: Uniform manifold approximation and projection for dimension reduction. arXiv preprint arXiv:180203426,

Mempel TR, Henrickson SE, Von Andrian UH (2004) T-cell priming by dendritic cells in lymph nodes occurs in three distinct phases. Nature, 427: 154–159

Mendoza A, Yewdell WT, Hoyos B, Schizas M, Bou-Puerto R, Michaels AJ, Brown CC, Chaudhuri J, Rudensky AY (2021) Assembly of a spatial circuit of T-bet–expressing T and B lymphocytes is required for antiviral humoral immunity. Science immunology, 6: eabi4710

Miro F, Nobile C, Blanchard N, Lind M, Filipe-Santos O, Fieschi C, Chapgier A, Vogt G, de Beaucoudrey L, Kumararatne DS, Le Deist F, Casanova JL, Amigorena S, Hivroz C (2006) T cell-dependent activation of dendritic cells requires IL-12 and IFN-gamma signaling in T cells. J Immunol, 177: 3625–3634

Miyagi T, Gil MP, Wang X, Louten J, Chu W-M, Biron CA (2007) High basal STAT4 balanced by STAT1 induction to control type 1 interferon effects in natural killer cells. The Journal of experimental medicine, 204: 2383–2396

Myles A, Gearhart PJ, Cancro MP (2017) Signals that drive T-bet expression in B cells. Cell Immunol, 321: 3–7

Nakayamada S, Kanno Y, Takahashi H, Jankovic D, Lu KT, Johnson TA, Sun HW, Vahedi G, Hakim O, Handon R, Schwartzberg PL, Hager GL, O’Shea JJ (2011) Early Th1 cell differentiation is marked by a Tfh cell-like transition. Immunity, 35: 919–931

Nguyen KB, Watford WT, Salomon R, Hofmann SR, Pien GC, Morinobu A, Gadina M, O’Shea JJ, Biron CA (2002) Critical role for STAT4 activation by type 1 interferons in the interferon-gamma response to viral infection. Science, 297: 2063–2066

Nguyen KB, Cousens LP, Doughty LA, Pien GC, Durbin JE, Biron CA (2000) Interferon α/β-mediated inhibition and promotion of interferon γ: STAT1 resolves a paradox. Nature immunology, 1: 70–76

Obeng-Adjei N, Portugal S, Holla P, Li S, Sohn H, Ambegaonkar A, Skinner J, Bowyer G, Doumbo OK, Traore B, Pierce SK, Crompton PD (2017) Malaria-induced interferon-γ drives the expansion of Tbethi atypical memory B cells. PLoS Pathog, 13: e1006576

Obeng-Adjei N, Portugal S, Tran TM, Yazew TB, Skinner J, Li S, Jain A, Felgner PL, Doumbo OK, Kayentao K, Ongoiba A, Traore B, Crompton PD (2015) Circulating Th1-Cell-type Tfh Cells that Exhibit Impaired B Cell Help Are Preferentially Activated during Acute Malaria in Children. Cell Rep, 13: 425–439

Oh SA, Seki A, Rutz S (2019) Ribonucleoprotein Transfection for CRISPR/Cas9-Mediated Gene Knockout in Primary T Cells. Curr Protoc Immunol, 124: e69

Oxenius A, Bachmann MF, Zinkernagel RM, Hengartner H (1998) Virus-specific major MHC class II-restricted TCR-transgenic mice: effects on humoral and cellular immune responses after viral infection. European journal of immunology, 28: 390–400

Oxenius A, Karrer U, Zinkernagel RM, Hengartner H (1999) IL-12 is not required for induction of type 1 cytokine responses in viral infections. The Journal of Immunology, 162: 965–973

Pien GC, Biron CA (2000) Compartmental differences in NK cell responsiveness to IL-12 during lymphocytic choriomeningitis virus infection. The Journal of Immunology, 164: 994–1001

Ray JP, Marshall HD, Laidlaw BJ, Staron MM, Kaech SM, Craft J (2014) Transcription factor STAT3 and type I interferons are corepressive insulators for differentiation of follicular helper and T helper 1 cells. Immunity, 40: 367–377

Reinhardt RL, Liang H-E, Locksley RM (2009) Cytokine-secreting follicular T cells shape the antibody repertoire. Nature immunology, 10: 385–393

Rubtsov AV, Marrack P, Rubtsova K (2017) T-bet expressing B cells - Novel target for autoimmune therapies. Cell Immunol, 321: 35–39

Rubtsova K, Rubtsov AV, Thurman JM, Mennona JM, Kappler JW, Marrack P (2017) B cells expressing the transcription factor T-bet drive lupus-like autoimmunity. J Clin Invest, 127: 1392–1404

Ryg-Cornejo V, Ioannidis LJ, Ly A, Chiu CY, Tellier J, Hill DL, Preston SP, Pellegrini M, Yu D, Nutt SL, Kallies A, Hansen DS (2016) Severe Malaria Infections Impair Germinal Center Responses by Inhibiting T Follicular Helper Cell Differentiation. Cell Rep, 14: 68–81

Sammicheli S, Kuka M, Di Lucia P, de Oya NJ, De Giovanni M, Fioravanti J, Cristofani C, Maganuco CG, Fallet B, Ganzer L, Sironi L, Mainetti M, Ostuni R, Larimore K, Greenberg PD, de la Torre JC, Guidotti LG, Iannacone M (2016) Inflammatory monocytes hinder antiviral B cell responses. Sci Immunol, 1:

Schijns VECJ, Haagmans BL, Wierda CMH, Kruithof B, Heijnen IAFM, Alber G, Horzinek MC (1998) Mice lacking IL-12 develop polarized Th1 cells during viral infection. The Journal of Immunology, 160: 3958–3964

Schoenborn JR, Wilson CB (2007) Regulation of interferon-gamma during innate and adaptive immune responses. Adv Immunol, 96: 41–101

Schulz EG, Mariani L, Radbruch A, Höfer T (2009) Sequential polarization and imprinting of type 1 T helper lymphocytes by interferon-gamma and interleukin-12. Immunity, 30: 673–683

Sercan O, Stoycheva D, Hämmerling GJ, Arnold B, Schüler T (2010) IFN-gamma receptor signaling regulates memory CD8+ T cell differentiation. J Immunol, 184: 2855–2862

Sheikh AA, Groom JR (2020) Transcription tipping points for T follicular helper cell and T-helper 1 cell fate commitment. Cell Mol Immunol,

Snell LM, Osokine I, Yamada DH, De la Fuente JR, Elsaesser HJ, Brooks DG (2016) Overcoming CD4 Th1 Cell Fate Restrictions to Sustain Antiviral CD8 T Cells and Control Persistent Virus Infection. Cell Rep, 16: 3286–3296

Stone SL, Peel JN, Scharer CD, Risley CA, Chisolm DA, Schultz MD, Yu B, Ballesteros-Tato A, Wojciechowski W, Mousseau B, Misra RS, Hanidu A, Jiang H, Qi Z, Boss JM, Randall TD, Brodeur SR, Goldrath AW, Weinmann AS, Rosenberg AF, Lund FE (2019) T-bet Transcription Factor Promotes Antibody-Secreting Cell Differentiation by Limiting the Inflammatory Effects of IFN-γ on B Cells. Immunity, 50: 1172–1187.e7

Stuart T, Butler A, Hoffman P, Hafemeister C, Papalexi E, Mauck WM, Hao Y, Stoeckius M, Smibert P, Satija R (2019) Comprehensive Integration of Single-Cell Data. Cell, 177: 1888–1902.e21

Szabo SJ, Kim ST, Costa GL, Zhang X, Fathman CG, Glimcher LH (2000) A novel transcription factor, T-bet, directs Th1 lineage commitment. Cell, 100: 655–669

Tuzlak S, Dejean AS, Iannacone M, Quintana FJ, Waisman A, Ginhoux F, Korn T, Becher B (2021) Repositioning TH cell polarization from single cytokines to complex help. Nat Immunol, 22: 1210– 1217

Unger S, Seidl M, van Schouwenburg P, Rakhmanov M, Bulashevska A, Frede N, Grimbacher B, Pfeiffer J, Schrenk K, Munoz L, Hanitsch L, Stumpf I, Kaiser F, Hausmann O, Kollert F, Goldacker S, van der Burg M, Keller B, Warnatz K (2018) The TH1 phenotype of follicular helper T cells indicates an IFN-γ-associated immune dysregulation in patients with CD21low common variable immunodeficiency. J Allergy Clin Immunol, 141: 730–740

Vinuesa CG, Linterman MA, Yu D, MacLennan IC (2016) Follicular Helper T Cells. Annu Rev Immunol, 34: 335–368

Wakil AE, Wang ZE, Ryan JC, Fowell DJ, Locksley RM (1998) Interferon gamma derived from CD4(+) T cells is sufficient to mediate T helper cell type 1 development. J Exp Med, 188: 1651–1656

Walsh KP, Mills KH (2013) Dendritic cells and other innate determinants of T helper cell polarisation. Trends Immunol, 34: 521–530

Weinstein JS, Laidlaw BJ, Lu Y, Wang JK, Schulz VP, Li N, Herman EI, Kaech SM, Gallagher PG, Craft J (2018) STAT4 and T-bet control follicular helper T cell development in viral infections. J Exp Med, 215: 337–355

Whitmire JK, Benning N, Whitton JL (2005) Cutting edge: early IFN-gamma signaling directly enhances primary antiviral CD4+ T cell responses. J Immunol, 175: 5624–5628

Wong LR, Zheng J, Wilhelmsen K, Li K, Ortiz ME, Schnicker NJ, Thurman A, Pezzulo AA, Szachowicz PJ, Li P, Pan R, Klumpp K, Aswad F, Rebo J, Narumiya S, Murakami M, Zuniga S, Sola I, Enjuanes L, Meyerholz DK, Fortney K, McCray PB, Perlman S (2022) Eicosanoid signalling blockade protects middle-aged mice from severe COVID-19. Nature, 605: 146–151

Xia Y, Sandor K, Pai JA, Daniel B, Raju S, Wu R, Hsiung S, Qi Y, Yangdon T, Okamoto M, Chou C, Hiam-Galvez KJ, Schreiber RD, Murphy KM, Satpathy AT, Egawa T (2022) BCL6-dependent TCF-1^+^ progenitor cells maintain effector and helper CD4^+^ T cell responses to persistent antigen. Immunity, 55: 1200–1215.e6

Xu L, Cao Y, Xie Z, Huang Q, Bai Q, Yang X, He R, Hao Y, Wang H, Zhao T (2015) The transcription factor TCF-1 initiates the differentiation of TFH cells during acute viral infection. Nature immunology, 16: 991–999

Zhang Y, Apilado R, Coleman J, Ben-Sasson S, Tsang S, Hu-Li J, Paul WE, Huang H (2001) Interferon gamma stabilizes the T helper cell type 1 phenotype. J Exp Med, 194: 165–172

Zhu J, Yamane H, Paul WE (2010) Differentiation of effector CD4 T cell populations (*). Annu Rev Immunol, 28: 445–489

Zumaquero E, Stone SL, Scharer CD, Jenks SA, Nellore A, Mousseau B, Rosal-Vela A, Botta D, Bradley JE, Wojciechowski W, Ptacek T, Danila MI, Edberg JC, Bridges SL, Kimberly RP, Chatham WW, Schoeb TR, Rosenberg AF, Boss JM, Sanz I, Lund FE (2019) IFNγ induces epigenetic programming of human T-bet^hi^ B cells and promotes TLR7/8 and IL-21 induced differentiation. Elife, 8: e41641

